# Biosynthetic diversification of peptaibol mediates fungus-mycohost interactions

**DOI:** 10.1101/2022.06.05.494846

**Authors:** Jie Fan, Jinwei Ren, Ruolin He, Peng-Lin Wei, Yuanyuan Li, Wei Li, Dawei Chen, Irina S. Druzhinina, Zhiyuan Li, Wen-Bing Yin

**Affiliations:** State Key Laboratory of Mycology, Institute of Microbiology, Chinese Academy of Sciences, Beijing 100101, PR China; Center for Quantitative Biology, Peking University, Beijing 100871, PR China; Savaid Medical School, University of Chinese Academy of Sciences, Beijing 100049, PR China; Fungal Genomics Laboratory (FungiG), Nanjing Agricultural University, Nanjing 210095, PR China; NHC Key Laboratory of Food Safety Risk Assessment, Chinese Academy of Medical Science Research Unit (2019RU014), China National Center for Food Safety Risk Assessment, Beijing 100021, PR China

## Abstract

Fungi have evolved a plethora of functionally diverse secondary metabolites (SMs) to enhance their adaptation to various environments. To understand how structurally diverse metabolites contribute to fungal adaptation, we elucidate fungus-mycohost specific interactions mediated by a family of polypeptides, *i.e.*, peptaibols. We specified that peptaibol structural diversification was attributed to the nonspecific substrate recognition by the highly conserved peptaibol synthetases (PSs) in dead wood inhabiting mycoparasitic fungi from the genus *Trichoderma*. Exemplified by investigation of *T. hypoxylon*, we characterized a library of 19 amino acid residue peptaibols, named trichohypolins, containing 42 derivatives synthesized by a single PS enzyme (NPS1_Th_). Elimination of trichohypolin production by the deletion of *nps1*_Th_ reduced the inhibitory activities of *T. hypoxylon* on at least 15 saprotrophic host fungi, indicating that peptaibols are essential for interactions of *Trichoderma* spp. with their mycohosts. Different antagonistic effects of five trichohypolin subfractions SF1–SF5 and two pure compounds trichohypolins A (**1**) and B (**2**) on saprotrophic host fungi revealed specific activities of peptaibol derivatives in mediating fungus-mycohost interaction. Our study provides insights into the role of metabolic diversity of biosynthetic pathways in interfungal interactions.

## Introduction

Fungi are one of the most diverse groups of eukaryotes conquering nearly all ecosystems, that can only be compared to insects^1-4^. It means that *in vivo* each fungus inevitably becomes a member of the dense and complex network of interfungal interactions. With only a few exceptions, the majority of interfungal interactions are adverse as fungi are either competitive with one another or combative^5^. The absorptive nutrition of fungi constrains the shapes of their bodies and does not allow them to form defensive or high jacking structures that are present in higher eukaryotes. Therefore, the majority of fungal warfares are won through chemical weapons, which are shown as structurally diverse secondary metabolites (SMs)^1,6^.

Likely due to this reason, and because of the similar interactions with the organisms from the other domains of life, a considerable part of fungal genomic functionality is occupied by the biosynthetic gene clusters (BGCs)^7^. Certain genes arranged in a BGC are commonly involved in the enzymatic assembly of a predominant SM and several biosynthetic intermediates. In several cases, a single enzyme can also catalyze the formation of structurally diverse metabolites, such as a unique group of nonribosomal peptide synthetase (NRPS), also named peptaibol synthetase (PS), being responsible for a set of peptaibol products^8, 9^. Peptaibols are structurally featured with a high content of non-proteinogenic amino acids such as α-amino-isobutyric acid (Aib, U) and isovaline (Iva), an *N*-terminal acetyl group, and a *C*-terminal amino alcohol^10, 11^. The first and the most extensively studied peptaibol is the 20-residue alamethicin, of which different analogs exhibit broad-spectrum biological activities against bacteria, fungi, plants, and animals^12-14^. Previous studies have shown that peptaibols are predominantly produced by *Trichoderma* genus (Hypocreales, Ascomycota), which is the largest taxon among mycoparasitic fungal genera with many ubiquitously distributed species^10,15,16^. Generally, a single *Trichoderma* species possesses tremendous capacities for producing diverse peptaibols^8, 9^. For instance, 24 and 41 peptaibol derivatives were identified from *T. pleuroti* TPhu1^17^ and *T. reesei* QM6a^18^, respectively. Such abundant peptaibol production in *Trichoderma* spp. suggests peptaibols as the efficient metabolites for the versatile interactions of *Trichoderma* with their mycohosts and the competitive ecological niches^15, 19^.

With respect to peptaibol biosynthesis, a PS enzyme consists of multiple modules that act like an assembly line for peptaibol biosynthesis, each module including adenylation (A), thiolation (T), as well as condensation (C) domains and incorporating one amino acid substrate into the peptide. Only two PSs, TEX1 and TEX2, are revealed in the genome of *T. virens* Gv29-8, but involved in the biosynthesis of a large variety of peptaibols. TEX1 catalyzes the synthesis of 18-residue peptaibols belonging to trichorzin TVB group^8, 20^, and TEX2 catalyzes the synthesis of 88 peptaibols, 35 of which contain 11 residues and 53 of which contain 14 residues^9^. Given the central role of the NRPS A domains in substrate recognition and activation^21^, substrate specificities by A domains in PS enzymes may determine the peptaibol product diversification. Thus, we hypothesize that structural diversification of peptaibols can be explained by the relatedness of PS A domains with the substrate selectivity.

In this study, we first carried out genome mining throughout the *Trichoderma* genus for characterizing more PS genes and understanding their catalytic specificities in peptaibol formation. A 19-module PS enzyme NPS1_Th_ from *T. hypoxylon* was identified for the generation of a peptaibol library with 19-amino acid residues containing 42 trichohypolin derivatives, which were further revealed as essential metabolites for fungus-mycohost interaction. Precise evaluation of the identified metabolites/fractions from the peptaibol library demonstrated their differentiated functions when the peptaibol producing fungus was confronted with a wide range of mycohosts in nature.

## Results

### Peptaibol synthetase is highly conserved for peptaibol synthesis

A total number of 45 PS-encoding genes comprising over 10 modules were predicted in 30 draft genomes of *Trichoderma* spp. (Supplementary Table S3). Each strain showed the presence of one or two PSs, totally 25 long PSs comprising 17–20 modules and 20 short PSs comprising 10–16 modules, indicating that PS-encoding genes are highly conserved throughout *Trichoderma* genus. For example, NPS1Tr from *T. reesei* with 18 modules and NPS2_Tr_ with 14 modules were observed in the most commonly occurring species *T. reesei* belonging to section *Longibrachiatum*^22^. However, these enzymes were only described in rare occasions for peptaibol biosynthesis. Aforementioned 18-module TEX1 and 14-module TEX2 from *T. virens* in *Harzianum*/*Virens* clades have been proven for catalyzing a large number of peptaibol products containing 18, 14, or 11 residues^8, 9^. The 19-module PBS1, together with two 18-module PSs, *i.e.*, NPS1_Ta_ and NPS1_Tp_, were also previously deduced from the genome sequence of *T. atroviride*, *T. aggressivum*, and *T. pleuroti*^17, 23^ and connected to peptaibol formation in the respective species due to the same counts of PS modules with peptaibol residues.

In order to understand how a single PS enzyme produces a large variety of peptaibols, we used computational analysis to specify the catalytic mechanism. Substrate-related inter-motif sequences (termed A3–A6 intermotifs) of A domains, which overlap the substrate’s binding pocket, were extracted from PS enzymes referring to the NRPS “standardization” for sequence distance analysis^24, 25^. As presented in Fig. 1, hierarchical clustering of A domain distance matrix resulted in five separated clades. Combination of the curated TEX1, TEX2, NPS1_Tp_, NPS1_Ta_, and PBS1 with their peptaibol products prompted us to characterize the substrate specificities of A domains in peptaibols^8,9,17,23^. We found that clustering of A domains depends on their substrate specificities. 78 out of the 106 A domains fall into the largest clade, revealing high sequence similarity within these A domains (blue block in the upper-left area in Fig. 1). They showed selective preference for Aib (U), which is the characteristic substrate for peptaibol formation and occupies a high portion among all substrates^26^. However, Aib (U) is not the only type of substrates falling into the large clade. Also locating in the same clade are A domains that selects Val (V), Iva, Leu (L), Ala (A), Ser (S), or Gly (G). Interestingly, interchanges within U, A, Val/Iva (V_x_), Leu/Ile (L_x_), S, and G occurred in this clade, indicating variable substrate selectivities of the corresponding A domains. This large clade was further observed in the extensive clustering for A3–A6 intermotifs of 513 A domain sequences from 21 long PSs and 9 short PSs, also implying a relatedness of A domains in their substrate selections (Extended Data Fig. 1). Therefore, such a clade with A domains similar in sequence but different in selectivity is a strong indicator of substrate promiscuity and peptaibol product diversity (Fig. 1).

**Fig. 1.**
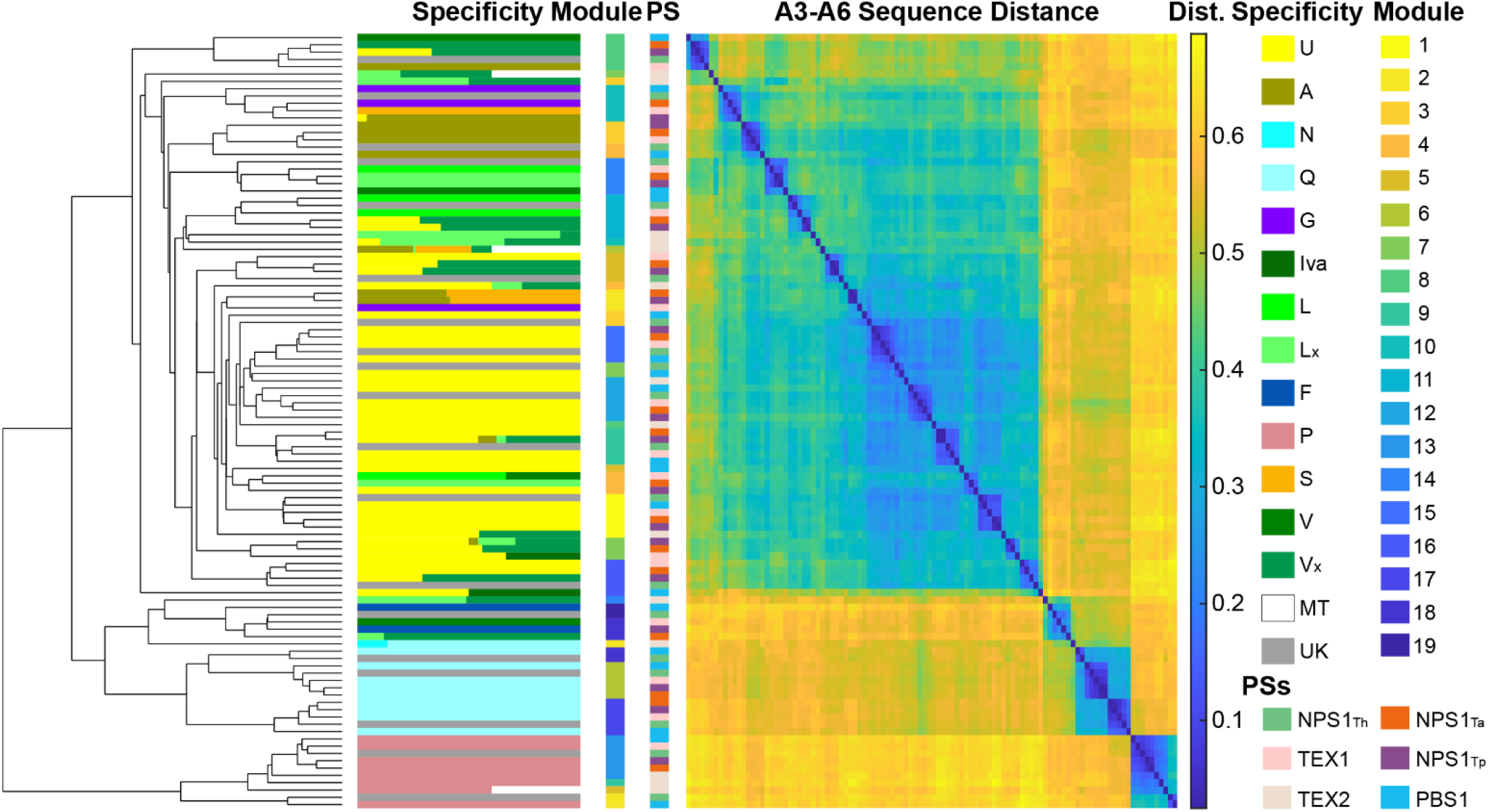
Structural diversity of peptaibols was revealed by clustering of the specificity-conferring region of A domains in peptaibol synthetases. 106 A domain sequences from an unknown peptaibol synthetase (PS) NPS1_Th_ in *T. hypoxylon* and five known PSs including TEX1 and TEX2 from *T. virens*, NPS1_Ta_ from *T. aggressivum*, NPS1_Tp_ from *T. pleuroti*, and PBS1 from *T. atroviride* are clustered as a phylogenetic tree on the left. Modules in PSs and their substrate specificities are attached to the multiple sequence alignment (the 2^nd^ and 3^rd^ columns from left). Distance matrix of A domain sequences is sorted using motif A3–A6 alignment, which overlaps the substrate’s binding pocket^24^. Substrates, PSs, and their containing modules are shown by colors in side bars. Substrates with similar chemical properties are represented by similar colors. U, Aib; L_x_, L or I; V_x_, V or Iva; MT, empty, that means this module could be skipped in the product synthesis; UK, unknown, that means the corresponding products of PS remain unknown. Amino acid sequences of PSs were obtained from the GenBank database. See more unknown PSs in Extended Data Fig. 1.

Other smaller clades are more specific in substrate selectivity. A domains selecting Gln (Q) residues cluster together, occasionally replaced by Glu (E). The residue Pro (P) was also specified to be extremely conserved and combined as Aib–Pro (U-P). The first A domain at *N*-terminus preferred to activate U for initial peptide assembly, undergoing an acetylation by the keto synthase (KS) and acyltransferase (AT) domains of PSs. Despite that most of the A domains exhibit clustering by their substrate selectivities, the last A domain at *C*-terminus was connected to variable residues, such as V, L, or Phe (F). This might be attributed to the substrate preference of reductase (R) domain but not only to the attached A domain.

With the above characterized association of A domains with their substrate specificities, peptaibol products can be predicted with more confidence using PS-encoding genes comparing with the existing prediction algorithms, that do not perform well in predicting non-proteinogenic amino acids^27^. We predicted the corresponding products of PS genes as displayed in the Dataset 1. According to our prediction, each PS gene is capable of generating diverse peptaibols with the same counts of residues but several variable positions. Interchanges among U, A, V_x_, L_x_, S, and G could occur at certain residues. Therefore, clustering of A domains helped us to understand the selection specificity and flexibility of A domains for activating amino acids, which were further combined to form structurally diverse peptaibol products.

### The structural diversity of peptaibols is determined by the peptaibol synthetase

To validate the product diversification by PSs, we selected a mycoparasite *T. hypoxylon* CGMCC 3.17906 where there was neither PSs nor peptaibols reported yet^28^. The putative 19-module NPS1_Th_ was encoded by a large 66.37 kb DNA region lacking introns and shared amino acid identities of 72.8% with the known 18-module TEX1 from *T. virens* and 76.3% with the putative 19-module NPS1_Tga_ from *T. gamsii* (Extended Data Fig. 2 and Supplementary Table S4). Deletion of the entire *nps1*Th in *T. hypoxylon* wild type (WT_Th_) and LC-MS analysis revealed complete disappearance of two dominant peaks, indicating the involvement of NPS1_Th_ in the formation of **1** and **2** (Fig. 2a).

**Fig. 2.**
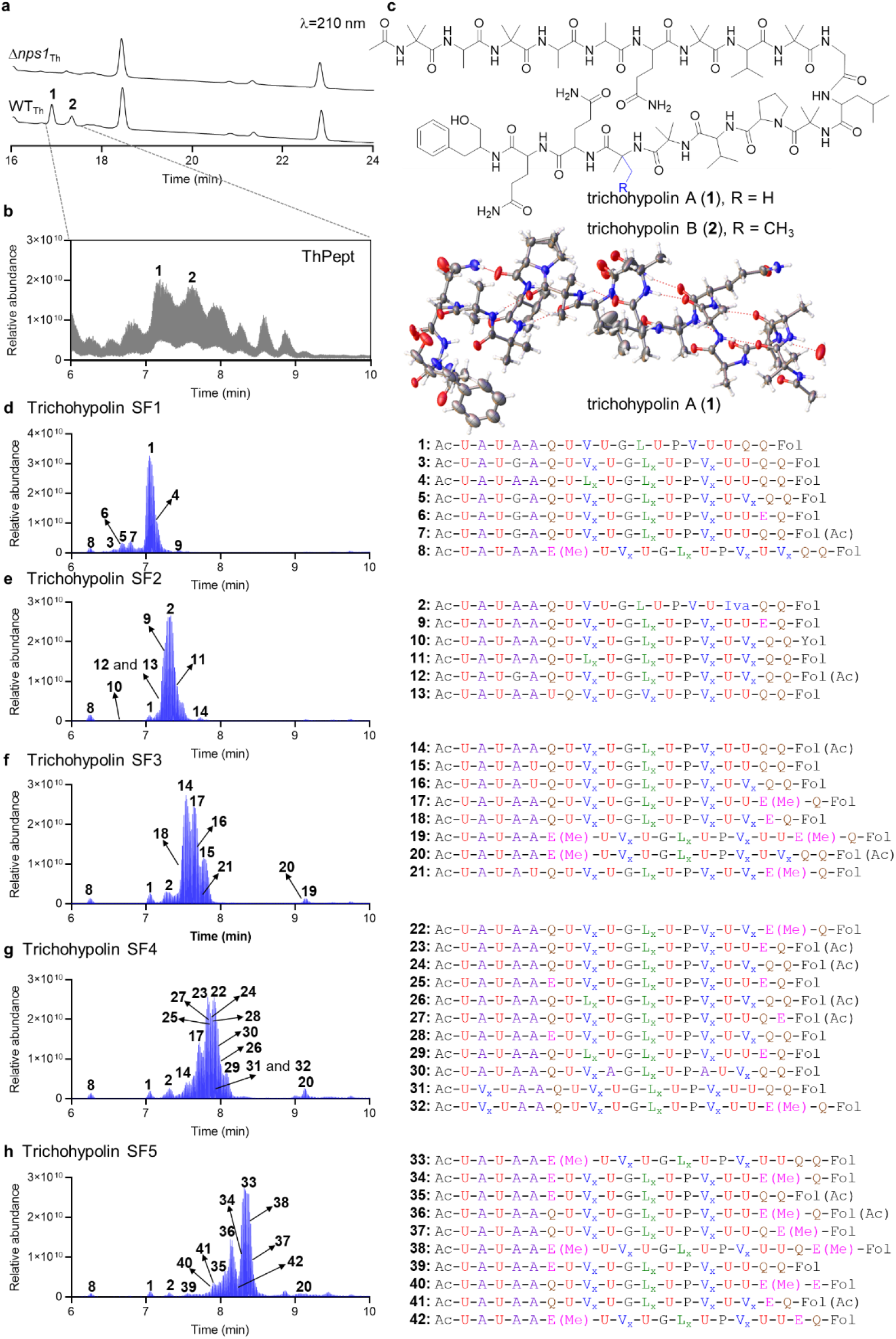
A peptaibol synthetase NPS1Th catalyzes the biosynthesis of 42 trichohypolin derivatives in *T. hypoxylon*. **a,** Analysis of metabolites from *T. hypoxylon* wild type (WT_Th_) and Δ*nps1*_Th_ by LC-MS. Peaks with retention time ranging from 16.6 to 17.6 min of WT_Th_ correspond to the products of NPS1_Th_, named trichohypolins. Trichohypolins A (**1**) and B (**2**) were shown as the predominant products. UV absorptions are illustrated at 210 nm. **b,** ESI-HRMS analysis of metabolites in a chemical mixture (ThPept) from of WTTh corresponding to the peaks with retention time arranging from 16.6 to 17.6 min on LC-MS in **a**. **c,** Structures of **1** and **2** with 19 residues identified by NMR and X-ray diffraction. **d**–**h**, 42 trichohypolin derivatives containing the predominant **1** and **2** detected in trichohypolin subfractions SF1–SF5. ESI-HRMS chromatograms of SF1–SF5 on the left correspond trichohypolins **1**–**42** in tabulated form on the right. Main components of each subfraction: SF1, **1** and **3**–**8**; SF2, **2** and **9**–**13**; SF3, **14**–**21**; SF4, **22**–**32**; and SF5, **33**–**42**. Residues in trichohypolin sequences are shown using the one-letter code. U, Aib; L_x_, L or I; V_x_, V or Iva; Ac, acetyl group; Fol, phenylalaninol; E(Me), methylated glutamate.

To identify the abolished products, large-scale cultivation of WT_Th_ was carried out and led to a separated chemical mixture (ThPept) corresponding to the peaks observed from 16.6 to 17.6 min on LC-MS (Fig. 2b). ESI-HRMS analysis of the mixture revealed dozens of compounds with **1** and **2** as the most abundant components. Subsequently, **1** and **2** were purified and characterized by ESI-HRMS and NMR analysis including ^1^H, ^13^C, ^1^H-^1^H COSY, HSQC, HMBC, ROESY, TOCSY, and HSQC_TOCSY (Supplementary Fig. S3–S18 and Table S5). Diagnostic fragmentation indicated the presence of b_12_ ^+^ and y_7_ ^+^ ions in the respective structures (*m/z* 1078.6274 and 774.4531 for **1**, *m/z* 1078.6267 and 788.4682 for **2**), which are characteristic fragment ions being indicative for a U–P bond of 19-residue peptaibol (Extended Data Fig. 3 and Fig. 4)^29^. Seven U residues, three A residues, three Q residues, two V_x_ residues, one G residue, one L_x_ residue, one P residue, and one phenylalaninol (Pheol, Fol) were observed by stepwise fragmentation of **1**. Combined with the detailed NMR data, **1**, named trichohypolin A, was thus assigned as Ac–U^1^–A^2^–U^3^–A^4^–A^5^–Q^6^–U^7^–V^8^–U^9^–G^10^–L^11^– U^12^–P^13^–V^14^–U^15^–U^16^–Q^17^–Q^18^–Fol^19^, which was confirmed by X-ray analysis (Fig. 2c). In comparison with **1**, the MS^2^ analysis of the y_7_ ^+^ ion demonstrated a V_x_ residue at position 16 in trichohypolin B (**2**) instead of a U residue in **1**, which was proven to be Iva by NMR analysis (Fig. 2c and Extended Data Fig. 3).

**Fig. 3.**
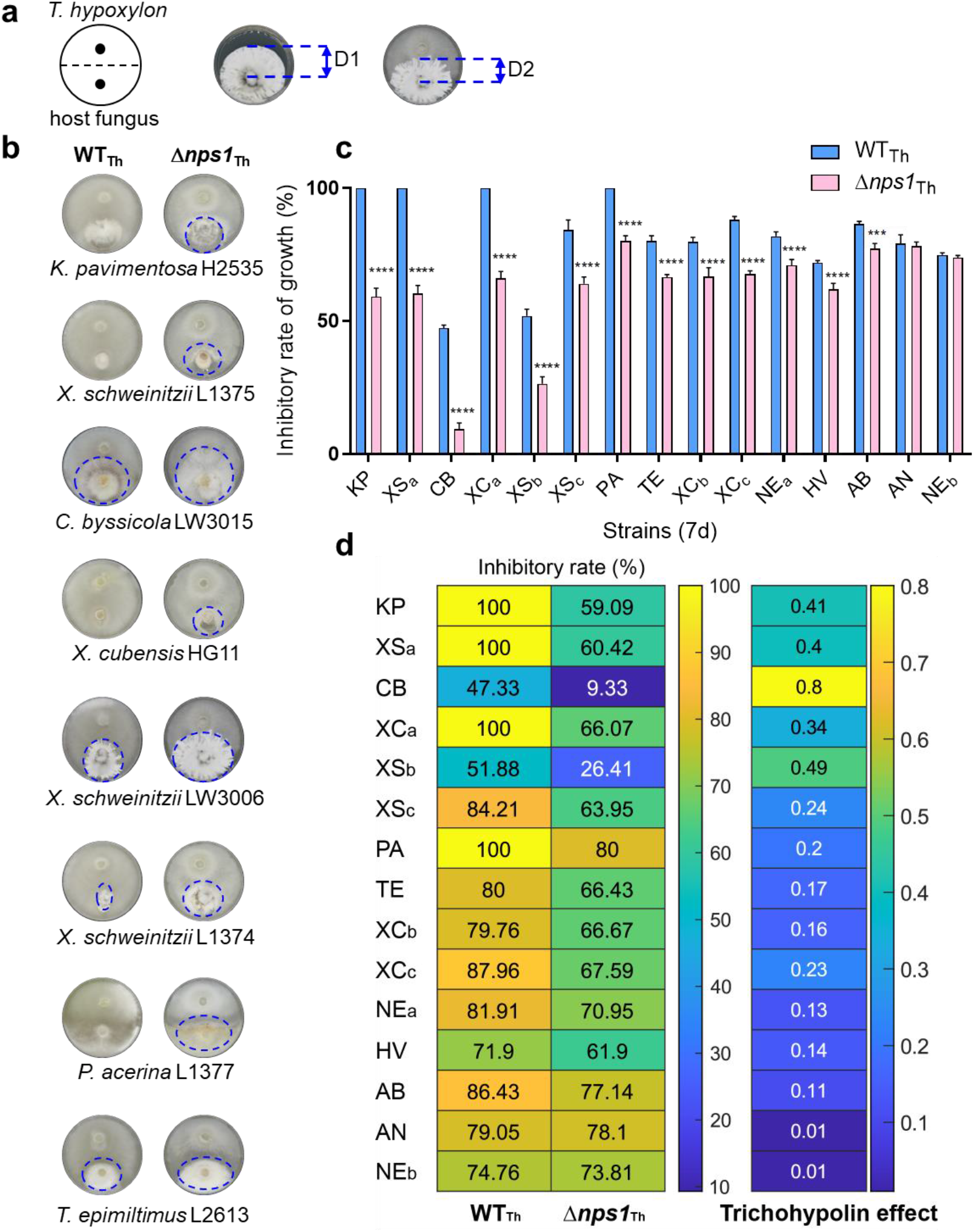
Elimination of peptaibol production reduces antagonistic effects of a *Trichoderma* spp. on its mycohosts. **a**, Schematic of experimental setup and calculation method for dual confrontation assays of *T. hypoxylon* (above) with a saprotrophic host fungus (below). D1 and D2 represent the growth radiuses of *T. hypoxylon* strain and the mycohost, respectively. **b,** Dual confrontation assays of *T. hypoxylon* wild type (WT_Th_) and Δ*nps1*_Th_ with eight representative host fungi. Culture front of plates are shown, and the detailed mophologies are displayed in Extended Data Fig. 8. Blue circles show the extension of fungal colonies. **c** and **d,** Antagonistic potential of *T. hypoxlon* strains against host fungi. For **c,** the shadowing column indicates WTTh (light blue) or Δ*nps1*Th (pink) capable of producing a diffusible inhibitor against host fungi. Bars represent standard errors of the mean for three replicates and asterisks indicate the significant differences according to two-way ANOVA test: \*\*\**P* < 0.001, \*\**P* < 0.01, \**P* < 0.05. For **d,** the colored patches indicate growth inhibitory of a fungus by *T. hypoxylon*, with yellow represents high inhibitory effect. The antagonistic effect of trichohypolins, which are formed by NPS1_Th_ catalysis, towards mycohosts is the inhibitory changes when *nps1*_Th_ was deleted in *T. hypoxylon* (see Method), with yellow represents high effect level. KP, *K. pavimentosa* H2535; XS_a_, *X. schweinitzii* L1375; XS_b_, *X. schweinitzii* LW3006; XS_c_, *X. schweinitzii* L1374; CB, *C. byssicola* LW3015; XC_a_, *X. cubensis* HG11; XC_b_, *X. cubensis* L1345; XC_c_, *X. cubensis* HG19; PA, *P. acerina* L1377; TE, *T. epimiltinus* L2613; NE_a_, *N. ellipsospora* LW3103; NE_b_, *N. ellipsospora* LW2361; HV, *H. vinosopulvinatum* LW2881; AB, *A. bovei* LW2607; AN, *A. nitens* LW3091.

**Fig. 4.**
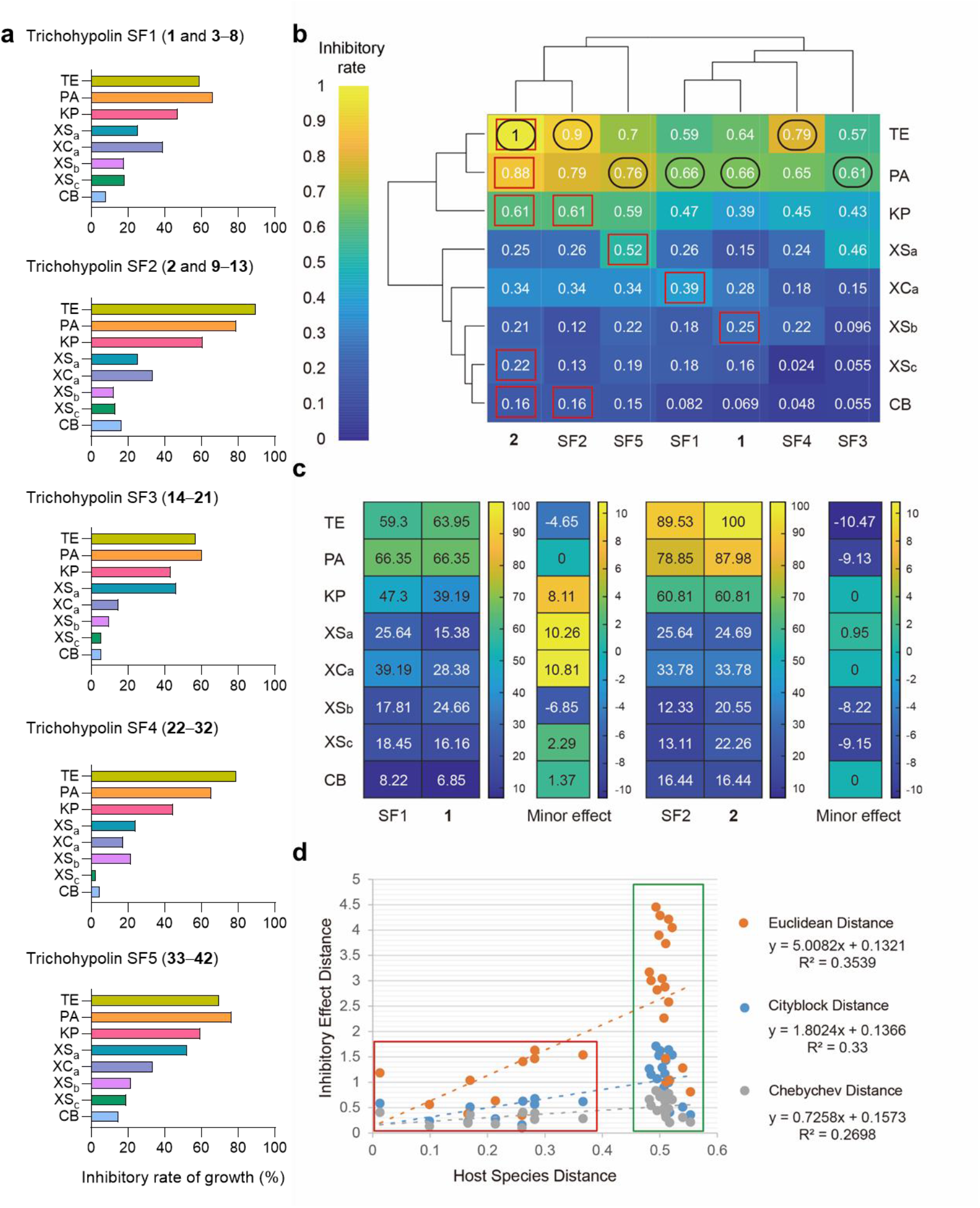
Specific antagonism of peptaibol derivatives towards saprotrophic host fungi. **a,** Antagonistic potential of peptabol (trichohypolin from *T. hypoxylon*) subfractions (SF1–SF5) against the mycohosts of *T. hypoxylon*. The shadowing columns in different colors represent growth inhibitory rates of host fungi. Main components in trichohypolin subfractions are trichohypolins A (**1**) and **3**–**8** in SF1, trichohypolins B (**2**) and **9**–**13** in SF2, **14**–**21** in SF3, **22**–**32** in SF4, and **33**–**42** in SF5. The growth inhibitory rates are calculated as seen in method. **b,** Inhibitory effects of the pure compounds **1** and **2**, as well as SF1–SF5 on host fungi in tabulated form. The colored patches indicate growth inhibitory of host fungi (in different rows) by trichohypolins (in different columns), with yellow represents high inhibitory effect. Patches in black circles represent the most sensitive saprotrophic strains antagonized by a trichohypolin subfraction or compound. Patches in red boxes represent trichohypolin subfraction or compound with the highest inhibitory effect on a host fungus. Both rows and columns are clustered to present fungal distance and inhibitory effect distance. Fungal distance was acquired by pairwise *p*-distance analysis of *LSU* nucleic acid sequences from different saprotrophic fungi. **c,** Effects of the minor components in SF1 (**3**–**8**) and SF2 (**9**–**13**) on host fungi. Yellow represents high effects of trichohypolin compounds. **d,** Relationship of inhibitory effect of trichohypolins with mycohost distance. Three special cases of Minkowski distance are carried out to calculate inhibitory effect distance, leading to Euclidean distance (organge), City block distance (blue), and Chebychev distance (grey). KP, *K. pavimentosa* H2535; XS_a_, *X. schweinitzii* L1375; XS_b_, *X. schweinitzii* LW3006; XS_c_, *X. schweinitzii* L1374; CB, *C. byssicola* LW3015; XC_a_, *X. cubensis* HG11; PA, *P. acerina* L1377; TE, *T. epimiltinus* L2613.

The ThPept mixture was also separated into five subfractions, *i.e.* trichohypolins SF1–SF5, as the elution order on semi-preparative HPLC (Fig. 2d–h). Referring to the precise structures of **1** and **2**, at least 40 more trichohypolin derivatives were recognized in SF1–SF5 by diagnostic fragmentation analysis (Supplementary Table S6 and Fig. S19–S58). Trichohypolins SF1 and SF2 mainly contain seven and six trichohypolin derivatives, of which **1** and **2** were shown with the highest portions, respectively (Fig. 2d,e). Eight trichohypolin derivatives, **14**–**21**, were identified in trichohypolin SF3. Trichohypolin SF4 possesses eleven derivatives with **22**–**24** as the most abundant molecules. Three out of ten derivatives, **33**, **34**, and **36**, were observed as predominant molecules in SF5 (**33**–**42**). All trichohypolin derivatives contain 19 amino acid residues but with variable ones at several positions. Sequence comparison reveals that seven residues, including residues U^1/3/9/10/12/15^ and P^13^, are conserved in all trichohypolin derivatives (Fig. 2 and Extended Data Fig. 5). Each compound comprises 26%–42% U and an extremely conserved U^12^–P^13^ bond, which was essential for the biological activity of peptaibols since U–P bond can break the helical structure^30^. Interchange between U and V_x_ at residue 16 is the most frequently found among all trichohypolins. The variable residues Q^6/17/18^/E^6/17/18^, A^2^/V_x_ ^2^, A^4^/G^4^, A^5^/U^5^, U^7^/Q^7^, V_x_ ^8/11^/L_x_ ^8/11^, as well as V_x_14/A^14^ are also observed. Modifications, such as methylation of E and acetylation of Fol, occur occasionally at certain positions. Thus, mapping of the substrate selectivities to the 19 A domains confirmed the substrate specificities and flexibilities, which led to trichohypolin product diversification (Extended Data Fig. 5). This corresponded well with the peptaibol prediction using the relatedness of A domains with their substrate selections.

### Peptaibols contribute to *Trichoderma* interactions with a wide range of mycohosts

The versatile mycoparasitism regarding interspecies interactions were historically studied in the environmental opportunistic species from the genus *Trichoderma* based on a few models and frequently soil-born plant pathogenic fungi^16,31,32^. However, the diversity of *Trichoderma* in soil is minor compared to it above ground^33, 34^. Therefore, to determine if the abundant peptaibol production in *Trichoderma* spp. is related to the fungus-mycohost interaction in the native environment of *Trichoderma*, we proceeded to carry out dual confrontation assays of *T. hypoxylon* with saprotrophic host fungi to mimic the ecological conditions (Extended Data Fig.6 and Supplementary Table S1). Considering the parasitism of *T. hypoxylon* against *Hypoxylon* (Xylariales, Ascomycota) species in the ecosystem^28^, a library of 15 saprotrophic fungi were identified for investigation of fungus-mycohost interaction (Extended Data Fig.8).

They are including three *Xylaria schweinitzii* strains (XS_a_, XS_b_, and XS_c_), three *Xylaria cubensis* strains (XC_a_, XC_b_, and XC_c_), two *Neopestalotiopsis ellipsospora* strains (NE_a_ and NE_b_), *Hypoxylon vinosopulvinatum* (HV), *Annulohypoxylon bovei* (AB), *Annulohypoxylon nitens* (AN), *Kretzschmaria pavimentosa* (KP), *Phlebia acerina* (PA), *Clonostachys byssicola* (CB), and *Tinctoporellus epimiltinus* (TE) (Extended Data Fig. 7).

As expected, the growth of all 15 fungi were obviously reduced when they were cocultivated with *T. hypoxylon* WT_Th_, with the growth inhibition ranging from 47% to 100% (Fig. 3 and Extended Data Fig. 8). Growth of four strains XS_a_, XC_a_, PA, and KP was completely inhibited, while XC_c_ and AB were inhibited but not killed. Growth inhibitory of the screening fungi can be differentiated even in different strains of *X. schweinitzii*, *X. cubensis*, and *N. ellipsospora*. Within *X. schweinitzii*, less antagonistic effects on XSb and XS_c_ were observed than that on XS_a_, with growth inhibitory rates of 52% and 84%, respectively. The WT_Th_ strain showed the least antagonistic ability towards CB (47%). These results were consistent with the mycoparasitic lifestyle of *Trichoderma* spp. attacking wood-rot that made use of plant biomass pre-degraded by earlier colonizers^15^. Such antagonism of *T. hypoxylon* on a wide range of mycohosts confirmed the remarkable mycoparasitic vigor at both genus and species level, contributing to the ubiquitous distribution of *Trichoderma* spp. and their adaptation to diverse habitats.

By comparison, the deletion of the *nps1* gene reduced antagonistic abilities towards all 15 fungi, as presented in Fig. 3b–d. Inhibitory rates ranging from 9% to 80% were shown for 15 saprotrophic fungi by Δ*nps1*_Th_ strain, decreasing 0.95%–41% in comparison with WT_Th_. Among all 15 strains, PA showed the highest inhibitory rate (80%) by Δ*nps1*_Th_ comparing with that of 100% by WT_Th_. Antagonism against XS_a_, XC_a_, and KP, which were completely inhibited by WT_Th_, were decreased to 60%, 66%, and 59%, respectively. Despite that *T. hypoxylon* WT_Th_ showed relatively less antagonistic abilities towards CB (47%) and XS_b_ (52%) than that towards other saprotrophic fungi, even lower growth inhibitory rates (9% and 26% respectively) were observed for CB and XS_b_ by *nps1*_Th_ deletion mutant.

Declined growth inhibitory rates indicated that elimination of trichohypolin production by *nps1*_Th_ deletion led to a reduction of fungal antagonistic effects towards the mycohosts. Therefore, we calculated the antagonistic effect of peptaibol products, trichohypolins, by comparing growth inhibitory rates by Δ*nps1*_Th_ with those by WT_Th_ (Fig. 3d). A ranking of antagonistic effects by trichohypolins were shown for 15 fungi ranging from 0.01 to 0.8. The most antagonistic effect (0.8) by trichohypolins was observed for the strain CB, followed by XS_b_ with an antagonistic effect of 0.49. Relative higher antagonistic effects were also shown for KP and XS_a_ at 0.41 and 0.4, respectively. However, no obvious inhibition changes were detected for AN and AB. Thus, we concluded that the PS enzyme NPS1_Th_ and its products trichohypolins can be deduced as the genetic and metabolic effectors in the interactions of *T. hypoxylon* with its mycohosts, but showing differentiated activities.

### Structural diverse peptaibols differentially mediate the specific fungus-mycohost interactions

To address how 42 trichohypolin derivatives control the interactions of WT_Th_ with a wide range of mycohosts, we assessed the inhibitory activities of the two purified trichohypolins **1** and **2**, and trichohypolin subfractions (SF1–SF5). As presented in Extended Data Fig. 9, XS_a_, XS_b_, XS_c_, XC_a_, CB, PA, KP, and TE were selected for peptaibol activity assessment because of the relatively higher antagonistic effects on them by trichohypolins (Fig. 3d). By ranking the inhibitory rates, we found quite different activities of the seven examined trichohypolin chemicals (**1**, **2**, and SF1–SF5) towards different fungal strains (Supplementary Table S7).

Of all strains, the remarkable antagonism was observed for PA and TE by trichohypolin chemicals comparing with other saprotrophic fungi, with inhibitory ranges of 60%–88% and 56%–100%, respectively (patches with black boxes in Fig. 4b). For example, trichohypolins SF1, SF3, SF5, and **1** showed antagonistic effects on PA with inhibitory rates of 66.35%, 60.58%, 76.44, and 66.35%, respectively. As presented in the patches with red boxes (Fig. 4b), the highest antagonistic effects by trichohypolin chemicals were exhibited for each screening fungus. **2** was revealed to possess relatively higher and broader inhibitory effects than other trichohypolin chemicals, particularly with a complete antagonistic effect towards TE. Since **2** was a predominant component in trichohypolin SF2, SF2 also showed a higher inhibitory effect towards TE at 90%. With respect to the fungal strain KP, **2** and its corresponding subfraction SF2 were the most antagonistic molecules among the seven examined trichohypolin chemicals, both with the inhibitory effect of 61%. Comparing with other trichohypolin chemicals, trichohypolins SF1 and SF5 exhibited higher antagonistic effects on XC_a_ (39%) and XS_a_ (52%), respectively.

Furthermore, we calculated the activities of the corresponding minor components in SF1 and SF2 by comparing the antagonistic effects of purified trichohypolins (**1** and **2**) with those of SF1 and SF2 (Fig. 4c). Even though **1** occupied the highest ratio in SF1, antagonistic effects of **1** and SF1 were obviously differentiated towards fungal strains. For example, trichohypolin SF1 showed an antagonistic effect of 39.19% on an XC_a_, whereas a less antagonistic effect was observed for **1** (28.38%). This indicated that the minor components **3**–**8** in SF1 also act as key molecules in response to XC_a_. Similar impacts of **3**–**8** were further found to antagonize XS_a_ and KP (Fig. 4c). Differing from SF1, no obvious inhibitory effect was observed for the minor components in SF2 (**9**– **13**). The minor components (**3**–**13**) even function inversely to **1** and **2** in several cases, for instance, inhibitory effects of **9**–**13** in SF2 at -10.47 and -8.22 for TE and XS_b_, respectively.

Collectively, trichohypolin chemicals including two purified molecules **1** and **2**, as well as subfractions SF1–SF5 functioned with differential antagonistic effects on diverse fungal strains. It might occur like that 42 trichohypolin derivatives functioned cumulatively and cooperatively when the peptaibol producing *T. hypoxylon* was confronted with its host fungi. Considering the phylogenetic distance of fungal strains, we found three strains locating in the same clade, including TE, PA, and KP, with relatively higher antagonistic effects by each trichohypolin chemical than the strains in other clades (Fig. 4b). Calculation of inhibitory effect distance prompted us to cluster trichohypolin chemicals into two major groups. The correlated trichohypolins, *e.g.*, **2**, SF2, and SF5 locating in one group, plausibly functioned with similar activities towards the mycohosts. Three special cases of Minkowski distance calculation revealed that similar antagonistic effects by peptaibols might occur on the phylogenetically close fungi (Fig. 4d). These results indicated that diverse peptaibols functioned differentially in response to fungus-mycohost interactions. In turn, mycohosts can determine metabolic requirement of a fungal strain for environmental adaptation.

## Discussion

Fungal SMs are required for the ecological interactions of interspecies and the rapid adaptation to changing micro-environment^5^. For instance, aflatoxin production provides a fitness advantage to *Aspergillus flavus* when encountering insects, and anthraquinone-based pigmentation in *A. flavus* sclerotia promotes the resistance to both biotic and abiotic stressors^6,35,36^. Exemplified by the identification of a 19-residue peptaibol library containing 42 trichohypolins in *T. hypoxylon*, we found that only a single PS enzyme (*e.g.*, NPS1_Th_) can catalyze the production of abundant peptaibols as potential adaptive molecules for interfungal interactions (Fig. 2).

The amino acid sequences of peptaibols are commonly determined by ESI-HRMS, NMR spectroscopy, and X-ray analysis of the pure peptaibol compounds for complete elucidation and ESI-HRMS fragmentation diagnosis of fungal extracts for sequence speculation^26^. It is usually time-consuming and high-cost due to the majority of peptaibols with very low yields. Here we searched for a more efficient method for peptaibol structure prediction on basis of the knowledge for understanding peptaibol biosynthesis (Fig. 1 and Extended Data Fig. 1). PS enzymes, together with their A domains, were highly conserved throughout *Trichoderma* genus. Substrate specificities of A domains in PS enzymes were specified by 513 A domain sequence alignment referring to the previous identification of TEX1, TEX2, NPS1_Tp_, NPS1_Ta_, and PBS1^8,9,17,23^. Interchanges at several certain residues can be observed throughout the peptaibol sequences, particularly those occurring for selectivity of U/A/V_x_/L_x_/S/G. This well explained the peptaibol diversification by a single PS enzyme and was then validated by trichohypolin production by NPS1_Th_ in *T. hypoxylon*. Thus, PS-encoding gene mining and in-depth analysis of A domains can be further applied as a tool for more peptaibol discovery.

Considering the ancestral mycoparasitic lifestyle of *Trichoderma* spp.^19,22,37^, we mimicked the fungus-mycohost interaction by confrontation assays of *T. hypoxylon* with a library of saprotrophic fungi (Fig. 3 and Extended Data Fig. 8). Elimination of trichohypolin production by entire *nps1*_Th_ deletion led to a reduced antagonistic abilities of *T. hypoxylon* against all screening mycohosts (Fig. 3). Therefore, the paralogous PS-encoding gene can be revealed as a genetic effector and its peptaibol products as metabolic effectors for contributing to the interactions of *Trichoderma* spp. with a wide range of mycohosts. The highest antagonistic effects by trichohypolins were observed for the *Clonostachys* strain CB, followed by an *X. schweinitzii* strain XS_b_ and the *Kretzschmaria* strain KP (Fig. 3d). In contrast, no obvious antagonism was shown for the two *Annulohypoxylon* strains by trichohypolins. These results demonstrated that peptaibol products differentiate their activities onto different mycohosts. Aside from peptaibols, other SMs might also be deployed by *Trichoderma* spp. for fungus-mycohost interaction in synergism with cell wall-degrading enzymes, proteolytic, and reactive oxygen species^38-40^.

In analogy to the previously identified peptaibols in *Trichoderma* spp., trichohypolin structural diversification was also revealed among 42 19-amino acid residue derivatives^17,18,26^. Inhibitory effects of seven trichohypolin chemicals, including two purified ones **1** and **2**, as well as five subfractions SF1–SF5, were assessed for fungal strains (Fig. 4 and Extended Data Fig. 9). 42 trichohypolin derivatives possessed their own preferences for recognizing and antagonizing the exogenous fungi, which were cumulated to show the total antagonistic effects (Fig. 3d). The predominant component **1** and the minor ones **3**–**8** in SF1 were both revealed with antagonistic effects, but differentiated on the host fungi. SF2, particularly its predominant component **2**, showed relatively broad-spectrum and higher antagonistic effects. Relevance of fungal phylogeny and trichohypolin effects indicated similar antagonistic effects on saprotrophic hosts with close phylogenetic distances. Thus, we concluded that a mycoparasite *Trichoderma* spp. can produce abundant peptaibols by a single PS enzyme and assign them to function as active molecules to interact with specific mycohosts. Ongoing metabolic adaptations in fungi are required for further growth and maintenance within the ecosystem.

## Methods

### Fungal strain isolation, cultivation, and identification

The fungal strain used in this study are summarized in Supplementary Table S1. *Trichoderma hypoxylon* CGMCC 3.17906 was isolated from the stroma of *Hypoxylon anthochroum* in Thailand^28^. The wild type (WT_Th_) strain and deletion mutants were cultivated at 25°C on potato dextrose agar (PDA, BD Difco™) for growth and on rice medium for detection of secondary metabolites (SMs). *E. coli* DH5α was grown in LB medium (1% NaCl, 1% tryptone, and 0.5% yeast extract) for standard DNA manipulation. 100 μg/mL ampicillin were supplemented for cultivation of recombinant *E. coli* strains.

15 saprotrophic fungi were isolated from wood collected in different areas of China (Extended Data Fig. 6 and Supplementary Table S1). They were cultivated on PDA plates at 25°C for growth and in potato dextrose broth (BD Difco™) culture for genomic DNA extraction as previously reported^41^. Phylogenetic analyses were conducted based on *ITS*, *LSU*, *RPB2*, and *β-tubulin* as DNA barcode fragments for species identification (Extended Data Fig. 7). PCR amplification was carried out by using Phusion® High-Fidelity DNA polymerase from New England Biolabs (NEB) on a T100TM Thermal cycler from Bio-Rad and the primer pairs listed in Supplementary Table S2. PCR reaction mixtures and thermal profiles were carried out as recommended by the manufacturer’s instruction. The DNA barcode fragments were obtained after DNA sequencing (Sangon Biotech, Shanghai, China).

DNA sequences of reference strains and *Capsaspora owczarzaki* ATCC 30864 used as an outgroup were retrieved from the GenBank (https://www.ncbi.nlm.nih.gov/genbank/^42"^. Sequences alignment was performed with MAFFT v 7 (https://mafft.cbrc.jp/alignment/server/index.html) and was improved with MEGA 7.0. Phylogenetic sequence database was conducted for the concatenated ITS-LSU-RPB2-Tub dataset by SequenceMatrix 1.7.8. Phylogenetic reconstructions were made by maximum likelihood (ML) analyses with RAxML 8.2.10 with 1000 replicates2 and the phylogenetic tree was visualized by FigTree v 1.4.4.

### Genome mining of peptaibol synthetases and product prediction

To exploit more peptaibol synthetases (PSs), we searched for the available genomic sequences of *Trichoderma* genus in the national center for biotechnology information (NCBI) database and the JGI Fungal program. With the known 18-module TEX1^20^ and 14-module TEX2^9^ as queries, we mined 43 putative PS genes comprising >10 modules in the 30 draft genomes of *Trichoderma* spp. (Supplementary Table S3)^22^. Two 18-module NRPS proteins encoded by *nps1*Tp from *T. pleuroti* and *nps1*Ta from *T. aggressivum*^17^ and a 19-module NRPS encoded by *pbs1* from *T. atroviride*^23^ are capable of producing peptaibols detected in the extracts of corresponding strains. Other PS genes are still unknown or not assigned to the corresponding peptaibol products.

Using a previously reported method^25^, we first extracted motif and inter-motif sequence of A domain from 21 PSs containing 18–20 modules and 9 PSs containing 12–15 modules. Then we proceeded the multiple sequence alignment (MSA) for each motif and inter-motif by Cluster Omega 1.2.4^43^ and calculated the sequence distance matrix by *p*-distance with BLOSUM62 score matrix. Sequence distance matrix was clustered by hierarchical clustering. Substrate specificity by each module would be patched and the proportion of different substrates was determined by its proportion attached in the detected peptaibol products.

Finally, the predicted specificities of A domain with unknown substrates were inferred by A domains with known substrate. If there were more than one kind of known substrates, the cluster would be named by their frequency in descending order. If there were no known substrate, it would be named as “unknown”. Clusters with same substrate specificity but different cluster were distinguished by a different suffix number. Substrate specificity, module, and PS genes were labeled by colored side bars.

### Construction of deletion cassettes and genetic manipulation

The oligonucleotide sequences for PCR amplification were given in Supplementary Table S2. For creation of *nps1*_Th_ deletion strain, approximately 2.7 kb sequences located upstream and downstream of the target gene were amplified from genomic DNA of *T. hypoxylon* using designated primer pairs hypoPS-5F-F/R and hypoPS-3F-F/R (Supplementary Table S2). These two fragments were fused with a hygromycin (*hph*) resistance gene from pUCH2-8 vector using primer pairs hypoPS-nest-F/R to construct the deletion cassette using the double-joint method described previously^44^. PEG-mediated protoplast transformation was performed as previously described^45, 46^. Transformants of TYYL2 were verified using three pairs of designated primers hypoPS-RT-F/R, hypoPS-5F-F/hyp-scr-5F-R, as well as hypoPS-3F-R/hyp-scr-3F-F and selected twice using 60 μg/mL hygromycin to obtain mitotic stability.

### Large-scale fermentation and preparation of trichohypolins

To isolate **1** and **2**, *T. hypoxylon* WT_Th_ strain was cultivated in 100 x 500 mL flasks each containing 100 g rice and 120 mL H_2_O at 25°C for 40 days. The rice cultures were extracted repeatedly with 15 L ethyl acetate and concentrated under reduced pressure to obtain a crude extract (83.6 g). The crude extract was applied to Silica gel column chromatography eluted with a stepwise gradient dichloromethane-acetone (v/v, 1:0, 100:1, 80:1, 50:1, 10:1, 8:1, 4:1, 2:1, 1:1, 1:4, 1:20, 1:50), yielding 21 fractions (Fr.1–Fr.21). Fr.13 (10.6 g) was subjected to reversed-phase C18 silica column chromatography eluted with methanol–H_2_O (v/v, 20:80, 30:70, 40:60, 50:50, 60:40, 70:30, 90:10, 100:0) to give 18 fractions (Fr.13-1 to Fr.13-18). Further purification of Fr.13-13, also named ThPept, containing peptaibols was carried out by semi-preparative HPLC on an SSI system (Teledyne SSI Lab Alliance Series III pump system and Series 1500 Photodiode Array Detector) with an Eclipse XDB-C18 column (9.4 × 250 mm, 5 μm, Agilent). Acetonitrile (A) and water with 0.1% formic acid (B) were used as solvents at flow rate of 2 mL/min. The substances were eluted with 48% (v/v) solvent A for 60 min. Trichohypolin A (**1**) was eluted at 33 min (20 mg) and trichohypolin B (**2**) was eluted at 40.6 min (18 mg).

Trichohypolin subfractions (SF1–SF5) was also prepared from the trichohypolin containing fraction ThPept (150 mg) on the semi-preparative HPLC system as mentioned above. Acetonitrile (A) and water with 0.1% formic acid (B) were used as solvents at flow rate of 2 mL/min. The substances were eluted with 48% (v/v) solvent A for 42 min, followed by a linear gradient from 48% to 57% (v/v) solvent A for another 38 min, and 57% (v/v) solvent A for 15 min. The column was then washed with 100% (v/v) solvent A for 10 min and equilibrated with 48% (v/v) solvent A for 5 min. SF1 (34.3 mg) was eluted from 20 to 38 min, SF2 (18.7 mg) was eluted from 38 to 50 min, SF3 (6.0 mg) was eluted from 50 to 60 min, SF4 (3.3 mg) was eluted from 60-71 min, and SF5 (3.1 mg) was eluted from 70 to 83 min.

### Antagonism effects on saprotrophic fungi by *T. hypoxylon*

Confrontation assays of *T. hypoxylon* strains with 15 different saprotrophic fungi were carried out as described in previous studies^47, 48^. The mycelia of *T. hypoxylon* strains (diameter 5 mm) and a saprotrophic fungus (diameter 5 mm) were cut from the edge of an actively growing colony on 3-day-old or 5-day-old PDA plates. They were then placed on a 60 mm-diameter PDA plate at opposite sides with a distance of 30 mm from each other (Fig. 3A). All cocultivation plates were cultured at 25°C for 7 days in the darkness. Growth inhibitory rate of a saprotrophic fungus was calculated using the following formula: Growth inhibition (%) = (D1–D2)/D1 x 100%. D1 and D2 represent the growth radiuses of *T. hypoxylon* and the saprotrophic host fungus, respectively. Growth inhibitory rates were presented as mean ± standard deviation of the mean (SD). Significance of growth inhibition was tested by a two-way ANOVA using Graphpad Prism 8.0, and differences were significant when *P* < 0.05.

Antagonism effects of trichohypolins, which are biosynthesized by NPS1_Th_, towards different saprotrophic fungi (Fig. 3d) are the inhibitory rate of WT_Th_ minus the inhibitory rate when *nps1*_Th_ was deleted and normalized by the inhibitory rate of WT_Th_:

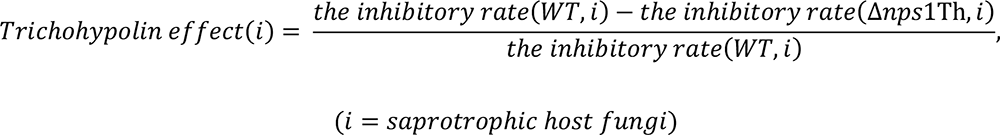

### Specific antagonism effects on host fungi by trichohypolin derivatives

10 mL PDA media containing 20 μg/mL trichohypolins A (**1**), B (**2**), or subfractions (SF1–SF5) in 0.1% DMSO were prepared in 60 mm petri dishes. Agar-mycelial plugs with 2 mm diameter from a 3-day-old culture were placed in the center of PDA plates. 10 mL PDA medium containing 0.1% DMSO was set as a control. All plates were cultivated at 25°C in the darkness. When the fungus grows to a mycelial area with a 60 mm diameter on the control plate, the growth inhibition was calculated as (R - r)/R x 100 (Extended Data Fig. 9). R is the radius of the saprotrophic fungal colony in the control plate, and r is the radius of the fungal colony in the treatment plate^49^. The clustering method of rows and columns in Fig. 4b was unweighted average distance (UPGMA) to present fungal species distance and inhibitory effect distance. Fungal distance was acquired by pairwise *p*-distance analysis of *LSU* nucleic acid sequences from different saprotrophic fungi. Inhibitory effect distance was calculated by inhibitory effect matrix. We termed the inhibitory effect pairwise distance of fungus i and fungus j as α(i, j), and i^th^ row in inhibitory effect matrix was a (1-by-n) row vector: X_i_ = x_i1_, x_i2_, …, x_in_, α(i, j) was defined as follows (Minkowski distance):

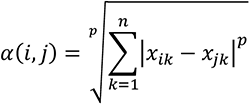

Effects of the minor components in SF1 (**3**–**8**) and SF2 (**9**–**13**) on a host fungus (Fig. 4c) was the inhibitory rate of mixture SF1 or SF2 except for the pure compound **1** or **2** towards different fungi subtracting the effect of compounds **1** or **2** with equal mass:

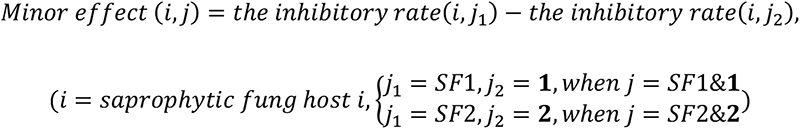

Mycohost distance was obtained based on Bayesian analysis of *LSU* nucleic acid sequences (Fig. 4b). Relationship analyses of inhibitory effect of trichohypolins with mycohost distance are carried out to calculate inhibitory effect distance (Fig. 4d). Three special cases of Minkowski distance to calculate inhibitory effect distance. For the special case of *p* = 1, the Minkowski distance gives the city block distance. For the special case of *p* = 2, the Minkowski distance gives the Euclidean distance. For the special case of *p* = ∞, the Minkowski distance gives the Chebychev distance.

### Structural diversity analysis of trichohypolins

For plotting the substrate promiscuity in *T. hypoxylon* (Extended Data Fig. 5), we first calculated the substrate frequency of each module by detected compound sequences (Fig. 2). Then we bulit a substrate promiscuity frequency matrix (n-by-n, n is the number of substrate types, the row is source and the column is target). First, one substrate (*e.g.* Q=0.71) frequency were divided equally to other activated substrates (*e.g.* U, E, E(Me)) in this module, and divided substrate frequency (0.71/3=0.24) was added up in the corresponding cell (*e.g.* Q (source)→U (target), Q (source)→E (target) and Q (source)→E(Me) (target)). Then we iterated through all modules and repeated this operation. Specially, the module which only had one substrate added one count in the cell of the diagonal line. Finally, this asymmetric matrix was used to plot the chord diagram by the function “chorddiag” in R script. This is the inner circle of Extended Data Fig. 5. The donut diagram (the outer of Extended Data Fig. 5) was ploted by the corresponding modules to each section.

### LC-MS analysis

Liquid chromatography tandem mass spectrometry (LC-MS) analysis for chemical profile was performed on an Agilent HPLC 1200 series system equipped with a single quadrupole mass selective detector and an Agilent 1100LC MSD model G1946D mass spectrometer by using a Venusil XBP C18 column (3.0 x 50 mm, 3 μm, Bonna-Agela Technologies, China). Acetonitrile (A) and Water with 0.1% (v/v) formic acid (B) were used as solvents at flow rate of 0.5 mL/min. The substances were eluted with a linear gradient from 5 to 100% solvent A in 30 min, then washed with 100% (v/v) solvent A for 5 min, and equilibrated with 5% (v/v) solvent A for 10 min. The mass spectrometer was set in electrospray positive ion mode for ionization.

The liquid chromatography-high resolution mass spectrometry (LC-HRMS) system was conducted from a Dionex Ultimate 3000 RSLC and Thermo Q-Exactive HRMS (Bremen, Germany). The analytes were separated on an analytical BEH C18 column (100 × 2.1 mm, 1.7 μm) with the 40°C column oven. Acetonitrile (A) and water with 0.1% formic acid (B) were used as solvents at a flow rate of 0.3 mL/min. The substances were eluted with a linear gradient from 0 to 40% (v/v) solvent A for 1.5 min, 40% (v/v) solvent A for 9.5 min, and then washed with 100% (v/v) solvent A for 2.5 min, followed by a pre-equilibrium for 3 min with 40% (v/v) solvent A. The HRMS instrument with an electrospray ionization (ESI) probe was tuned and calibrated in ESI^+^ using positive calibration solutions once a week. The MS parameters were set as follows: spray voltage (3.8 kV), capillary temperature (320°C), probe heater temperature (400°C), and S-Lens (60 V). The HRMS was acquired in full scan (FS) or data-dependent fragmentation acquisition (dd-MS_2_) mode. In FS, mass resolution, AGC target, and maximum IT were set at 70000 FWHM, 3.0×106, and 100 ms, respectively. In dd -MS^2^, mass resolution (17500 FWHM), AGC target (1.0×105), maximum IT (30 ms), Top N (5), and stepped normalized collision energy (NCE) (15%, 35% and 55%) were set.

### NMR analysis

NMR spectra were recorded on a Bruker Avance-500 MHz spectrometer at room temperature (Bruker Corporation, Karlsruhe, Germany). All spectra were processed with TopSpin 3.5 (Bruker Corporation, Karlsruhe, Germany). Chemical shifts are referenced to those of the solvent signals. NMR data are given in Supplementary Table S5 and spectra in Supplementary Fig. S3–S18.

### X-ray crystallographic analysis

Colorless crystals of trichohypolin A (**1**) were obtained in MeOH/H_2_O. Crystallographic data for **1** (Cu Ka radiation) has been deposited in the Cambridge Crystallographic Data Centre with the deposition number CCDC 2171447. These data can be obtained free of charge *via* www.ccdc.cam.ac.uk/data_request/cif.

## Acknowledgements

We thank Prof. Dr. Wen-Ying Zhuang from Institute of Microbiology, Chinese Academy of Sciences for helpful discussion and suggestion. This research was supported by National Key Research and Development Program of China (2020YFA0907800), National Natural Science Foundation of China (32170066), and Key Research Program of Frontier Sciences, Chinese Academy of Sciences (ZDBS-LY-SM016) as well as the Biological Resources Program, Chinese Academy of Sciences (KFJ-BRP-009-005). Jie Fan is a recipient from China Postdoctoral Science Foundation (YJ20200201 and 2020M680720).

## Author contributions

W.-B. Y. conceived of the project and revised the manuscript. J. F., P.-L. Wei, and Y. Li performed experiments including gene deletion, biological activity screening of purified trichohypolins and peptaibol subfractions, as well as antagonism analysis. J. F. and J. R. performed trichohypolin isolation and identification, completed data analysis and wrote the manuscript with input from all authors. W. L. helped concerning fungal isolation. I. S. D. provided assistance and guidance for fungal identification, discussed and revised the manuscript. Z. L. and R. H. performed the majority of computational analyses. D. C. performed ESI-HRMS analysis. All authors read and approved the final manuscript.

## Additional information

**Extended data** is available for this paper at xxxxx.

**Supplementary Information** The online version contains supplementary material available at https://doi.org/xxxxxx.

**Correspondence and requests for materials** should be addressed to W.-B. Y..

**Peer review Information** *Nature Chemical Biology* thanks xxx, xxx and the other, anonymous, reviewer(s) for their contribution to the peer review of this work.

**Reprints and permissions Information** is available at http://www.nature.com/reprints.

---

**Extended Data Fig. 1.**
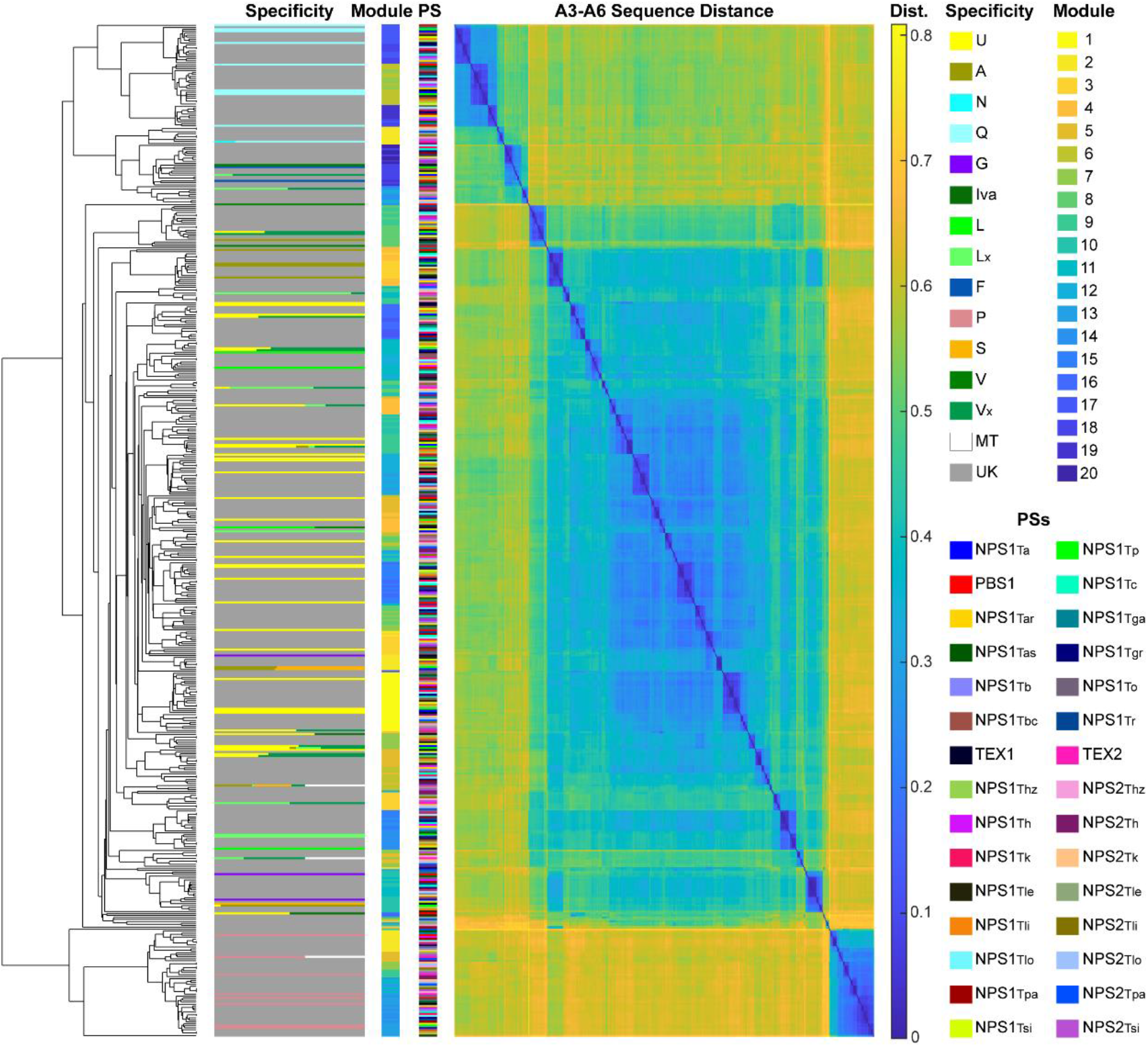
Prediction of peptaibol sequences possibly synthesized by potential peptaibol synthetases in *Trichoderma* genus. 513 A domain sequences of peptaibol synthetases (PSs) are clustered as a phylogenetic tree on the left. Modules in PSs and their substrate specificities are attached to the multiple sequence alignment (the 2^nd^ and 3^rd^ columns from left). Twenty-one PSs containing 18–20 modules and nine PSs containing 12–15 modules are shown. PSs are listed by colors in lower right panel (details see Supplementary Information Table S3). Distance matrix of A domain sequences is sorted using motif A3–A6 alignment, which overlaps the substrate’s binding pocket^24^. Modules of PSs and their corresponding specific substrates are shown by colors in side bars. Substrates with similar chemical properties are represented by similar colors. L_x_, L or I; V_x_, V or Iva; MT, empty, that means this module could be skipped in the product synthesis; UK, unknown, that means the corresponding products of PS remain unknown. Amino acid sequences of PSs were obtained from the GenBank database.

**Extended Data Fig. 2.**
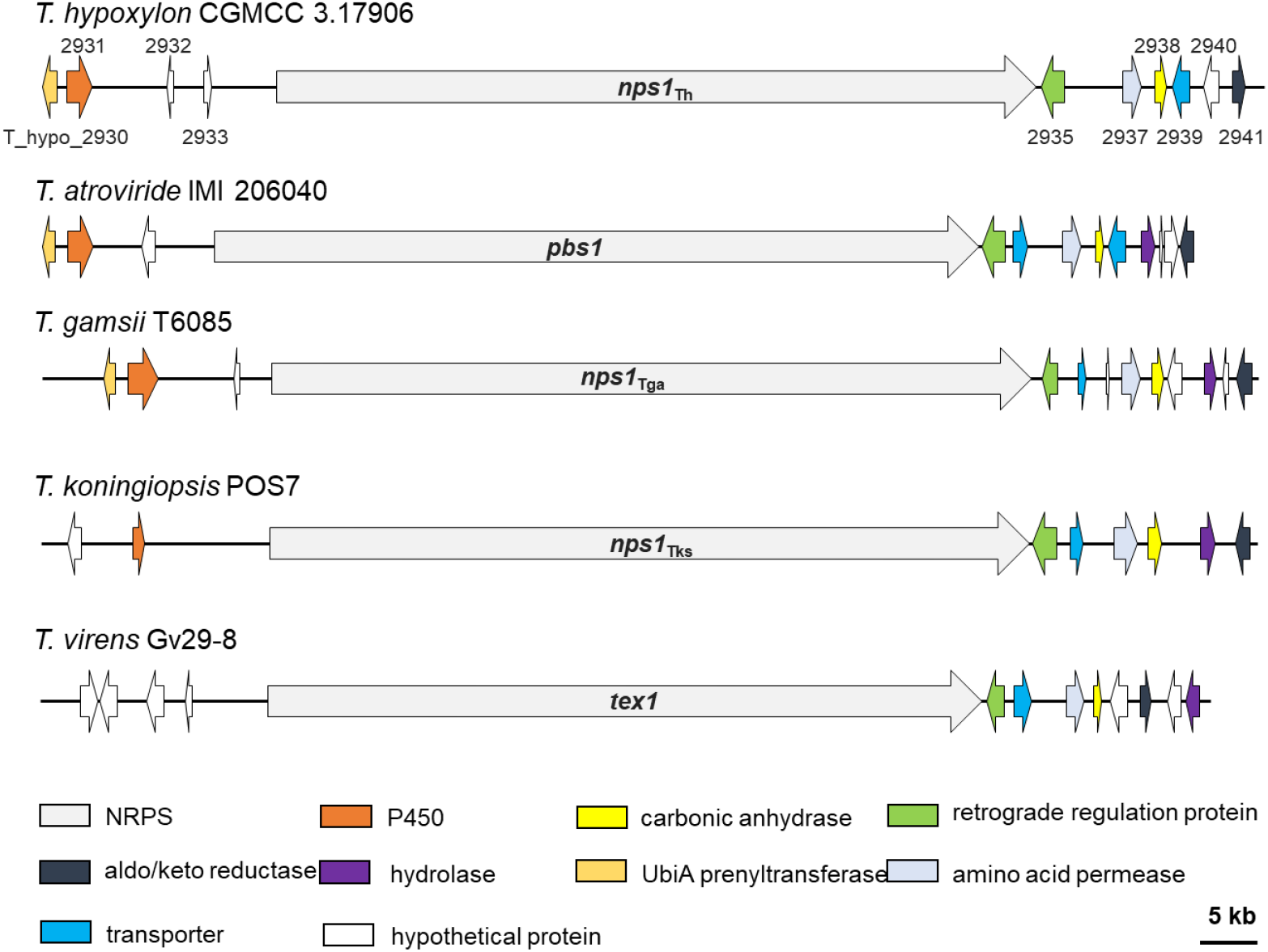
Representative biosynthetic gene clusters of peptaibols in *Trichoderma* genus. Peptaibol biosynthetic gene clusters (BGCs) containing peptaibol synthetase encoding genes in the genomes of *T. hypoxylon*, *T. atroviride*, *T. gamsii*, *T. koningiopsis*, and *T. virens*. For example, a BGC of trichohypolins is composed of a NRPS encoding gene *nps1*_Th_, a P450 oxygenase encoding gene, a carbonic anhydrase encoding gene, a UbiA prenyltransferase encoding gene, a reductase encoding gene, a hydrolase encoding gene, a retrograde regulator, two putative transporters, and three other hypothetical genes from *T. hypoxylon*. *nps1*_Th_ from *T. hypoxylon*, *pbs1* from *T. atroviride*, *nps1*_Tga_ from *T. gamsii*, and *nps1*_Tks_ from *T. koningiopsis* with 19 modules; *tex1* from *T. virens* with 18 modules. Gene functions were shown with different colors as seen in the lower panel.

**Extended Data Fig. 3.**
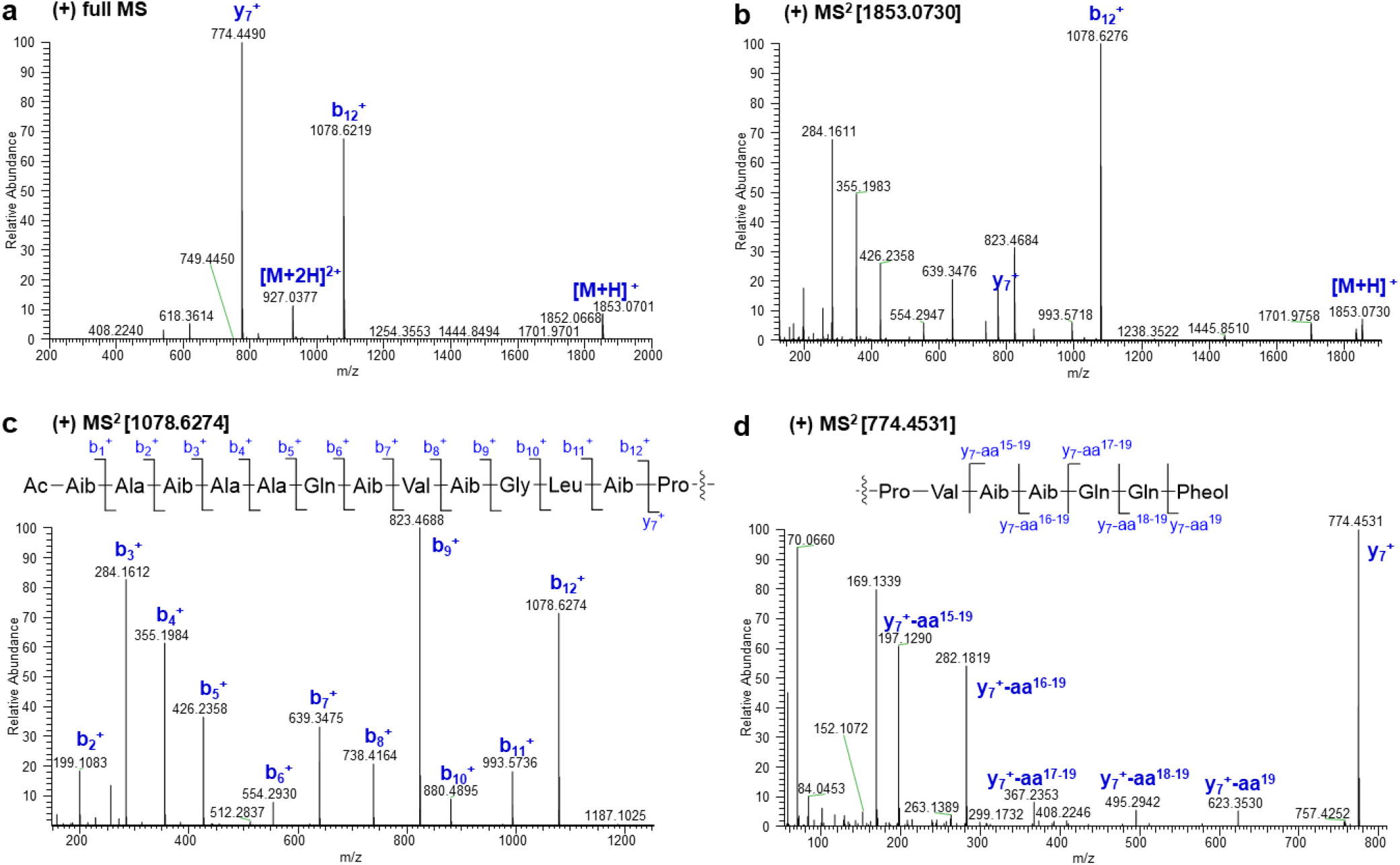
Diagnostic fragment ions [*m/z*] of trichohypolin A (1) by ESI-HRMS analysis.

**Extended Data Fig. 4.**
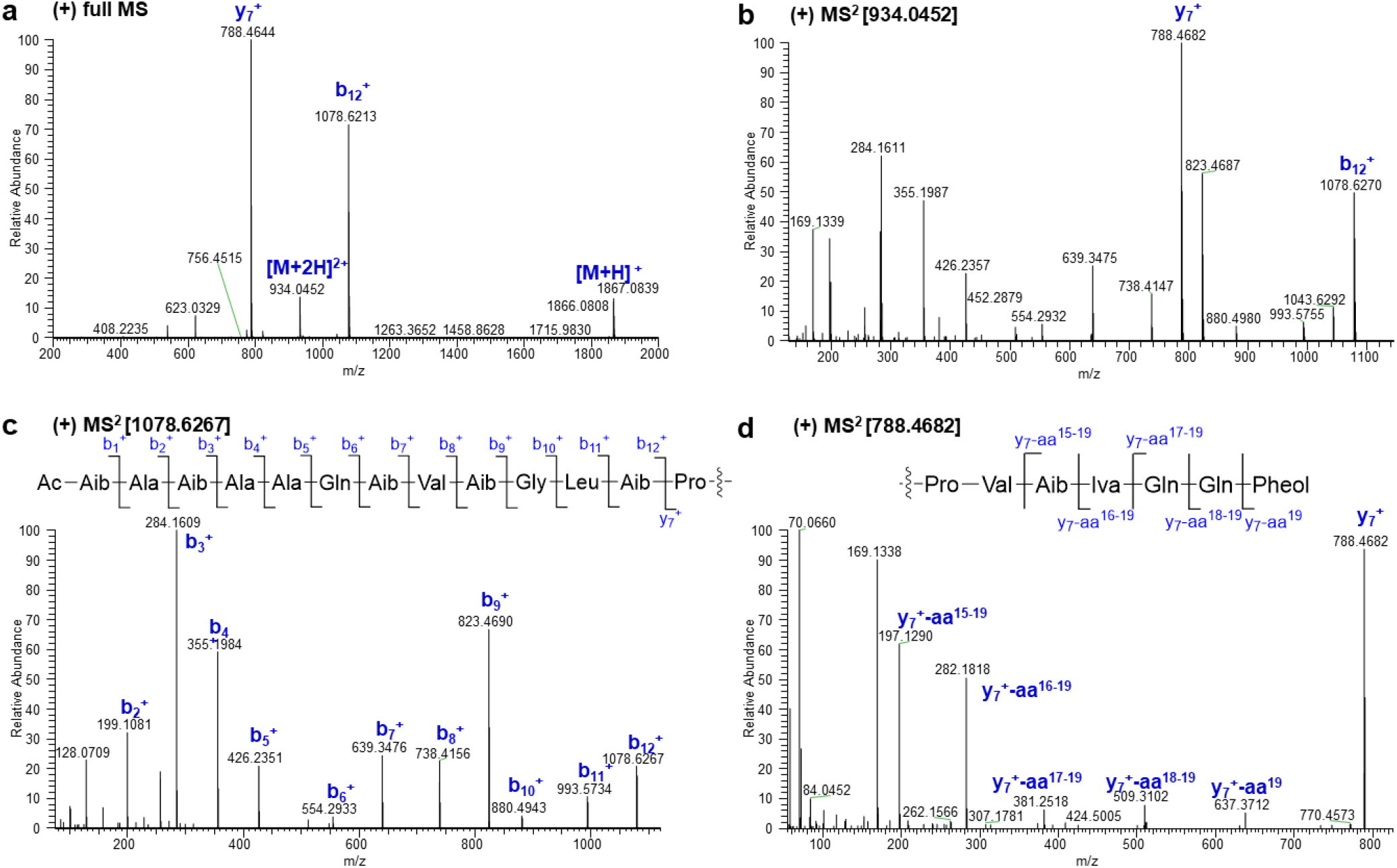
Diagnostic fragment ions [*m/z*] of trichohypolin B (2) by ESI-HRMS analysis.

**Extended Data Fig. 5.**
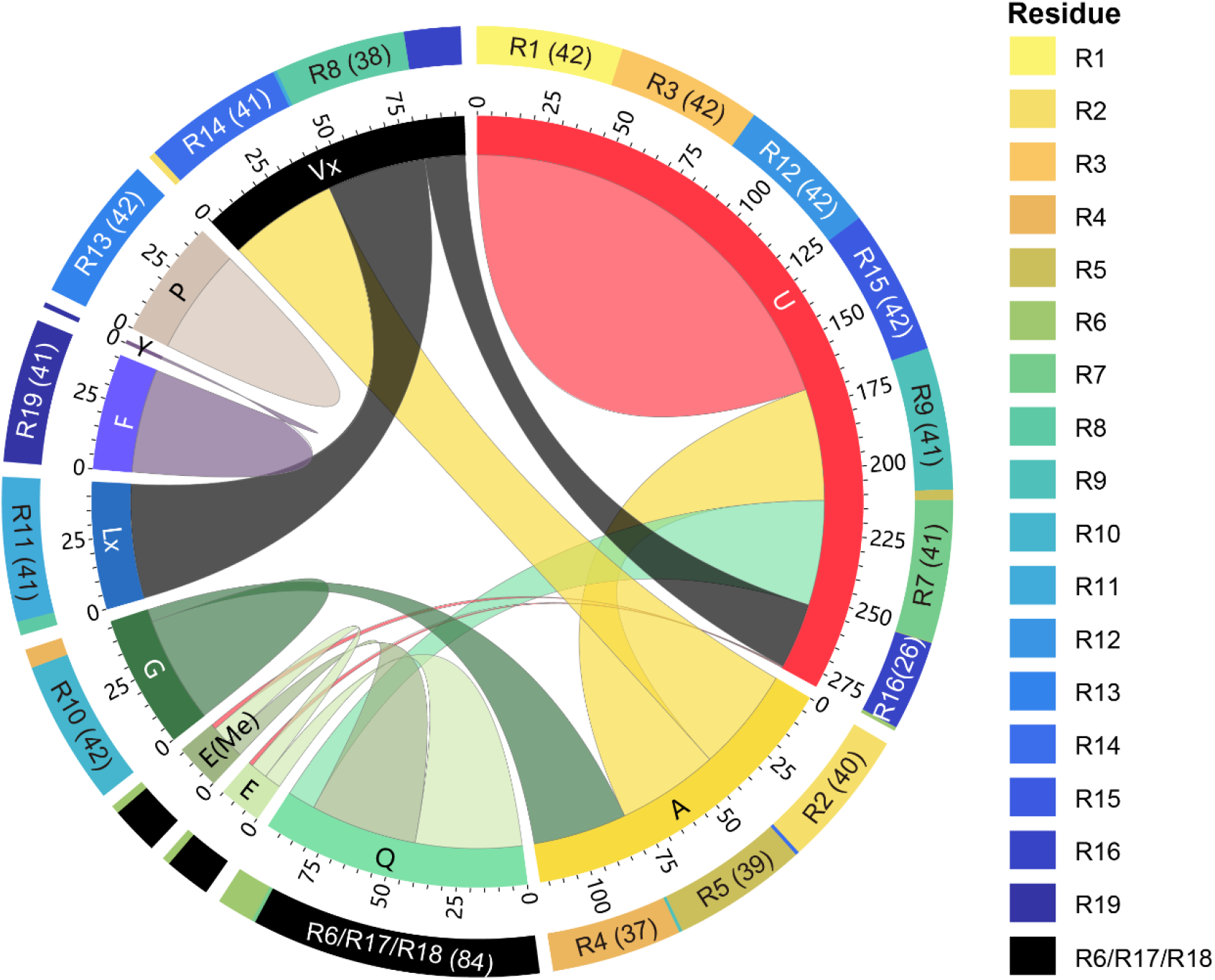
Structual diversity of trichohypolins and substrate promiscuity of each module in NPS1Th from *T. hypoxylon*. Structures of 42 trichohypolin derivatives are compared for plotting the substrate promiscuity of the peptaibol synthetase NPS1_Th_. 19 amino acid residues of each trichohypolin are summarized by chord diagram. The source and target of each chord represent two substrates which could be activated by the same module (see method). Residues are shown in different colors, and their corresponding modules are ploted by donut diagram. Numbers in brackets represent the counts of modules (>25) activating the corresponding amino acids. U, Aib; L_x_, L or I; V_x_, V or Iva; E(Me), methylated glutamate.

**Extended Data Fig. 6.**
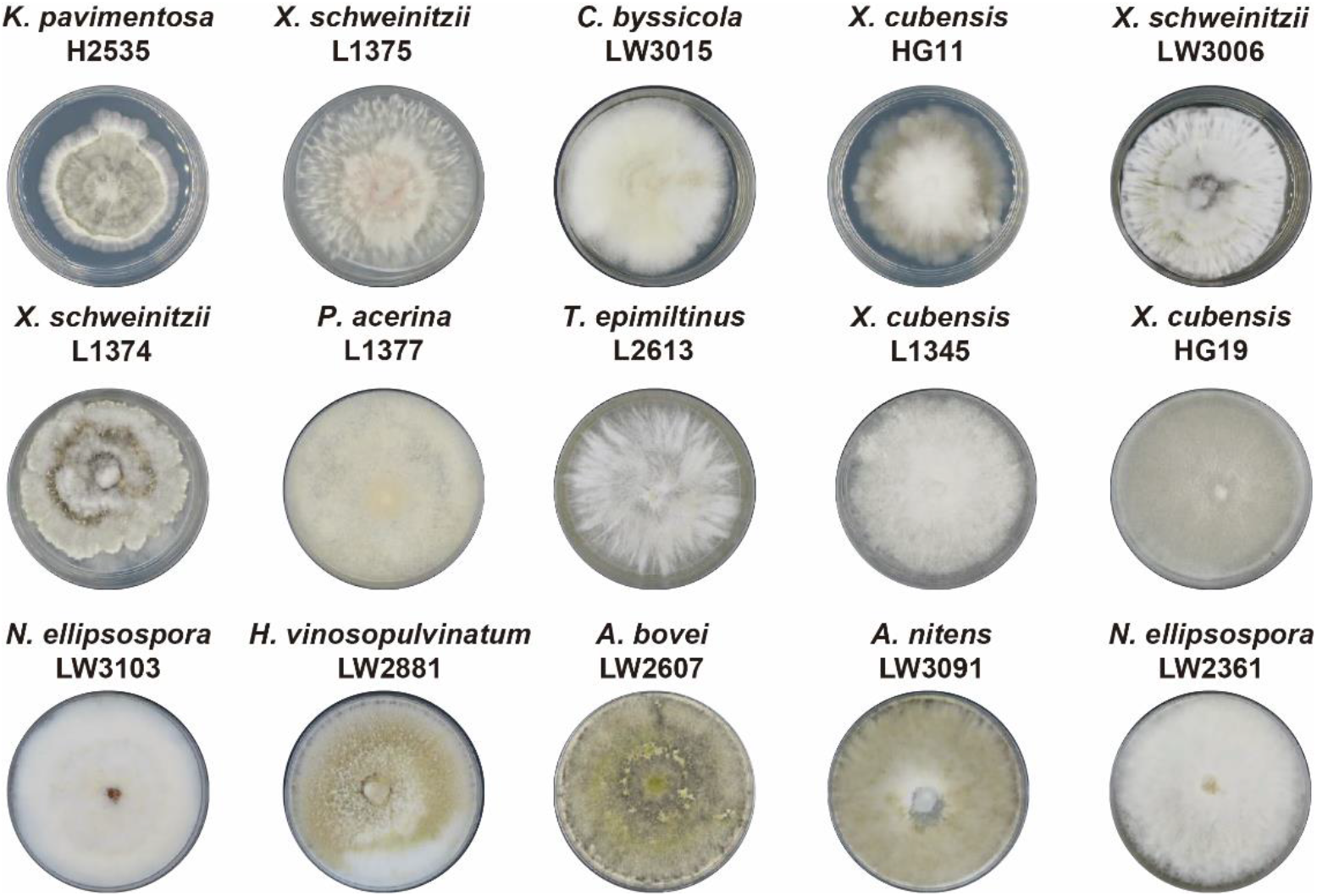
Morphologies of 15 saprotrophic fungi used in this study. 15 saprotrophic fungi were identified by phylogenetic analysis (see Extended Data Fig. 7) of *ITS*, *LSU*, *RPB2*, and *β-tubulin* as DNA barcode fragments referring to the corresponding reports about species identification. They are *X. schweinitzii* L1374, *X. schweinitzii* L1375, *X. schweinitzii* LW3006^50^, *X. cubensis* L1345, *X. cubensis* HG19, *X. cubensis* HG11^51^, *H. vinosopulvinatum* LW2881^52^, *A. bovei* LW2607^53^, *A. nitens* LW3091^54^, *K. pavimentosa* H2535^55^, *P. acerina* L1377^56^, *N. ellipsospora* LW2361, *N. ellipsospora* LW3103^57^, *C. byssicola* LW3015^58, 59^, and *T. epimiltinus* L2613^60^.

**Extended Data Fig. 7.**
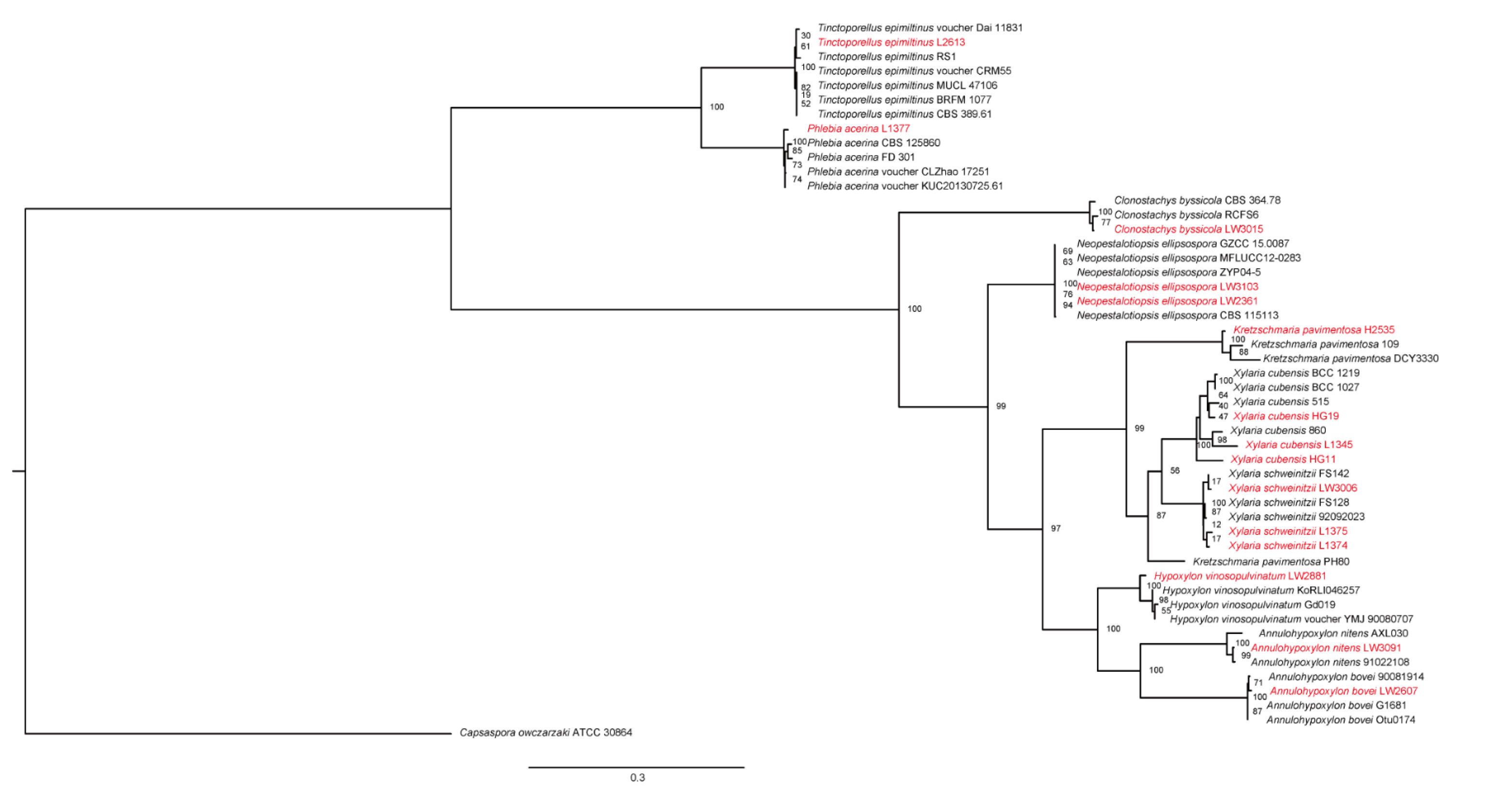
Phylogenetic analysis of 15 saprotrophic fungal strains.

**Extended Data Fig. 8.**
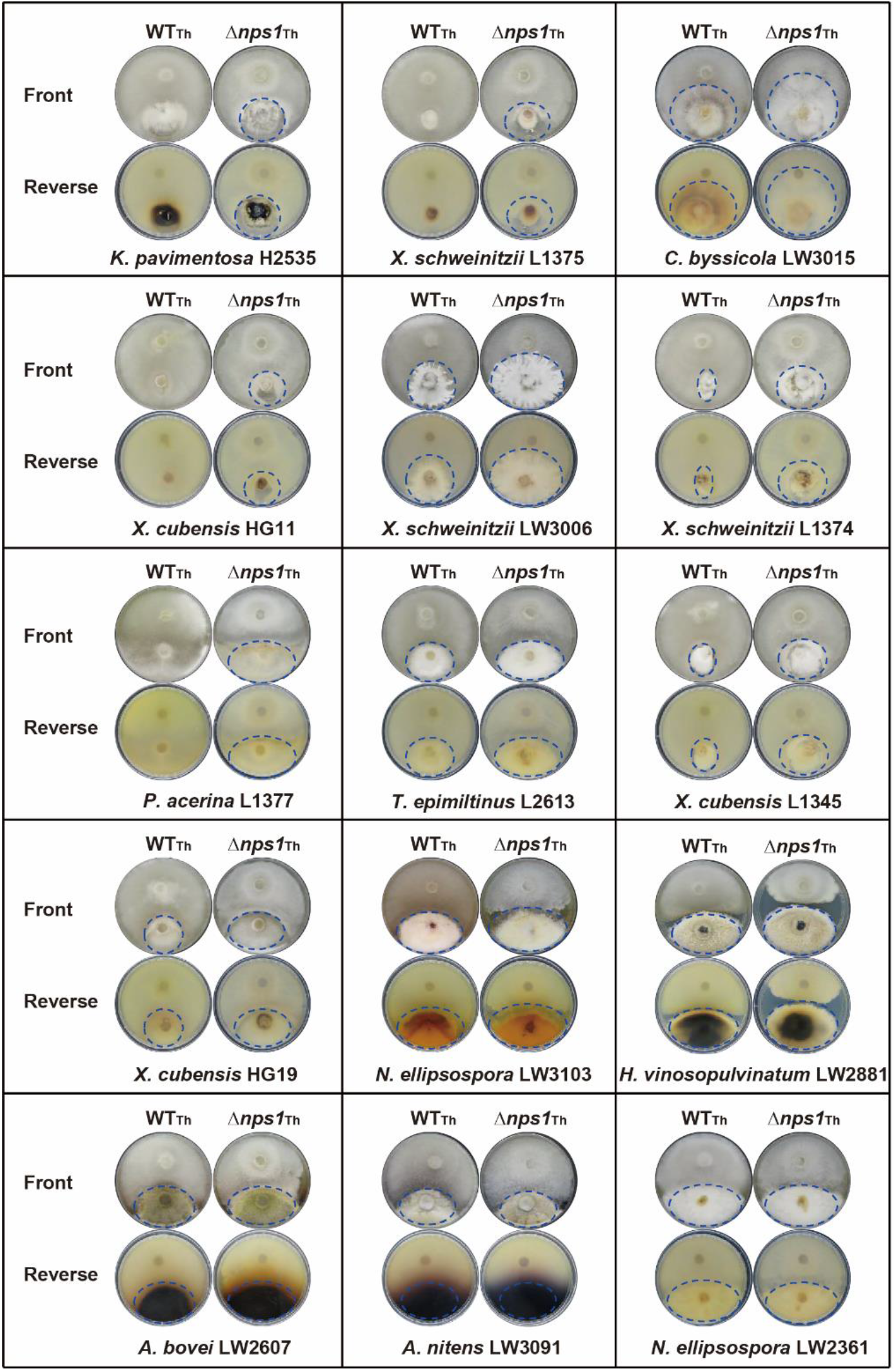
Cocultivation of *T. hypoxylon* wild type (WTTh) and Δ*nps1*Th with 15 saprotrophic fungi. Confrontation assay of *T. hypoxylon* (above) with a saprotrophic host fungus (below). Culture front and reverse of plates are shown, with blue circles indicating the extension of saprotrophic fungal colonies. Three replicates are carried out for growth inhibiory calculation. Scheme of experimental setup and the calculation method are shown in Fig. 3.

**Extended Data Fig. 9.**
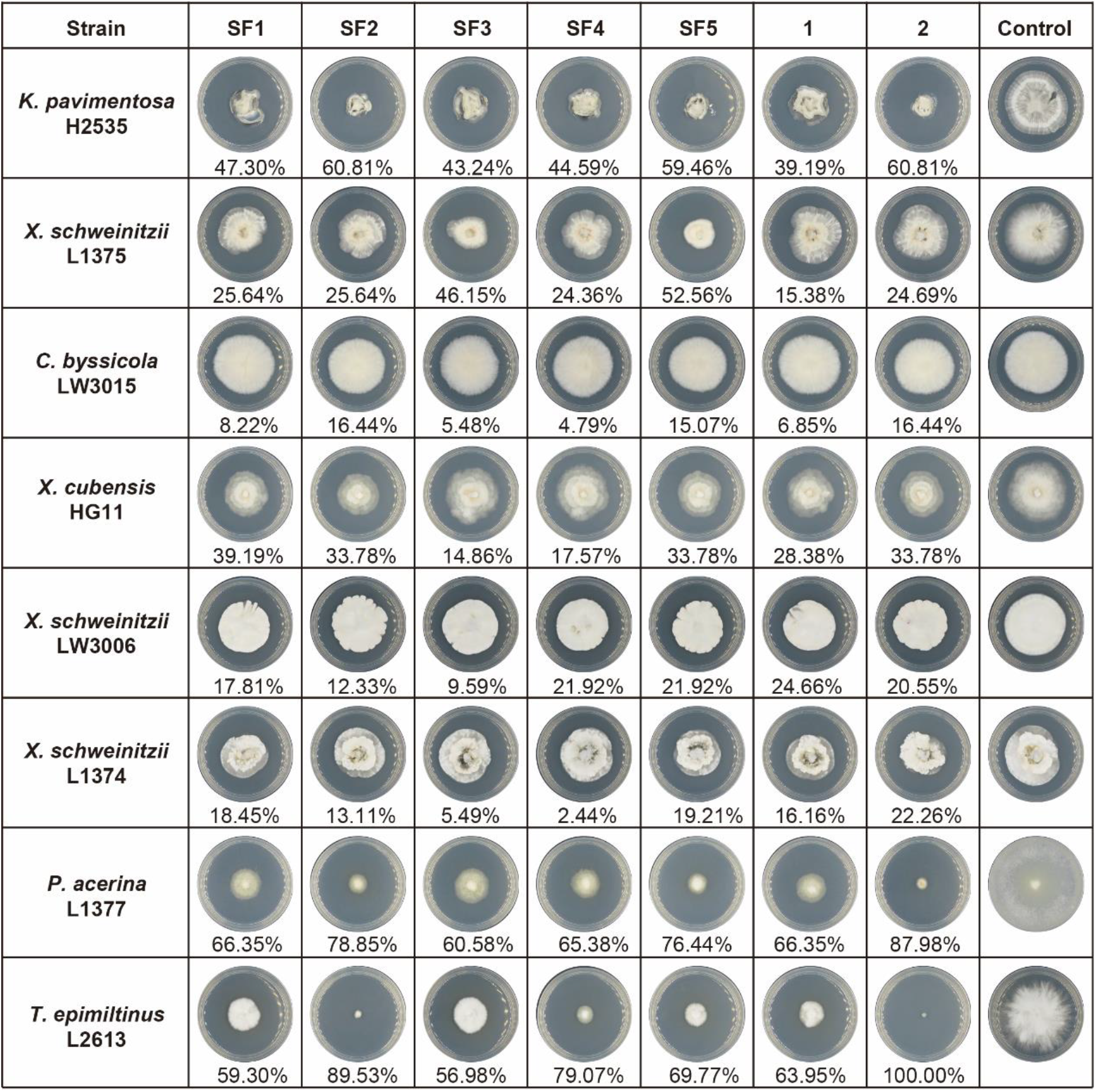
Antagonistic abilities of trichohypolins A (1) and B (2), as well as subfractions (SF1–SF5) against saprotrophic fungi.

