## Supplementary Table S1-S7, Supplementary Fig. S1-S58 for "Biosynthetic diversification of peptaibol mediates fungus-mycohost interactions"

**Fig. S12 (B) 1D TOCSY spectrum trichohypolin A (1) with selective excitation of 2-NH of Gln$^{18}$ (δ 7.51)22**

**Fig. S12 (C) 1D TOCSY spectrum trichohypolin A (1) with selective excitation of 2-NH of Gln$^{17}$ (δ 7.93)22**

**Fig. S12 (D) 1D TOCSY spectrum trichohypolin A (1) with selective excitation of 2-NH of Val$^{8}$ (δ 7.66)22**

**Fig. S12 (E) 1D TOCSY spectrum trichohypolin A (1) with selective excitation of 2-NH of Leu$^{11}$ (δ 7.74)23**

**Fig. S12 (F) 1D TOCSY spectrum trichohypolin A (1) with selective excitation of 2-NH of Ala$^{4}$ (δ 8.01)23**

**Fig. S12 (G) 1D TOCSY spectrum trichohypolin A (1) with selective excitation of 2-NH of Ala$^{5}$ (δ 7.83)23**

**Fig. S12 (H) 1D TOCSY spectrum trichohypolin A (1) with selective excitation of 2-NH of Gln$^{6}$ (δ 7.61)24**

**Table S1.** Strains used in this study

| Strain | Abbreviation | Genotype | Source |
|---|---|---|---|
| *Trichoderma hypoxylon* CGMCC 3.17906 | | wild type ($WT_{Th}$) | [1] |
| TYYL2.1 | | Δ*nps1*$_{Th}$::*hph* in *T. hypoxylon* $WT_{Th}$ | This study |
| *Kretzschmaria pavimentosa* H2535 | KP | wild type, from Wuzhi mountain, Hainan, China | This study |
| *Xylaria schweinitzii* L1375 | $XS_a$ | wild type from Dayaoshan natural reserve, Guangxi, China | This study |
| *Xylaria schweinitzii* LW3006 | $XS_b$ | wild type from Xishuangbanna tropical botanical garden, Yunnan, China | This study |
| *Xylaria schweinitzii* L1374 | $XS_c$ | wild type from Dayaoshan natural reserve, Guangxi, China | This study |
| *Xylaria cubensis* HG11 | $XC_a$ | wild type from Datang Bay, Leigong mountain natural reserve, Leishan County Guizhou, China | This study |
| *Xylaria cubensis* L1345 | $XC_b$ | wild type from Maoershan natural reserve, Guangxi, China | This study |
| *Xylaria cubensis* HG19 | $XC_c$ | wild type from Datang Bay, Leigong mountain natural reserve, Leishan County, Guizhou, China | This study |
| *Clonostachys byssicola* LW3015 | CB | wild type from Mengyang wild elephant valley, Jinghong, Yunnan, China | This study |
| *Phlebia acerina* L1377 | PA | wild type from Dayaoshan natural reserve, Guangxi, China | This study |
| *Tinctoporellus epimiltinus* L2613 | TE | wild type from Chebaling, Shixing County, Guangdong, China | This study |
| *Hypoxylon vinosopulvinatum* LW2881 | HV | wild type, from Xishuangbanna tropical botanical garden, Yunnan, China | This study |
| *Annulohypoxylon bovei* LW2607 | AB | wild type from Kuankuoshui natural reserve, Suiyang County, Guizhou, China | This study |
| *Annulohypoxylon nitens* LW3091 | AN | wild type from Jinghong Jinuo, Yunnan, China | This study |
| *Neopestalotiopsis ellipsospora* LW3103 | $NE_a$ | wild type from Dadugang, Jinghong, Yunnan, China | This study |
| *Neopestalotiopsis ellipsospora* LW2361 | $NE_b$ | wild type, from Datang Bay, Leigong mountain natural reserve, Leishan County, Guizhou, China | This study |

**Table S2.** Primers used in this study

| Primers | Sequence 5' to 3' | Targeted amplification |
|---|---|---|
| hypoPS-RT-F | gctcactagtgaactccgg | 1262 bp partial fragment of *nps1*$_{Th}$ |
| hypoPS-RT-R | ggtacacaccgagagaagc | |
| hypoPS-5F-F | cagctctccacaatgtgctc | 2763 bp upstream fragment of *nps1*$_{Th}$ |
| hypoPS-5F-R | taactgtgataaactaccgcattaaagctgcactgttgcatactcgtgg | |
| hypoPS-3F-F | aattgcgcgcttggcgtaatcatggtcccagatgcaatgtgtgaatgtg | 2769 bp downstream fragment of *nps1*$_{Th}$ |
| hypoPS-3F-R | gttctcgtggattgaggcag | |
| hypoPS-nest-F | gtctctcgcatgcggatgag | 7528 bp fragment containing upstream and downstream of *nps1*$_{Th}$ as well as *hph* gene |
| hypoPS-nest-R | ggcgagagaactcgtgagag | |
| hyp-scr-5F-R | gcctatgcctacagcatcc | upstream and downstream of hph to verify $\Delta$*nps1*$_{Th}$ mutant |
| hyp-scr-3F-F | cgtggtcgagctacaaagc | |
| ITS1 | tccgtaggtgaacctgcgg | internal transcribed spacer regions and intervening 5.8S nrDNA (ITS) |
| ITS2 | gctgcgttcttcatcgatgc | |
| LR0R | acccgctgaacttaagc | partial 28S large subunit nrDNA (LSU) |
| LR7 | tactaccaccaagatct | |
| RPB2-5F2 | ggggwgaycagaagaaggc | the β-tubulin (*tub2*) gene region |
| RPB2-7cR | cccatrgcttgtyyrcccat | |
| Btub2Fd | gtbcacctycaraccggycartg | partial regions of RNA polymerase II second largest subunit (*rpb2*) |
| Btub4Rd | ccrgaytgrccraaracraagttgtc | |

**Table S3.** Peptaibol synthetases (PSs) with >10 A domains throughout *Trichoderma* genus

| Strain | No. of PSs | Name | Accession number | No. of A domain | Length (aa) | Peptaibol products |
|---|---|---|---|---|---|---|
| *T. hypoxylon* CGMCC 3.17906 | 2 | $NPS1_{Th}$ | | 19 | 22,052 | This study |
| | | $NPS2_{Th}$ | | 15 | 17,728 | This study |
| *T. virens* Gv29-8 | 2 | TEX1[2] | AAM78457.1 | 18 | 20,925 | [2] |
| | | TEX2[3] | XP_013957420.1 | 14 | 16,510 | [3] |
| *T. afroharzianum* MRI39 | 1 | $NPS1_{Taf}$ | | 17 | 20,508 | [4] |
| *T. aggressivum* f. europaeum 433.95 | 1 | $NPS1_{Ta}$[5] | AQV12034.1 | 18 | 20,978 | [5] |
| *T. arundinaceum* IBT 40837 | 2 | $NPS1_{Tar}$ | RFU80400.1 | 20 | 23,058 | [6] |
| | | $NPS2_{Tar}$ | RFU79003.1 | 11 | 13,245 | |
| *T. asperellum* CBS 433.97 | 1 | $NPS1_{Tas}$ | XP_024761182.1 | 18 | 20,866 | [7-9] |
| *T. atrobrunneum* ITEM 908 | 1 | $NPS1_{Tat}$ | | 18 | 20,519 | |
| *T. atroviride* IMI 206040 | 1 | PBS1[10] | XP_013944039.1 | 19 | 21,901 | [10-12] |
| *T. brevicompactum* IBT40841 | 1 | $NPS1_{Tb}$ | | 20 | 23,015 | [13] |
| *T. brevicrassum* TC967 | 2 | $NPS1_{Tbc}$ | | 18 | 20,735 | |
| | | $NPS2_{Tbc}$ | | 14 | 16,337 | |
| *T. citrinoviride* TUCIM 6016 | 2 | $NPS1_{Tc}$ | XP_024748081.1 | 18 | 20,874 | [14] |
| | | $NPS2_{Tc}$ | XP_024752339.1 | 10 | 13,601 | |
| *T. erinaceum* CRRI-T2N1 | 1 | $NPS1_{Te}$ | | 17 | 21,419 | |
| *T. gamsii* T6085 | 2 | $NPS1_{Tga}$ | XP_024405762.1 | 19 | 21,960 | [15] |
| | | $NPS2_{Tga}$ | XP_024405290.1 | 10 | 12,640 | |
| *T. gracile* HK011-1 | 2 | $NPS1_{Tgr}$ | KAH0491796.1 | 20 | 22,994 | |
| | | $NPS2_{Tgr}$ | KAH0492734.1 | 11 | 14,293 | |
| *T. guizhouense* NJAU 4742 | 1 | $NPS1_{Tgu}$ | OPB36439.1 | 12 | 15,994 | |
| *T. hamatum* GD12 | 1 | $NPS1_{Thm}$ | | 15 | 19,914 | |
| *T. harzianum* CBS 226.95 | 2 | $NPS1_{Thz}$ | XP_024769971.1 | 18 | 20,877 | [16-20] |
| | | $NPS2_{Thz}$ | XP_024775665.1 | 12 | 14,364 | |
| *T. koningii* JCM 1883 | 2 | $NPS1_{Tk}$ | | 20 | 23,040 | [21] |
| | | $NPS2_{Tk}$ | | 12 | 14,878 | |
| *T. koningiopsis* POS7 | 1 | $NPS1_{Tks}$ | | 19 | 21,752 | [15] |
| *T. lentiforme* CFAM-422 | 2 | $NPS1_{Tle}$ | KAF3066167.1 | 18 | 20,735 | |
| | | $NPS2_{Tle}$ | KAF3068249.1 | 14 | 16,614 | |
| *T. lixii* MUT 3171 | 2 | $NPS1_{Tli}$ | | 18 | 20,759 | |
| | | $NPS2_{Tli}$ | | 12 | 14,187 | |
| *T. longibrachiatum* ATCC 18648 | 2 | $NPS1_{Tlo}$ | PTB76898.1 | 20 | 23,089 | [9,22,23] |
| | | $NPS2_{Tlo}$ | PTB77615.1 | 12 | 14,409 | |
| *T. oligosporum* CGMCC 3.17527 | 1 | $NPS1_{To}$ | | 20 | 23,025 | |
| *T. parareesei* CBS 125925 | 2 | $NPS1_{Tpa}$ | OTA01063.1 | 20 | 22,960 | |
| | | $NPS2_{Tpa}$ | OTA02976.1 | 14 | 16,450 | |
| *T. pleuroti* Tphu1 | 1 | $NPS1_{Tp}$[5] | AQV12033.1 | 18 | 20,962 | [5] |

**Table S3.** Peptaibol synthetases (PSs) with >10 A domains over *Trichoderma* species (continued)

| Strain | No. of PSs | Name | Accession number | No. of A domain | Length (aa) | Peptaibol products |
|---|---|---|---|---|---|---|
| *T. reesei* QM6a | 2 | $NPS1_{Tr}$ | XP_006968566.1 | 18 | 20,873 | [24] |
| | | $NPS2_{Tr}$ | XP_006968900.1 | 14 | 16,534 | |
| *T. simmonsii* GH-Sj1 | 2 | $NPS1_{Tsi}$ | QYT04456.1 | 18 | 20,748 | [4] |
| | | $NPS2_{Tsi}$ | QYS95804.1 | 14 | 16,622 | |
| *T. semiorbis* FJ059 | 1 | $NPS1_{Tse}$ | | 15 | 19,899 | |
| *Trichoderma* sp. IMV 00454 | 1 | $NPS1_{454}$ | | 13 | 18,392 | |
| *Trichoderma* sp. TW21990 | 1 | $NPS1_{990}$ | | 16 | 19,846 | |

**Table S4.** Blast analysis of HypoPS1 from *T. hypoxylon* saprophytic fungi

| **Strain** | **Name** | **Gene ID** | **Cover/Identity** | **No. of A domain** | **Length (aa)** |
|---|---|---|---|---|---|
| *T. virens* Gv29-8 | Tex1 | AAM78457.1 (XP_013953110.1) | 99/72.80 | 18 | 20,925 |
| *T. atroviride* IMI 206040 | AtvPS/Pbs1 | XP_013944039.1 | 99/76.29 | 19 | 21,901 |
| *T. gamsii* T6085 | GamPS1 | XP_024405762.1 | 99/76.34 | 19 | 21,960 |
| *T. koningiopsis* POS7 | KosPS | | 99/73.66 | 19 | 21,752 |
| *T. arundinaceum* IBT 40837 | ArdPS1 | RFU80400.1 | 99/80.28 | 20 | 23,058 |
| *T. longibrachiatum* ATCC 18648 | LogPS1 | PTB76898.1 | 99/75.19 | 20 | 23,089 |

**Table S5.** $^{1}H$ and $^{13}C$ NMR data of trichohypolins A (**1**) and B (**2**) in DMSO-$d_6$

| | | trichohypolin A (**1**) | | | trichohypolin B (**2**) | |
|---|---|---|---|---|---|---|
| residue | position | $\delta_C{}^a$, mult. | $\delta_H{}^b$ ($J$ in Hz) | residue | $\delta_C{}^a$, mult. | $\delta_H{}^b$ ($J$ in Hz) |
| **Ac** | 1 | 171.1 | | **Ac** | 171.1 | |
| | 2 | 23.6 | 1.94 (s) | | 23.6 | 1.93 (s) |
| **Aib¹** | 1 | 176.4 | | **Aib¹** | 176.4 | |
| | 2 | 55.9 | | | 55.9 | |
| | 2-NH | | 8.58 (s) | | | 8.53 (s) |
| | 3 | 26.1~27.0 | ~1.46 (o) | | 26.2~26.9 | ~1.46 |
| | 4 | 23.2~23.6 | ~1.35 (o) | | 23.2~23.6 | ~1.35 |
| **Ala²** | 1 | 175.5 | | **Ala²** | 175.9 | |
| | 2 | 51.8 | 3.96 (o) | | 51.8 | 3.96 |
| | 2-NH | | 8.41 (brs) | | | 8.23 (brs) |
| | 3 | 16.8 | 1.35 (o) | | 16.8 | 1.35 (o) |
| **Aib³** | 1 | 176.8 | | **Aib³** | 176.8 | |
| | 2 | 56.6 | | | 56.6 | |
| | 2-NH | | 7.96 (s) | | | 7.90 (s) |
| | 3 | 26.1~27.0 | ~1.46 (o) | | 26.2~26.9 | ~1.46 |
| | 4 | 23.2~23.6 | ~1.35 (o) | | 23.2~23.6 | ~1.35 |
| **Ala⁴** | 1 | 175.3 | | **Ala⁴** | 175.3 | |
| | 2 | 52.0 | 3.95 (o) | | 52.1 | 3.96 |
| | 2-NH | | 8.01 (brs) | | | 7.96 |
| | 3 | 16.8 | 1.35 (o) | | 16.8 | 1.35 |
| **Ala⁵** | 1 | 176.0 | | **Ala⁵** | 176.0 | |
| | 2 | 52.0 | 3.95 (o) | | 52.0 | 3.96 |
| | 2-NH | | 7.83 (brs) | | | 7.80 |
| | 3 | 16.5 | 1.42 (o) | | 16.5 | 1.42 |
| **Gln⁶** | 1 | 174.1 | | **Gln⁶** | 174.1 | |
| | 2 | 55.1 | 3.87 (m) | | 55.9 | 3.87 |
| | 2-NH | | 7.61 (brs) | | | 7.62 |
| | 3 | 26.9 | 2.34 (m), 1.97 (m) | | 26.3 | 2.16 (o), 1.98 (o) |
| | 4 | 32.0 | 2.09 (m), 2.20 (m) | | 32.2 | 2.36 (m), 1.99 (o) |
| | 5 | 173.4 | | | 173.3 | |
| | 5-$NH_2$ | | 6.72 (brs), 7.21 (brs) | | | 6.72 (brs), 7.21 (brs) |
| **Aib⁷** | 1 | 176.1 | | **Aib⁷** | 176.0 | |
| | 2 | 55.3 | | | 55.9 | |
| | 2-NH | | 7.62 (s) | | | 7.64 (o) |
| | 3 | 26.1~27.0 | ~1.46 (o) | | 26.2~26.9 | ~1.46 |
| | 4 | 23.2~23.6 | ~1.35 (o) | | 23.2~23.6 | ~1.35 |

o, overlapping signals

**Table S5.** $^{1}$H and $^{13}$C NMR data of trichohypolins A (**1**) and B (**2**) in DMSO-$d_6$ (continued)

| | | trichohypolin A (**1**) | | | trichohypolin B (**2**) | |
|---|---|---|---|---|---|---|
| residue | position | $\delta_C$$^a$, mult. | $\delta_H$$^b$ ($J$ in Hz) | residue | $\delta_C$$^a$, mult. | $\delta_H$$^b$ ($J$ in Hz) |
| **Val$^8$** | 1 | 173.6 | | **Val$^8$** | 173.8 | |
| | 2 | 63.5 | 3.56 (o) | | | 3.58 (o) |
| | 2-NH | | 7.66 (d, 4.27) | | | 7.63 (o) |
| | 3 | 29.1 | 2.13 (o) | | 29.1 | 2.14 (o) |
| | 4 | 19.5 | 0.89 (s) | | 19.5 | 0.88 (o) |
| | 5 | 20.4 | 1.00 (s) | | 20.3 | 1.00 (6.4) |
| **Aib$^9$** | 1 | 176.4 | | **Aib$^9$** | 176.3 | |
| | 2 | 56.3 | | | 56.3 | |
| | 2-NH | | 7.91 (s) | | | 7.90 (o) |
| | 3 | 26.1~27.0 | ~1.46 (o) | | 26.2~26.9 | ~1.46 (o) |
| | 4 | 23.2~23.6 | ~1.35 (o) | | 23.2~23.6 | ~1.35 (o) |
| **Gly$^{10}$** | 1 | 170.1 | | **Gly$^{10}$** | 170.1 | |
| | 2 | 44.3 | 3.49 (o), 3.76 (o) | | 44.3 | 3.49 (o), 3.77 (o) |
| | 2-NH | | 8.13, (t, 5.13) | | | 8.11 (t, 5.24) |
| **Leu$^{11}$** | 1 | 173.9 | | **Leu$^{11}$** | 173.8 | |
| | 2 | 52.0 | 4.29 (o) | | 52.0 | 4.30 (o) |
| | 2-NH | | 7.74 (o) | | | 7.73 (o) |
| | 3 | 40.3 | 1.48 (o) | | 40.2 | 1.49 (o) |
| | | | 1.80 (o) | | | 1.80 (o) |
| | 4 | 24.5 | 1.78 (o) | | 24.5 | 1.78 (o) |
| | 5 | 21.3 | 0.82 (d, 6.1) | | 21.4 | 0.82 (d, 5.90) |
| | 6 | 23.2 | 0.86 (d, 6.1) | | 23.2 | 0.86 (d, 5.90) |
| **Aib$^{12}$** | 1 | 173.5 | | **Aib$^{12}$** | 173.8 | |
| | 2 | 56.6 | | | 56.5 | |
| | 2-NH | | 8.20 (s) | | | 8.20 (s) |
| | 3 | 26.1~27.0 | ~1.46 (o) | | 26.2~26.9 | ~1.46 (o) |
| | 4 | 23.2~23.6 | ~1.35 (o) | | 23.2~23.6 | ~1.35 (o) |
| **Pro$^{13}$** | 1 | 174.0 | | **Pro$^{13}$** | 173.8 | |
| | 2 | 63.0 | 4.30 (o) | | 63.1 | 4.29 (o) |
| | 3 | 32.1 | 2.22 (o) | | 32.1 | 2.23 (o) |
| | 4 | 29.1 | 1.62 (o) | | 29.5 | 1.69 (o) |
| | 5 | 49.0 | 3.73 (o), 3.55 (o) | | 49.0 | 3.72 (o), 3.58(o) |
| **Val$^{14}$** | 1 | 175.8 | | **Val$^{14}$** | 161.575.7 | |
| | 2 | 61.6 | 3.77 (o) | | | 3.79 (o) |
| | 2-NH | | 7.59 (o) | | | 7.52 (o) |
| | 3 | 29.1 | 2.25 (o) | | 29.1 | 2.24 (o) |
| | 4 | 19.4 | 0.89 (o) | | 19.4 | 0.88 (o) |
| | 5 | 19.5 | 0.98 (d, 6.4) | | 19.5 | 0.95 (d, 6.4) |

o, overlapping signals

**Table S5.** $^1$H and $^{13}$C NMR data of trichohypolins A (**1**) and B (**2**) in DMSO-*d*$_6$ (continued)

| | | trichohypolin A (**1**) | | | trichohypolin B (**2**) | |
|---|---|---|---|---|---|---|
| residue | position | $\delta_C$$^a$, mult. | $\delta_H$$^b$ (*J* in Hz) | residue | $\delta_C$$^a$, mult. | $\delta_H$$^b$ (*J* in Hz) |
| **Aib$^{15}$** | 1 | 174.2 | | **Aib$^{15}$** | 174.0 | |
| | 2 | 56.0 | | | 56.5 | |
| | 2-NH | | 7.58 (o) | | | 7.64 (o) |
| | 3 | 26.1~27.0 | ~1.46 (o) | | 26.2~26.9 | ~1.46 (o) |
| | 4 | 23.2~23.6 | ~1.35 (o) | | 23.2~23.6 | ~1.35 (o) |
| **Aib$^{16}$** | 1 | 175.9 | | **Iva$^{16}$** | 173.8 | |
| | 2 | 56.1 | | | 59.2 | |
| | 2-NH | | 7.74 (o) | | | 7.51 (s) |
| | 3 | 26.1~27.0 | ~1.46 (o) | | 26.5 | 1.44 (o) |
| | 4 | 23.2~23.6 | ~1.35 (o) | | 29.5 | 1.69 (m), 2.15 (m) |
| | | | | | 7.59 | 0.71 (t, 7.25) |
| **Gln$^{17}$** | 1 | 172.5 | | **Gln$^{17}$** | 172.5 | |
| | 2 | 55.2 | 3.83 (o) | | 56.3 | 3.83 (o) |
| | 2-NH | | 7.93 (brs) | | | 7.89 (o) |
| | 3 | 27.1 | 1.98 (o) 2.09 (o) | | 26.3 | 1.96 (o), 2.01 (o) |
| | 4 | 31.0 | 2.23 (o), 2.21 (o) | | 31.6 | 2.23 (o), 2.10 (o) |
| | 5 | 174.1 | | | 173.5 | |
| | 5-NH2 | | 6.77 (brs), 7.17 (brs) | | | 6.77 (brs), 7.16 (brs) |
| **Gln$^{18}$** | 1 | 171.4 | | **Gln$^{18}$** | 171.5 | |
| | 2 | 54.0 | 3.97 (o) | | 54.0 | 3.95 (o) |
| | 2-NH | | 7.51 (o) | | | 7.51 (o) |
| | 3 | 27.2 | 1.77 (o), 1.87 (o) | | 27.2 | 1.77 (o), 1.87 (o) |
| | 4 | 32.0 | 1.99 (o), 2.10 (o) | | 32.1 | 1.99 (o), 2.10 (o) |
| | 5 | 173.8 | | | 173.4 | |
| | 5-NH2 | | 6.62 (brs), 7.06 (brs) | | | 6.62 (brs), 7.06 (brs) |
| **Pheol$^{19}$** | 1 | 63.4 | 3.34 (o) | **Pheol$^{19}$** | 63.3 | 3.33 (o) |
| | 2 | 52.8 | 3.89 (m) | | 52.6 | 3.92 (o) |
| | 2-NH | | 6.96 (d, 8.9) | | | 6.92 (d, 8.70) |
| | 3 | 37.2 | 2.91 (dd, 4.96, 13.8) | | 37.0 | 2.92 (dd, 4.92, 14.0) |
| | | | 2.55 (dd, 8.96, 13.8) | | | 2.55 (dd, 9.13, 14.0) |
| | 4 | 139.5 | | | 139.5 | |
| | 5, 9 | 128.3 | 7.20 (m) | | 128.4 | 7.18-7.22 (m) |
| | 6, 8 | 129.6 | 7.23 (m) | | 129.6 | 7.18-7.22 (m) |
| | 7 | 126.3 | 7.13 (m) | | 126.2 | 7.14 (m) |

o, overlapping signals

**Table S6. Calculated mass of diagnostic fragment ions [*m/z*] of trichohypolins produced by *T. hypoxylon***

| **Comp.** | $b_{12}^+$ | $b_{11}^+$ | $b_{10}^+$ | $b_9^+$ | $b_8^+$ | $b_7^+$ | $b_6^+$ | $b_5^+$ | $b_4^+$ | $b_3^+$ | $b_2^+$ | $y_7^+$ | $y_7^+$-aa$^{19}$ | $y_7^+$-aa$^{18-19}$ | $y_7^+$-aa$^{17-19}$ | $y_7^+$-aa$^{16-19}$ | $y_7^+$-aa$^{15-19}$ |
|---|---|---|---|---|---|---|---|---|---|---|---|---|---|---|---|---|---|
| **1** | 1078.6255 | 993.5728 | 880.4887 | 823.4672 | 738.4145 | 639.3461 | 554.2933 | 426.2347 | 355.1976 | 284.1605 | 199.1077 | 774.4509 | 623.3511 | 495.2926 | 367.2340 | 282.1812 | 197.1285 |
| **2** | 1078.6255 | 993.5728 | 880.4887 | 823.4672 | 738.4145 | 639.3461 | 554.2933 | 426.2347 | 355.1976 | 284.1605 | 199.1077 | 788.4665 | 637.3668 | 509.3082 | 381.2496 | 282.1812 | 197.1285 |
| **3** | 1064.6099 | 979.5571 | 866.4730 | 809.4516 | 724.3988 | 625.3304 | 540.2776 | 412.2191 | 341.1819 | 284.1605 | 199.1077 | 774.4509 | 623.3511 | 495.2926 | 367.2340 | 282.1812 | 197.1285 |
| **4** | 1092.6412 | 1007.5884 | 894.5043 | 837.4829 | 752.4301 | 639.3461 | 554.2933 | 426.2347 | 355.1976 | 284.1605 | 199.1077 | 774.4509 | 623.3511 | 495.2926 | 367.2340 | 282.1812 | 197.1285 |
| **5** | 1064.6099 | 979.5571 | 866.4730 | 809.4516 | 724.3988 | 625.3304 | 540.2776 | 412.2191 | 341.1819 | 284.1605 | 199.1077 | 788.4665 | 637.3668 | 509.3082 | 381.2496 | 282.1812 | 197.1285 |
| **6** | 1064.6099 | 979.5571 | 866.4730 | 809.4516 | 724.3988 | 625.3304 | 540.2776 | 412.2191 | 341.1819 | 284.1605 | 199.1077 | 775.4349 | 624.3352 | 496.2766 | 367.2340 | 282.1812 | 197.1285 |
| **7** | 1064.6099 | 979.5571 | 866.4730 | 809.4516 | 724.3988 | 625.3304 | 540.2776 | 412.2191 | 341.1819 | 284.1605 | 199.1077 | 816.4614 | 623.3511 | 495.2926 | 367.2340 | 282.1812 | 197.1285 |
| **8** | 1093.6252 | 1008.5724 | 895.4884 | 838.4669 | 753.4141 | 654.3457 | 569.2930 | 426.2347 | 355.1976 | 284.1605 | 199.1077 | 788.4665 | 637.3668 | 509.3082 | 381.2496 | 282.1812 | 197.1285 |
| **9** | 1078.6255 | 993.5728 | 880.4887 | 823.4672 | 738.4145 | 639.3461 | 554.2933 | 426.2347 | 355.1976 | 284.1605 | 199.1077 | 775.4349 | 624.3352 | 496.2766 | 367.2340 | 282.1812 | 197.1285 |
| **10** | 1078.6255 | 993.5728 | 880.4887 | 823.4672 | 738.4145 | 639.3461 | 554.2933 | 426.2347 | 355.1976 | 284.1605 | 199.1077 | 804.4614 | 637.3668 | 509.3082 | 381.2496 | 282.1812 | 197.1285 |
| **11** | 1092.6412 | 1007.5884 | 894.5043 | 837.4829 | 752.4301 | 639.3461 | 554.2933 | 426.2347 | 355.1976 | 284.1605 | 199.1077 | 788.4665 | 637.3668 | 509.3082 | 381.2496 | 282.1812 | 197.1285 |
| **12** | 1064.6099 | 979.5571 | 866.4730 | 809.4516 | 724.3988 | 625.3304 | 540.2776 | 412.2191 | 341.1819 | 284.1605 | 199.1077 | 830.4771 | 637.3668 | 509.3082 | 381.2496 | 282.1812 | 197.1285 |
| **13** | 1064.6099 | 979.5571 | 880.4887 | 823.4672 | 738.4145 | 639.3461 | 511.2875 | 426.2347 | 355.1976 | 284.1605 | 199.1077 | 774.4509 | 623.3511 | 495.2926 | 367.2340 | 282.1812 | 197.1285 |
| **14** | 1078.6255 | 993.5728 | 880.4887 | 823.4672 | 738.4145 | 639.3461 | 554.2933 | 426.2347 | 355.1976 | 284.1605 | 199.1077 | 816.4614 | 623.3511 | 495.2926 | 367.2340 | 282.1812 | 197.1285 |
| **15** | 1092.6412 | 1007.5884 | 894.5043 | 837.4829 | 752.4301 | 653.3617 | 568.3089 | 440.2504 | 355.1976 | 284.1605 | 199.1077 | 774.4509 | 623.3511 | 495.2926 | 367.2340 | 282.1812 | 197.1285 |
| **16** | 1092.6412 | 1007.5884 | 894.5043 | 837.4829 | 752.4301 | 653.3617 | 568.3089 | 440.2504 | 355.1976 | 284.1605 | 199.1077 | 788.4665 | 637.3668 | 509.3082 | 381.2496 | 282.1812 | 197.1285 |
| **17** | 1078.6255 | 993.5728 | 880.4887 | 823.4672 | 738.4145 | 639.3461 | 554.2933 | 426.2347 | 355.1976 | 284.1605 | 199.1077 | 789.4505 | 638.3508 | 510.2922 | 367.2340 | 282.1812 | 197.1285 |
| **18** | 1078.6255 | 993.5728 | 880.4887 | 823.4672 | 738.4145 | 639.3461 | 554.2933 | 426.2347 | 355.1976 | 284.1605 | 199.1077 | 789.4505 | 638.3508 | 510.2922 | 381.2496 | 282.1812 | 197.1285 |
| **19** | 1093.6252 | 1008.5724 | 895.4884 | 838.4669 | 753.4141 | 654.3457 | 569.2930 | 426.2347 | 355.1976 | 284.1605 | 199.1077 | 789.4505 | 638.3508 | 510.2922 | 367.2340 | 282.1812 | 197.1285 |
| **20** | 1093.6252 | 1008.5724 | 895.4884 | 838.4669 | 753.4141 | 654.3457 | 569.2930 | 426.2347 | 355.1976 | 284.1605 | 199.1077 | 830.4771 | 637.3668 | 509.3082 | 381.2496 | 282.1812 | 197.1285 |
| **21** | 1092.6412 | 1007.5884 | 894.5043 | 837.4829 | 752.4301 | 653.3617 | 568.3089 | 440.2504 | 355.1976 | 284.1605 | 199.1077 | 803.4662 | 652.3665 | 524.3079 | 381.2496 | 282.1812 | 197.1285 |

### Table S7. Calculated mass of diagnostic fragment ions [*m/z*] of trichohypolins produced by *T. hypoxylon* (continued)

| Comp. | $b_{12}^+$ | $b_{11}^+$ | $b_{10}^+$ | $b_9^+$ | $b_8^+$ | $b_7^+$ | $b_6^+$ | $b_5^+$ | $b_4^+$ | $b_3^+$ | $b_2^+$ | $y_7^+$ | $y_7^+$-aa$^{19}$ | $y_7^+$-aa$^{18-19}$ | $y_7^+$-aa$^{17-19}$ | $y_7^+$-aa$^{16-19}$ | $y_7^+$-aa$^{15-19}$ |
|---|---|---|---|---|---|---|---|---|---|---|---|---|---|---|---|---|---|
| **22** | 1078.6255 | 993.5728 | 880.4887 | 823.4672 | 738.4145 | 639.3461 | 554.2933 | 426.2347 | 355.1976 | 284.1605 | 199.1077 | 803.4662 | 652.3665 | 524.3079 | 381.2496 | 282.1812 | 197.1285 |
| **23** | 1078.6255 | 993.5728 | 880.4887 | 823.4672 | 738.4145 | 639.3461 | 554.2933 | 426.2347 | 355.1976 | 284.1605 | 199.1077 | 817.4454 | 624.3352 | 496.2766 | 367.2340 | 282.1812 | 197.1285 |
| **24** | 1078.6255 | 993.5728 | 880.4887 | 823.4672 | 738.4145 | 639.3461 | 554.2933 | 426.2347 | 355.1976 | 284.1605 | 199.1077 | 830.4771 | 637.3668 | 509.3082 | 381.2496 | 282.1812 | 197.1285 |
| **25** | 1079.6095 | 994.5568 | 881.4727 | 824.4512 | 739.3985 | 640.3301 | 555.2773 | 426.2347 | 355.1976 | 284.1605 | 199.1077 | 775.4349 | 624.3352 | 496.2766 | 367.2340 | 282.1812 | 197.1285 |
| **26** | 1092.6412 | 1007.5884 | 894.5043 | 837.4829 | 752.4301 | 639.3461 | 554.2933 | 426.2347 | 355.1976 | 284.1605 | 199.1077 | 830.4771 | 637.3668 | 509.3082 | 381.2496 | 282.1812 | 197.1285 |
| **27** | 1078.6255 | 993.5728 | 880.4887 | 823.4672 | 738.4145 | 639.3461 | 554.2933 | 426.2347 | 355.1976 | 284.1605 | 199.1077 | 817.4454 | 624.3352 | 495.2926 | 367.2340 | 282.1812 | 197.1285 |
| **28** | 1079.6095 | 994.5568 | 881.4727 | 824.4512 | 739.3985 | 640.3301 | 555.2773 | 426.2347 | 355.1976 | 284.1605 | 199.1077 | 788.4665 | 637.3668 | 509.3082 | 381.2496 | 282.1812 | 197.1285 |
| **29** | 1092.6412 | 1007.5884 | 894.5043 | 837.4829 | 752.4301 | 639.3461 | 554.2933 | 426.2347 | 355.1976 | 284.1605 | 199.1077 | 775.4349 | 624.3352 | 496.2766 | 367.2340 | 282.1812 | 197.1285 |
| **30** | 1064.6099 | 979.5571 | 866.4730 | 809.4516 | 738.4145 | 639.3461 | 554.2933 | 426.2347 | 355.1976 | 284.1605 | 199.1077 | 760.4352 | 609.3355 | 481.2769 | 353.2183 | 254.1499 | 169.0972 |
| **31** | 1106.6568 | 1021.6041 | 908.5200 | 851.4985 | 766.4458 | 667.3774 | 582.3246 | 454.2660 | 383.2289 | 312.1918 | 227.1390 | 774.4509 | 623.3511 | 495.2926 | 367.2340 | 282.1812 | 197.1285 |
| **32** | 1106.6568 | 1021.6041 | 908.5200 | 851.4985 | 766.4458 | 667.3774 | 582.3246 | 454.2660 | 383.2289 | 312.1918 | 227.1390 | 789.4505 | 638.3508 | 510.2922 | 367.2340 | 282.1812 | 197.1285 |
| **33** | 1093.6252 | 1008.5724 | 895.4884 | 838.4669 | 753.4141 | 654.3457 | 569.2930 | 426.2347 | 355.1976 | 284.1605 | 199.1077 | 774.4509 | 623.3511 | 495.2926 | 367.2340 | 282.1812 | 197.1285 |
| **34** | 1079.6095 | 994.5568 | 881.4727 | 824.4512 | 739.3985 | 640.3301 | 555.2773 | 426.2347 | 355.1976 | 284.1605 | 199.1077 | 789.4505 | 638.3508 | 510.2922 | 367.2340 | 282.1812 | 197.1285 |
| **35** | 1079.6095 | 994.5568 | 881.4727 | 824.4512 | 739.3985 | 640.3301 | 555.2773 | 426.2347 | 355.1976 | 284.1605 | 199.1077 | 816.4614 | 623.3511 | 495.2926 | 367.2340 | 282.1812 | 197.1285 |
| **36** | 1078.6255 | 993.5728 | 880.4887 | 823.4672 | 738.4145 | 639.3461 | 554.2933 | 426.2347 | 355.1976 | 284.1605 | 199.1077 | 831.4611 | 638.3508 | 510.2922 | 367.2340 | 282.1812 | 197.1285 |
| **37** | 1078.6255 | 993.5728 | 880.4887 | 823.4672 | 738.4145 | 639.3461 | 554.2933 | 426.2347 | 355.1976 | 284.1605 | 199.1077 | 789.4505 | 638.3508 | 495.2926 | 367.2340 | 282.1812 | 197.1285 |
| **38** | 1093.6252 | 1008.5724 | 895.4884 | 838.4669 | 753.4141 | 654.3457 | 569.2930 | 426.2347 | 355.1976 | 284.1605 | 199.1077 | 789.4505 | 638.3508 | 495.2926 | 367.2340 | 282.1812 | 197.1285 |
| **39** | 1079.6095 | 994.5568 | 881.4727 | 824.4512 | 739.3985 | 640.3301 | 555.2773 | 426.2347 | 355.1976 | 284.1605 | 199.1077 | 774.4509 | 623.3511 | 495.2926 | 367.2340 | 282.1812 | 197.1285 |
| **40** | 1078.6255 | 993.5728 | 880.4887 | 823.4672 | 738.4145 | 639.3461 | 554.2933 | 426.2347 | 355.1976 | 284.1605 | 199.1077 | 790.4345 | 639.3348 | 510.2922 | 367.2340 | 282.1812 | 197.1285 |
| **41** | 1078.6255 | 993.5728 | 880.4887 | 823.4672 | 738.4145 | 639.3461 | 554.2933 | 426.2347 | 355.1976 | 284.1605 | 199.1077 | 831.4611 | 638.3508 | 510.2922 | 381.2496 | 282.1812 | 197.1285 |
| **42** | 1093.6252 | 1008.5724 | 895.4884 | 838.4669 | 753.4141 | 654.3457 | 569.2930 | 426.2347 | 355.1976 | 284.1605 | 199.1077 | 775.4349 | 624.3352 | 496.2766 | 367.2340 | 282.1812 | 197.1285 |

**Table S7.** Inhibitory effects by trichohypolins on saprotrophic fungi

| **Trichohypolins** | **Inhibitory effect ranking of saprotrophic fungi** |
|---|---|
| trichohypolin A (**1**) | PA > TE > KP > $XC_a$ > $XS_b$ > $XS_c$ > $XS_a$ > CB |
| trichohypolin B (**2**) | TE > PA > KP > $XC_a$ > $XS_a$> $XS_c$ > $XS_b$ > CB |
| trichohypolin SF1 (**1** and **3**–**8**) | PA > TE >KP > $XC_a$ > $XS_a$ > $XS_b$ = $XS_c$ > CB |
| trichohypolin SF2 (**2** and **9**–**13**) | TE > PA > KP > $XC_a$ > $XS_a$ > CB > $XS_c$ > $XS_b$ |
| trichohypolin SF3 (**14**–**21**) | PA > TE > $XS_a$ > KP > $XC_a$ > $XS_b$ > CB > $XS_c$ |
| trichohypolin SF4 (**22**–**32**) | TE > PA > KP > $XS_a$ > $XS_b$ > $XC_a$ > CB > $XS_c$ |
| trichohypolin SF5 (**33**–**42**) | PA > TE > KP > $XS_a$ > $XC_a$ > $XS_b$ > $XS_c$ > CB |

KP, *K. pavimentosa* H2535; $XS_a$, *X. schweinitzii* L1375; $XS_b$, *X. schweinitzii* LW3006; $XS_c$, *X. schweinitzii* L1374; CB, *C. byssicola* LW3015; $XC_a$, *X. cubensis* HG11; PA, *P. acerina* L1377; TE, *T. epimiltinus* L2613.

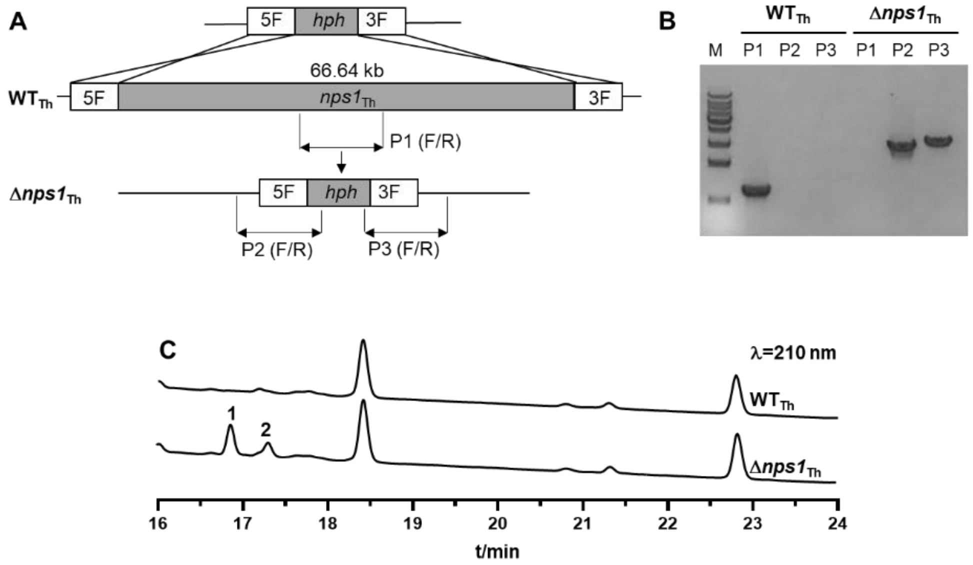

**Fig. S1.** Gene deletion of *nps1*$_{Th}$ in *T. hypoxylon* (A and B) as well as LCMS analysis of crude extracts of wild type and Δ*nps1*$_{Th}$.

UV absorptions at 210 nm are illustrated.

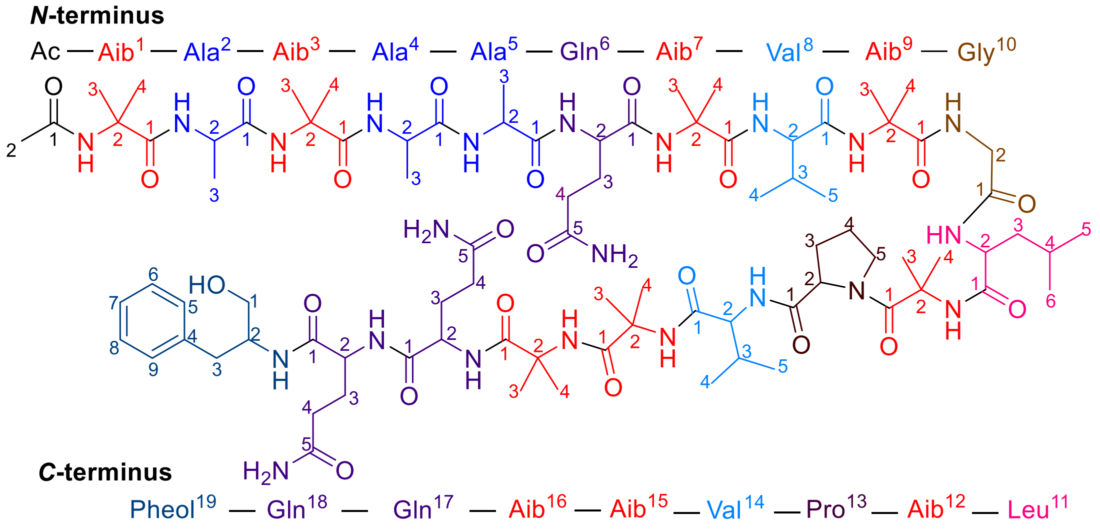

trichohypolin A (**1**)

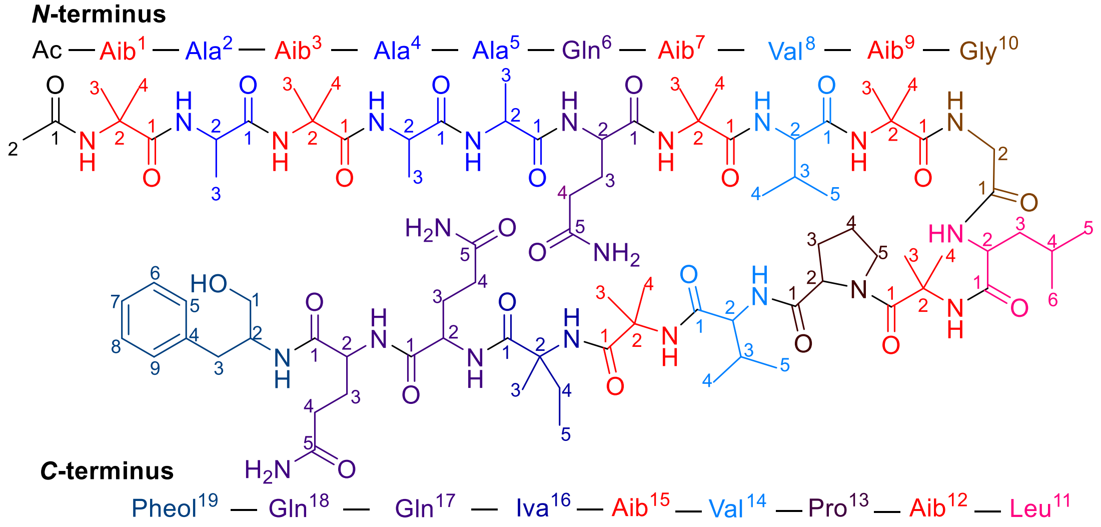

trichohypolin B (**2**)

**Fig. S2** Structures of trichohypolins A (**1**) and B (**2**)

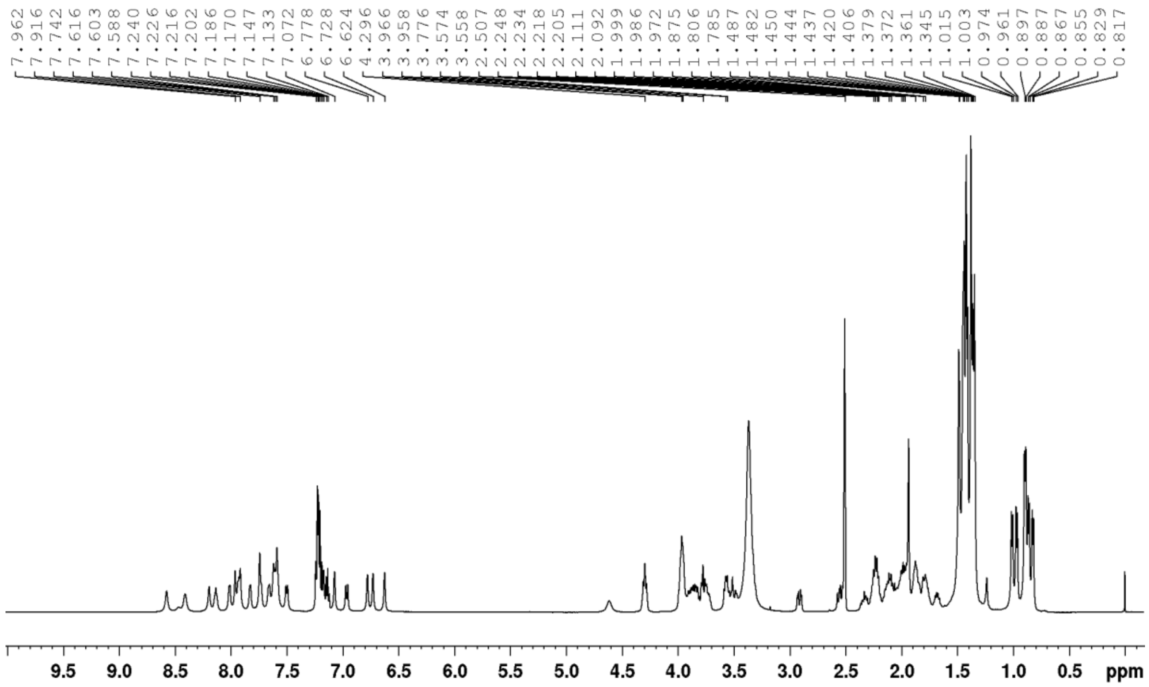

**Fig. S3** $^{1}$H NMR spectrum of trichohypolin A (**1**) in DMSO-$d_6$ (500 MHz)

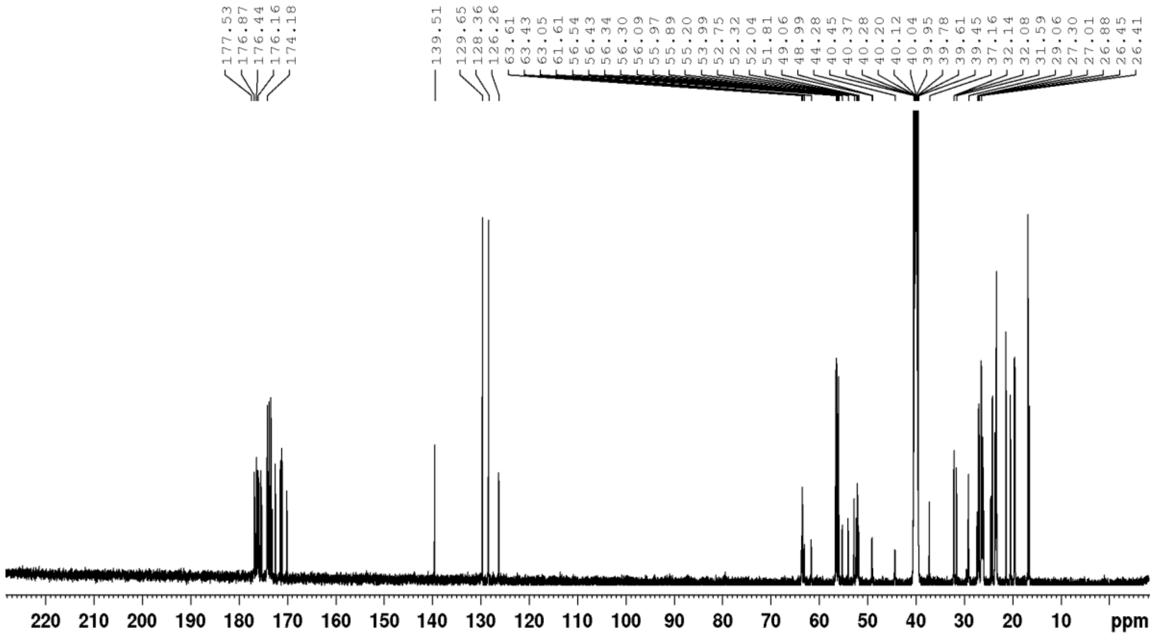

**Fig. S4** $^{13}$C NMR spectrum of trichohypolin A (**1**) in DMSO-$d_6$ (125 MHz)

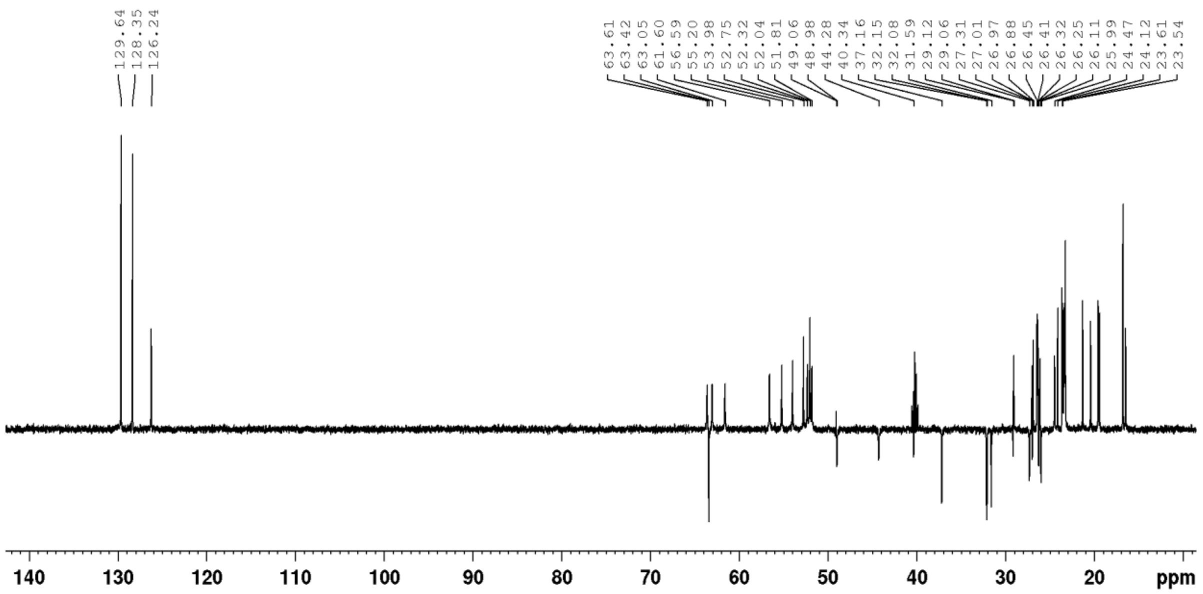

**Fig. S5** DEPT-135 NMR spectrum of trichohypolin A (**1**) in DMSO-*d6* (125 MHz)

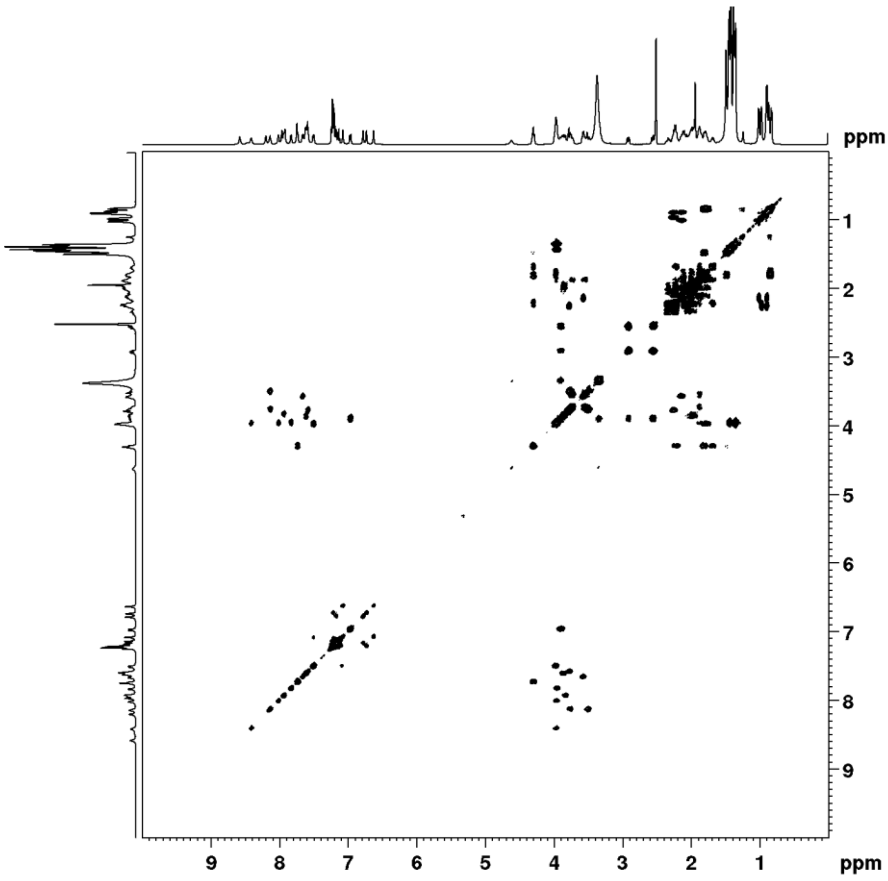

**Fig. S6** $^{1}$H-$^{1}$H COSY spectrum of trichohypolin A (**1**) in DMSO-*d6*

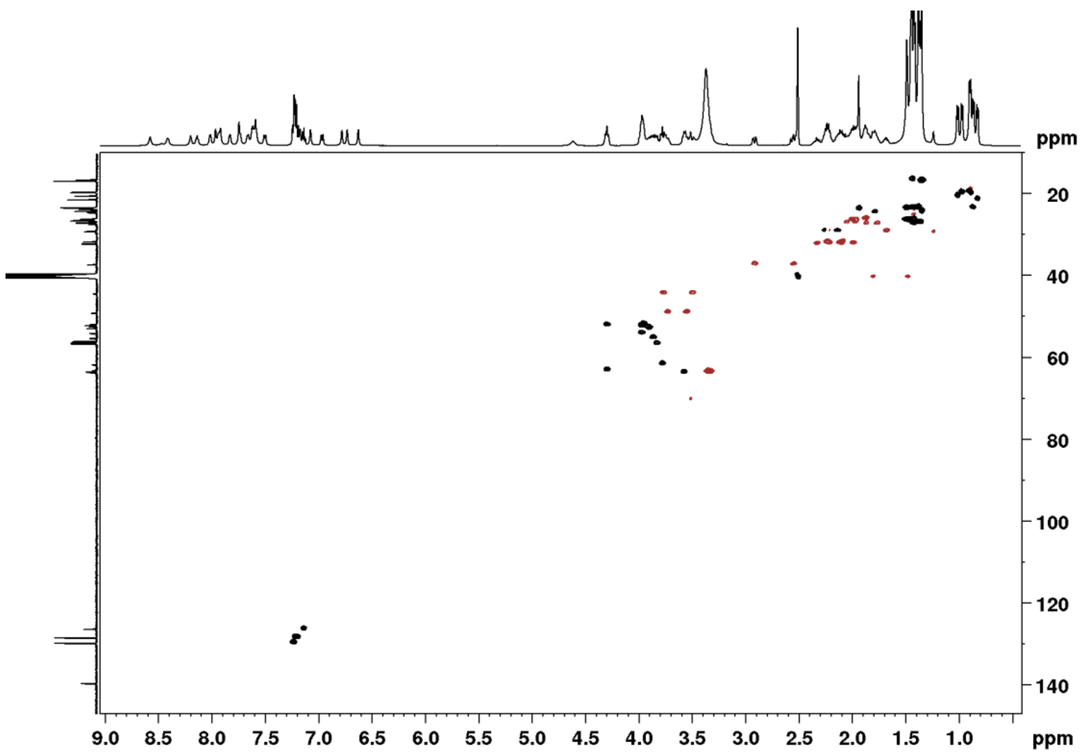

**Fig. S7** HSQC spectrum of trichohypolin A (**1**) in DMSO-$d_6$

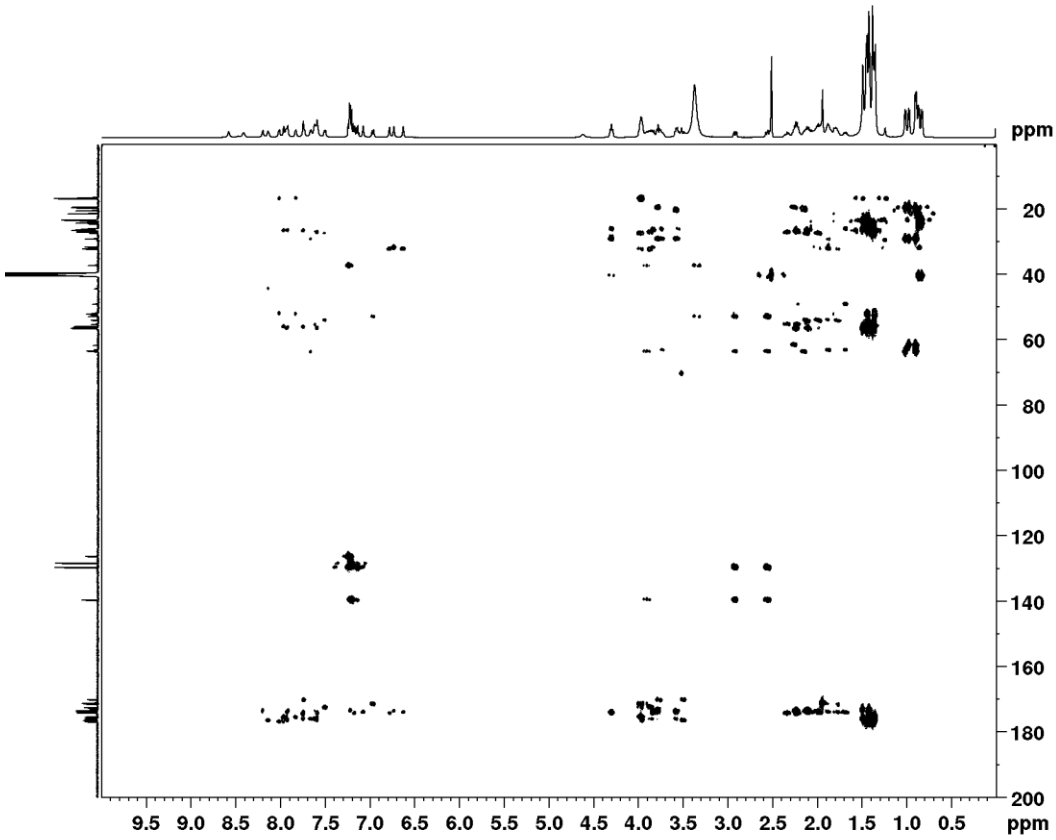

**Fig. S8** HMBC spectrum of trichohypolin A (**1**) in DMSO-$d_6$

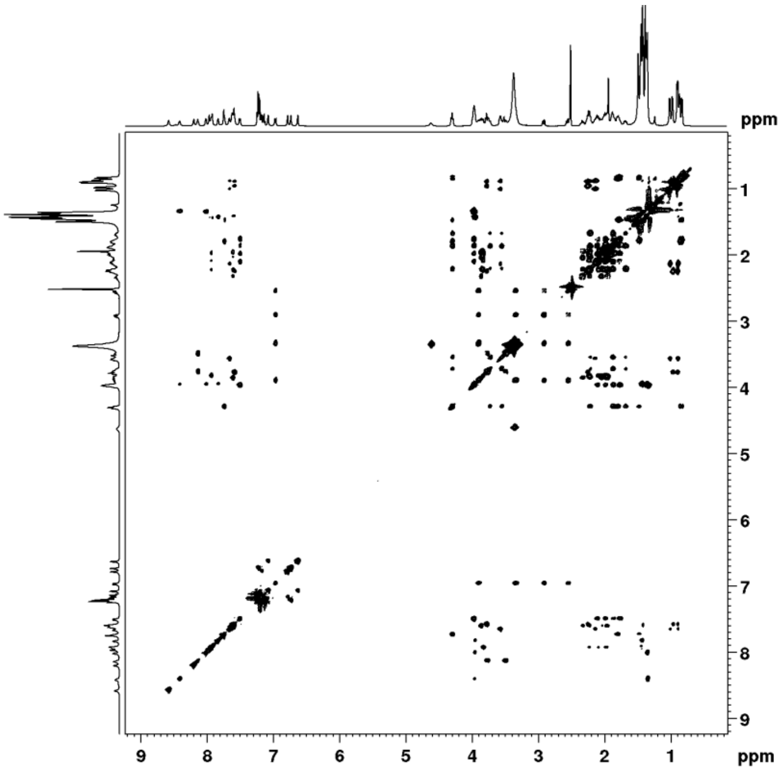

**Fig. S9** TOCSY spectrum of trichohypolin A (**1**) in DMSO-$d_6$

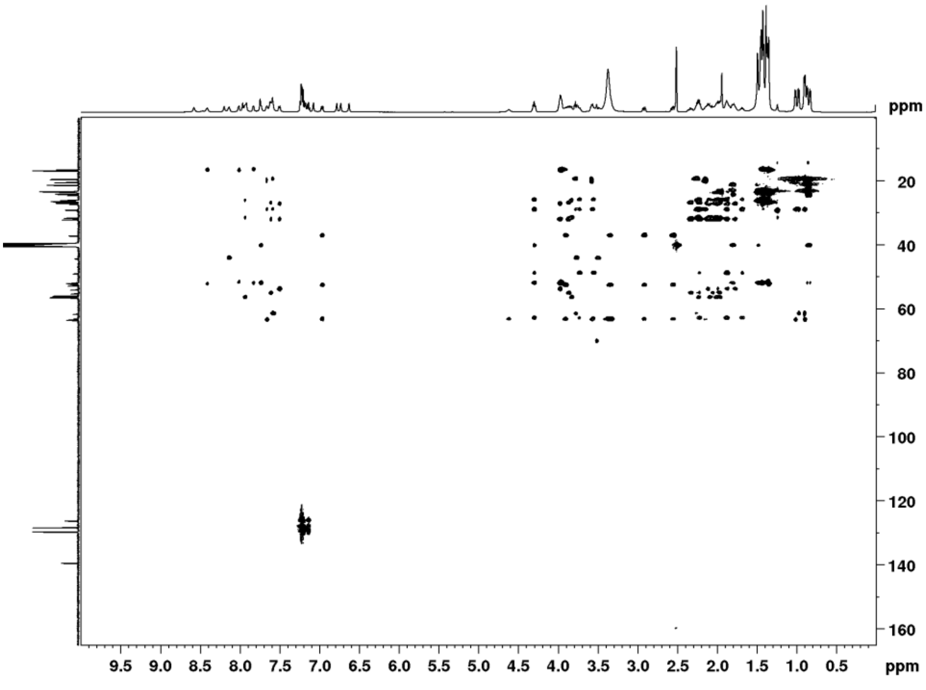

**Fig. S10** HSQC_TOCSY spectrum of trichohypolin A (**1**) in DMSO-$d_6$

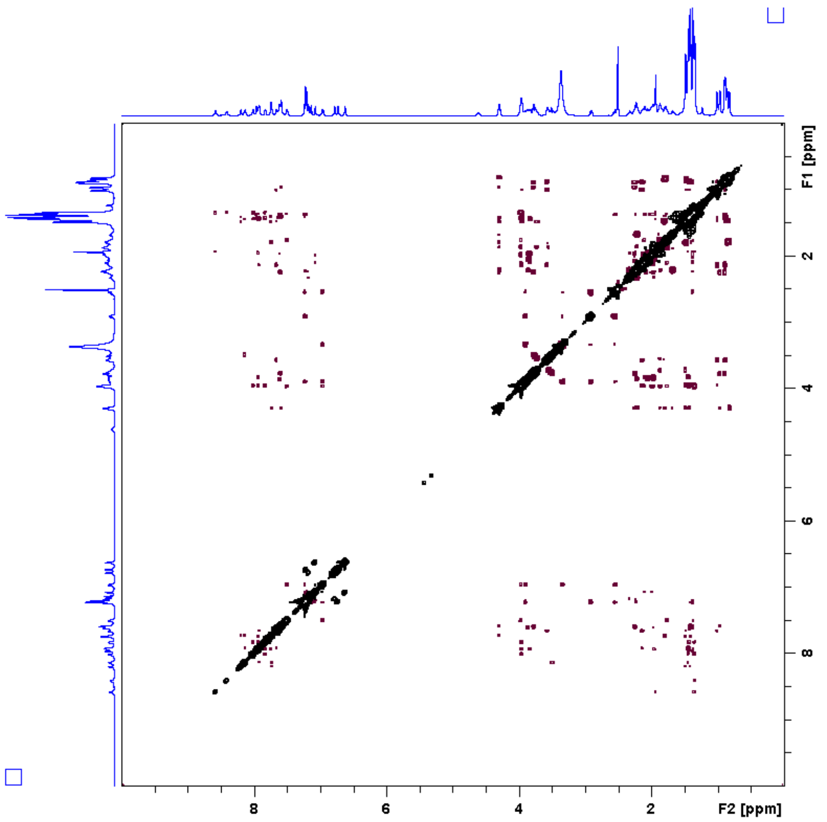

**Fig. S11** $^1$H-$^1$H ROESY spectrum of trichohypolin A (**1**) in DMSO-$d_6$

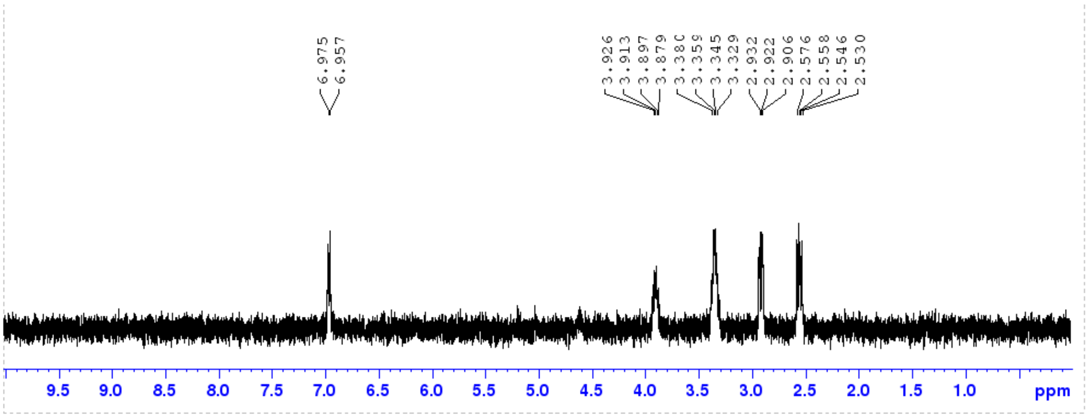

**Fig. S12 (A)** 1D TOCSY spectrum trichohypolin A (**1**) with selective excitation of 2-NH of pheol[19] (δ 6.96)

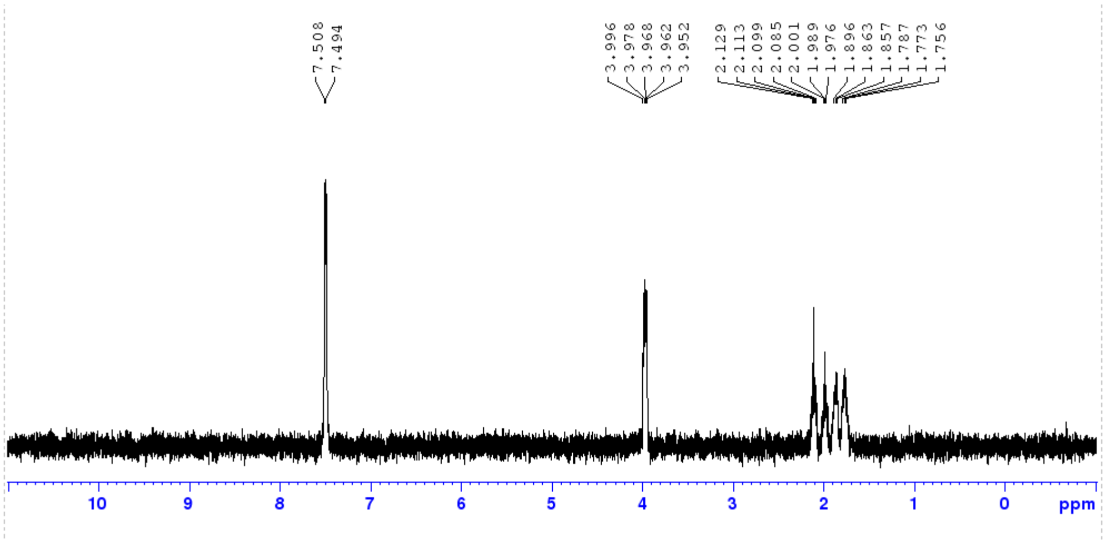

**Fig. S12 (B)** 1D TOCSY spectrum trichohypolin A (**1**) with selective excitation of 2-NH of Gln[18] (δ 7.51)

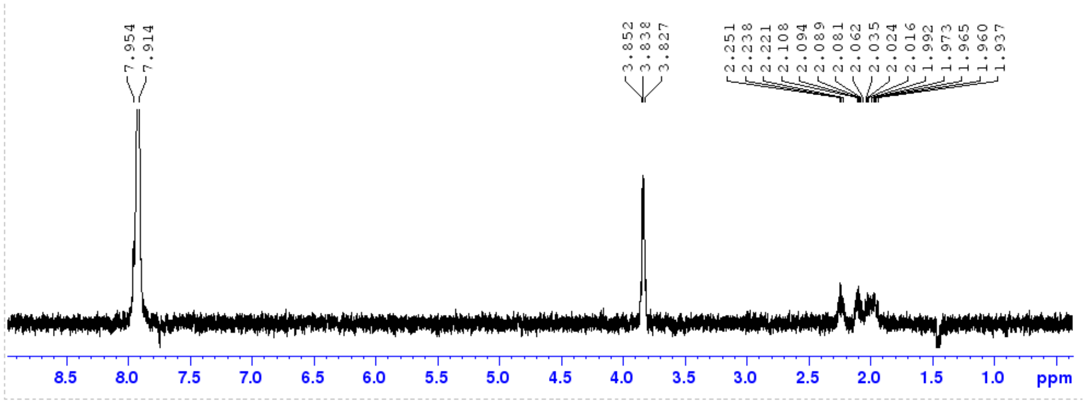

**Fig. S12 (C)** 1D TOCSY spectrum trichohypolin A (**1**) with selective excitation of 2-NH of Gln[17] (δ 7.93)

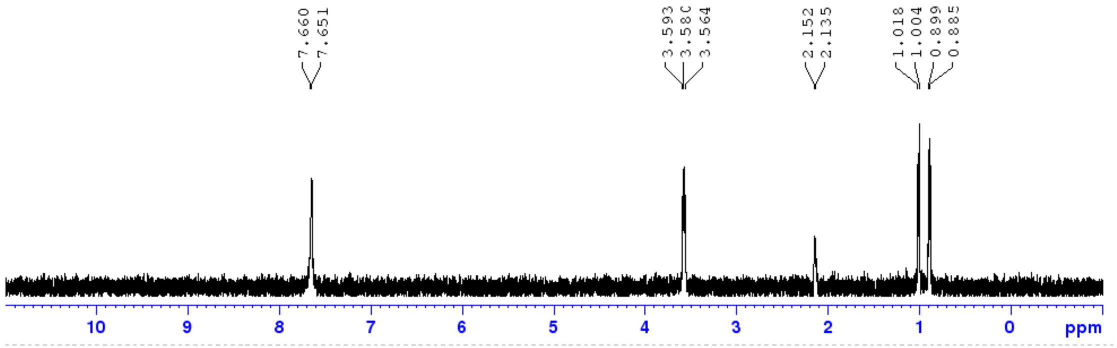

**Fig. S12 (D)** 1D TOCSY spectrum trichohypolin A (**1**) with selective excitation of 2-NH of Val[8] (δ 7.66)

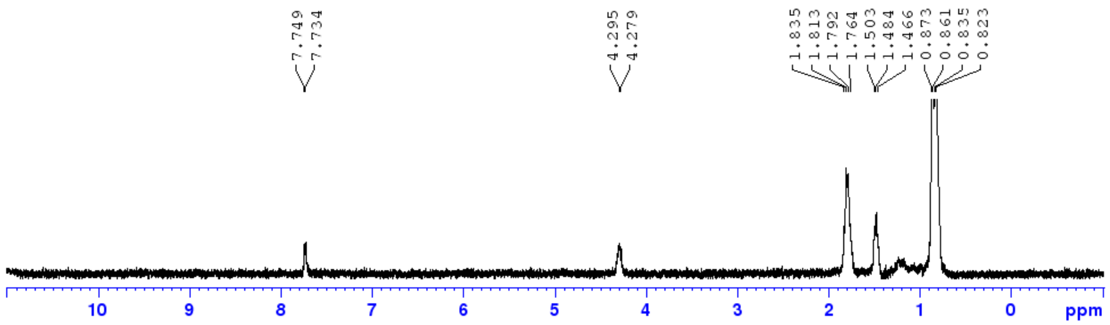

**Fig. S12 (E)** 1D TOCSY spectrum trichohypolin A (**1**) with selective excitation of 2-NH of $Leu^{11}$ (δ 7.74)

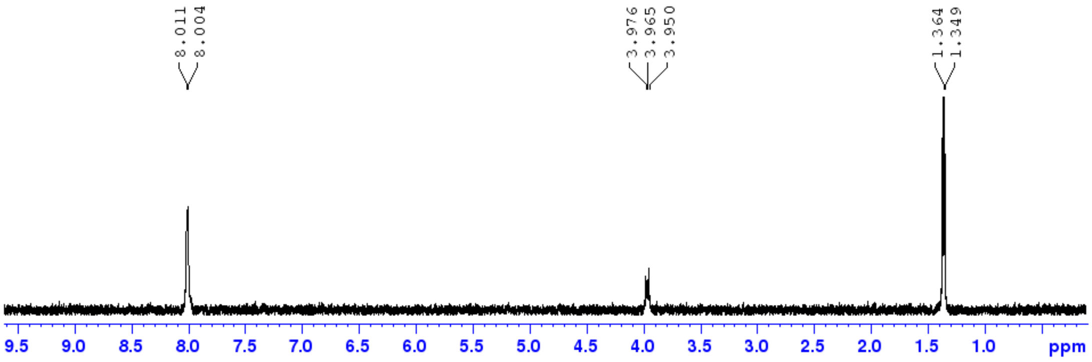

**Fig. S12 (F)** 1D TOCSY spectrum trichohypolin A (**1**) with selective excitation of 2-NH of $Ala^{4}$ (δ 8.01)

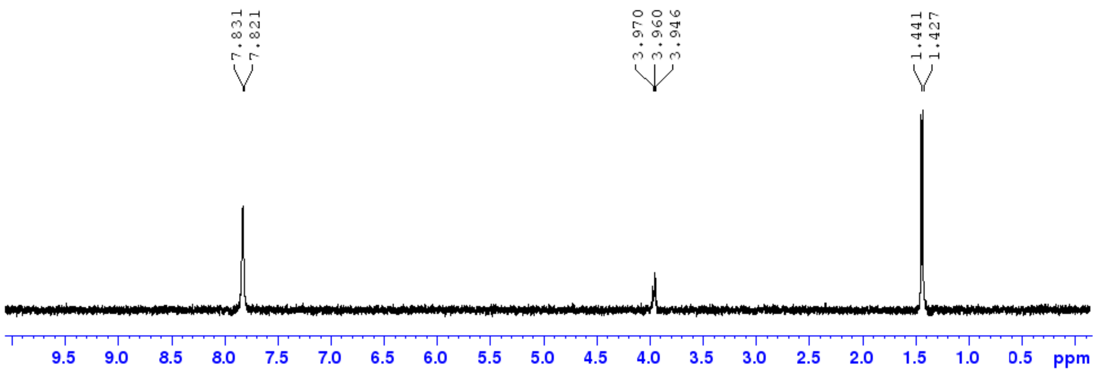

**Fig. S12 (G)** 1D TOCSY spectrum trichohypolin A (**1**) with selective excitation of 2-NH of $Ala^{5}$ (δ 7.83)

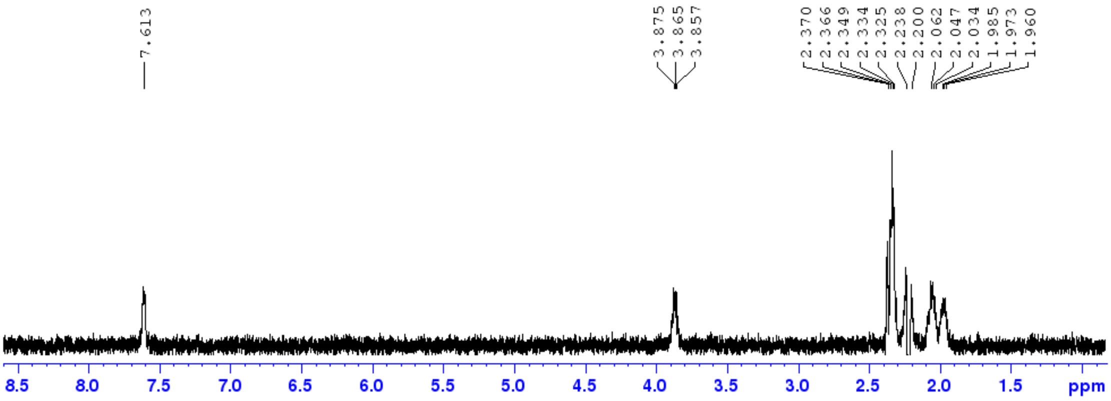

**Fig. S12 (H)** 1D TOCSY spectrum trichohypolin A (**1**) with selective excitation of 2-NH of Gln$^{6}$ (δ 7.61)

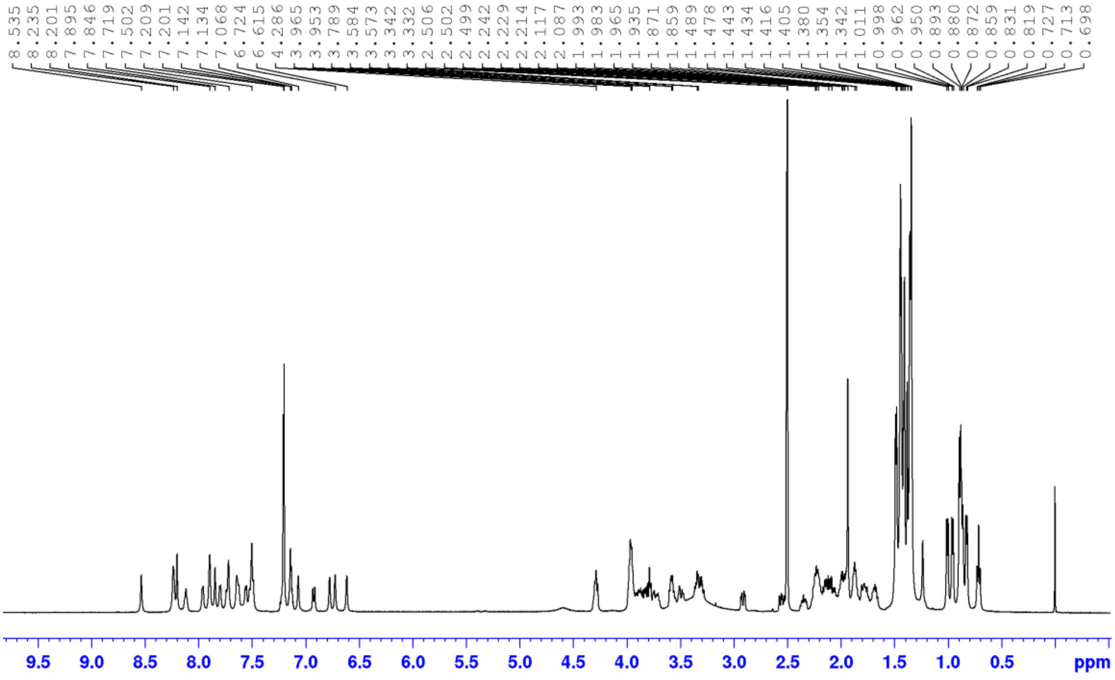

**Fig. S13** $^{1}H$ NMR spectrum of trichohypolin B (**2**) in DMSO-$d_6$ (500 MHz)

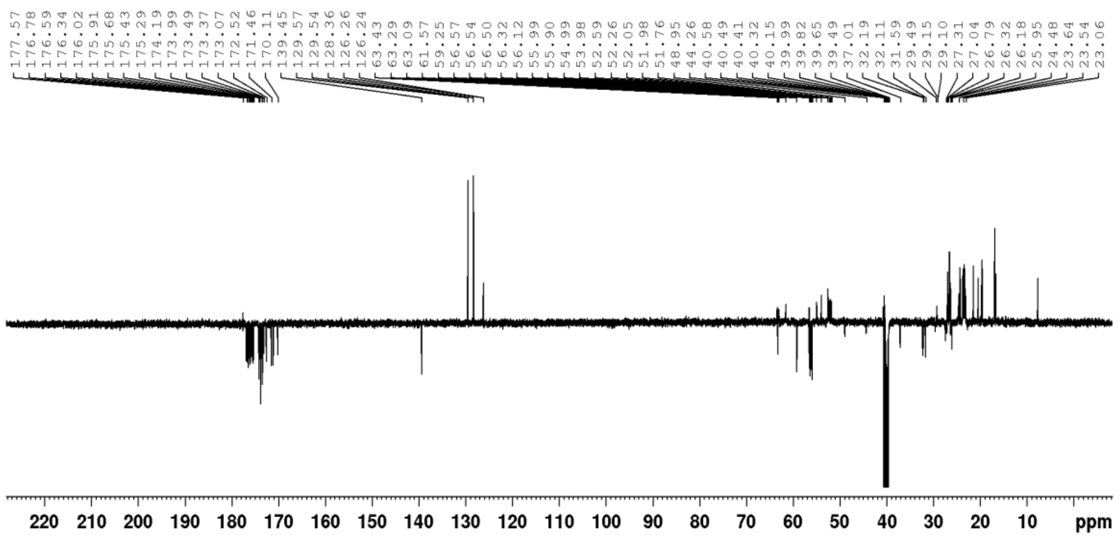

**Fig. S14** APT-$^{13}$C spectrum of trichohypolin B (**2**) in DMSO-*d*$_6$ (125 MHz)

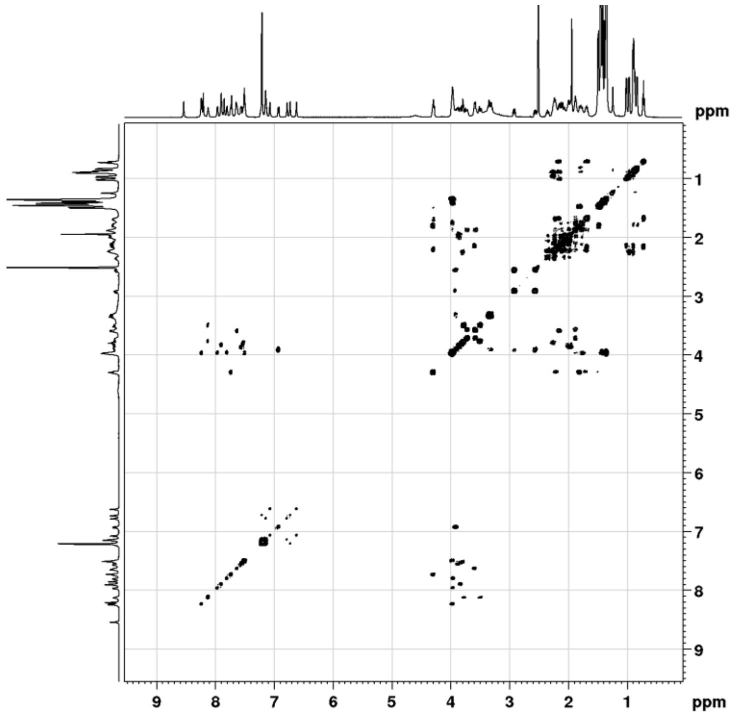

**Fig. S15** $^1$H-$^1$H COSY spectrum of trichohypolin B (**2**) in DMSO-*d*$_6$

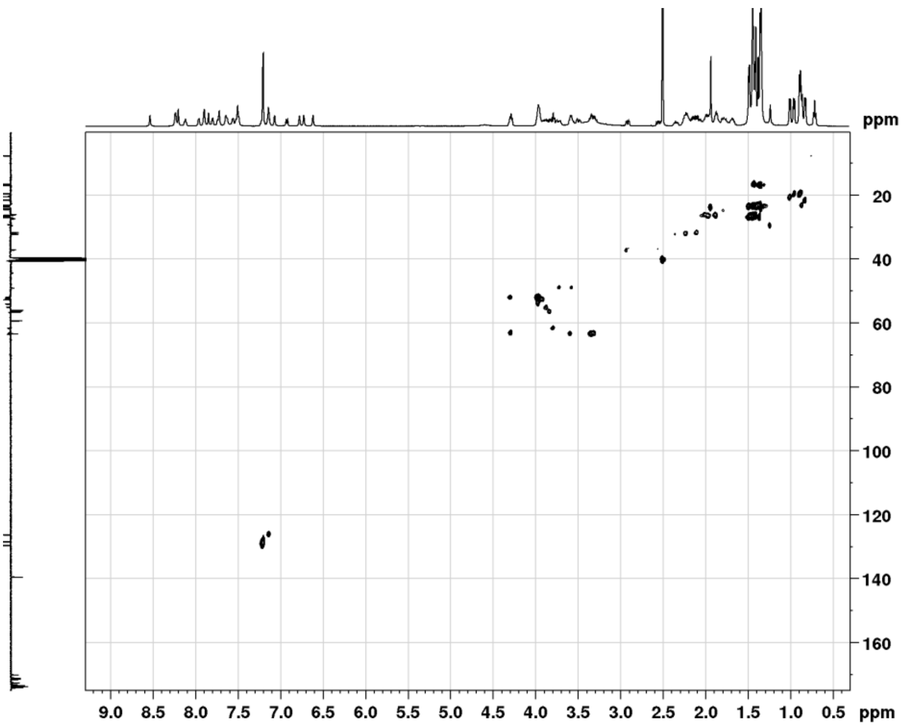

**Fig. S16** HSQC spectrum of trichohypolin B (**2**) in DMSO-$d_6$

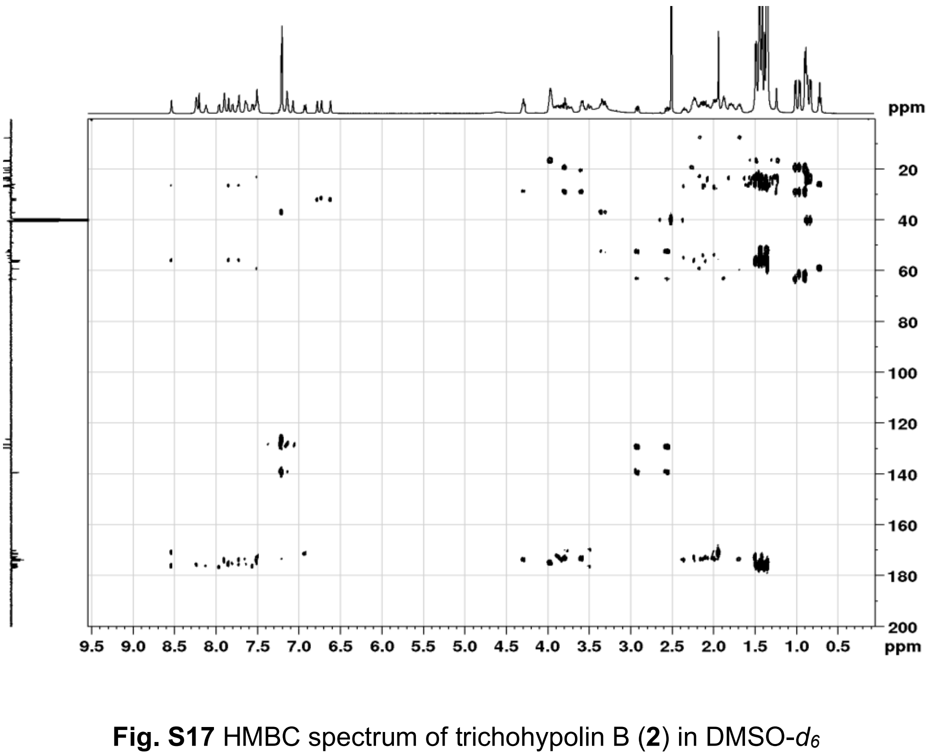

**Fig. S17** HMBC spectrum of trichohypolin B (**2**) in DMSO-$d_6$

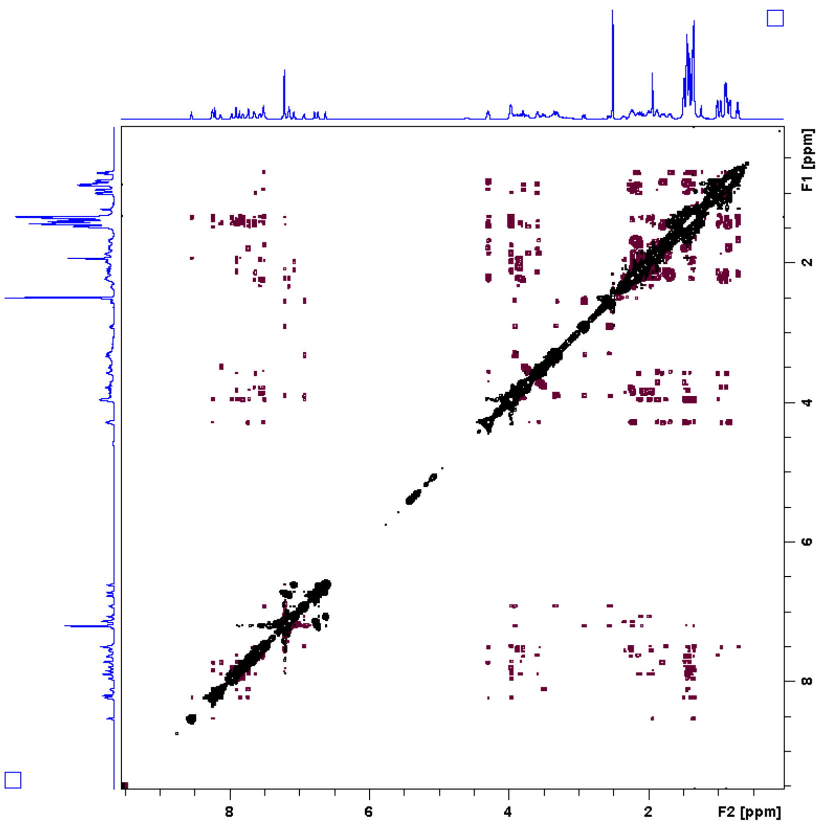

**Fig. S18** ROESY spectrum of trichohypolin B (**2**) in DMSO-$d_6$

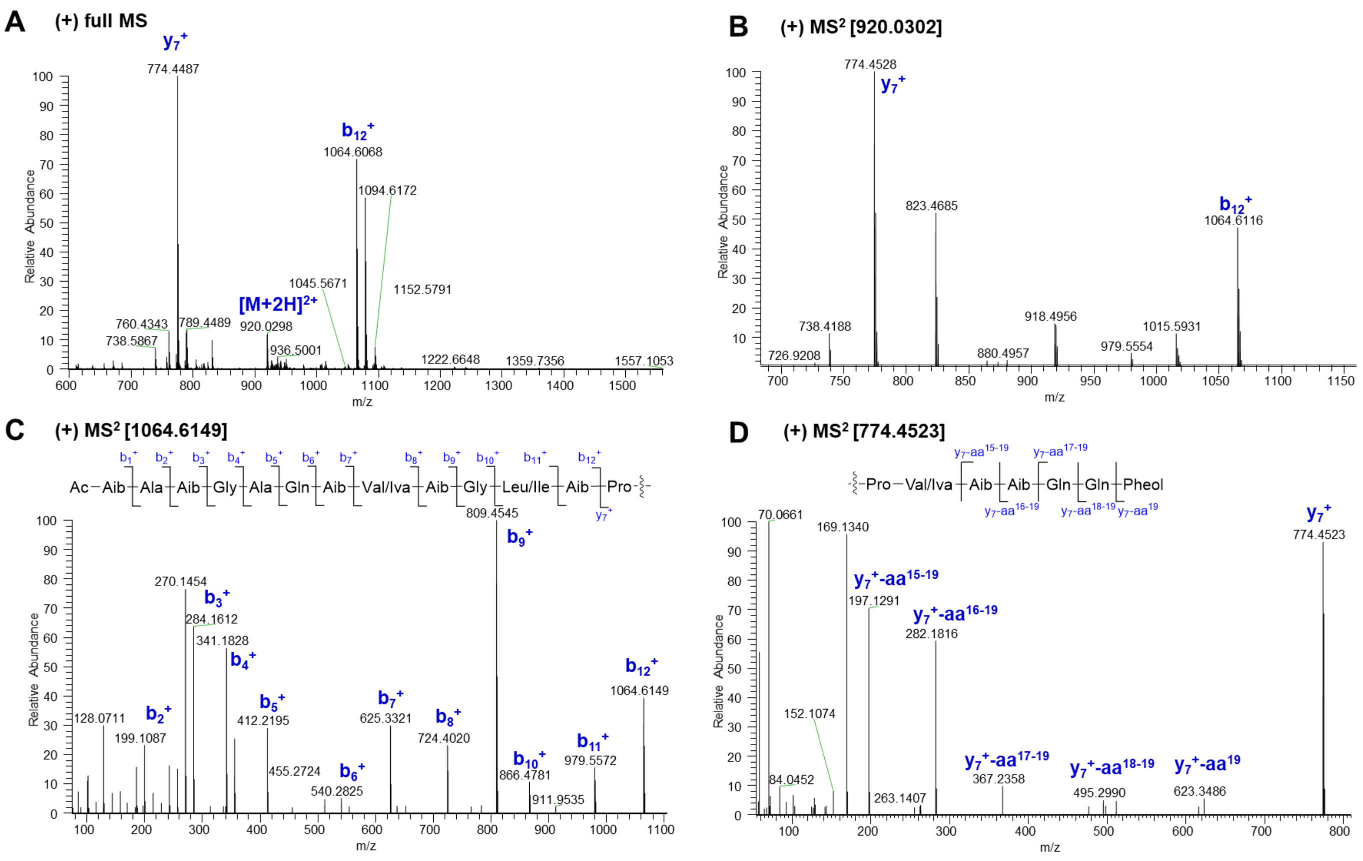

**Fig. S19** Diagnostic fragment ions [*m/z*] of **3** in subfraction 1 (SF1) by ESI-HRMS analysis

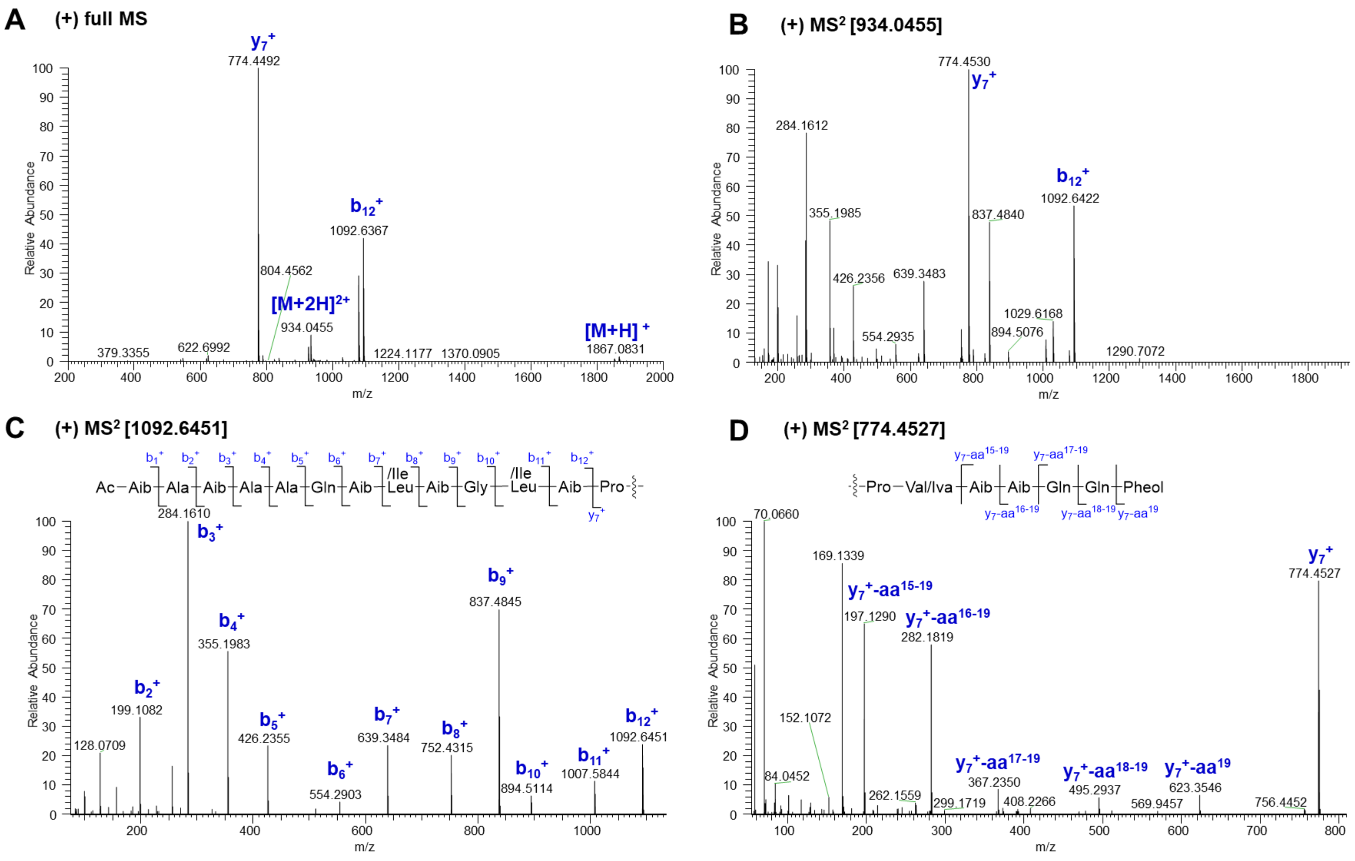

**Fig. S20** Diagnostic fragment ions [*m/z*] of **4** in subfraction 1 (SF1) by ESI-HRMS analysis

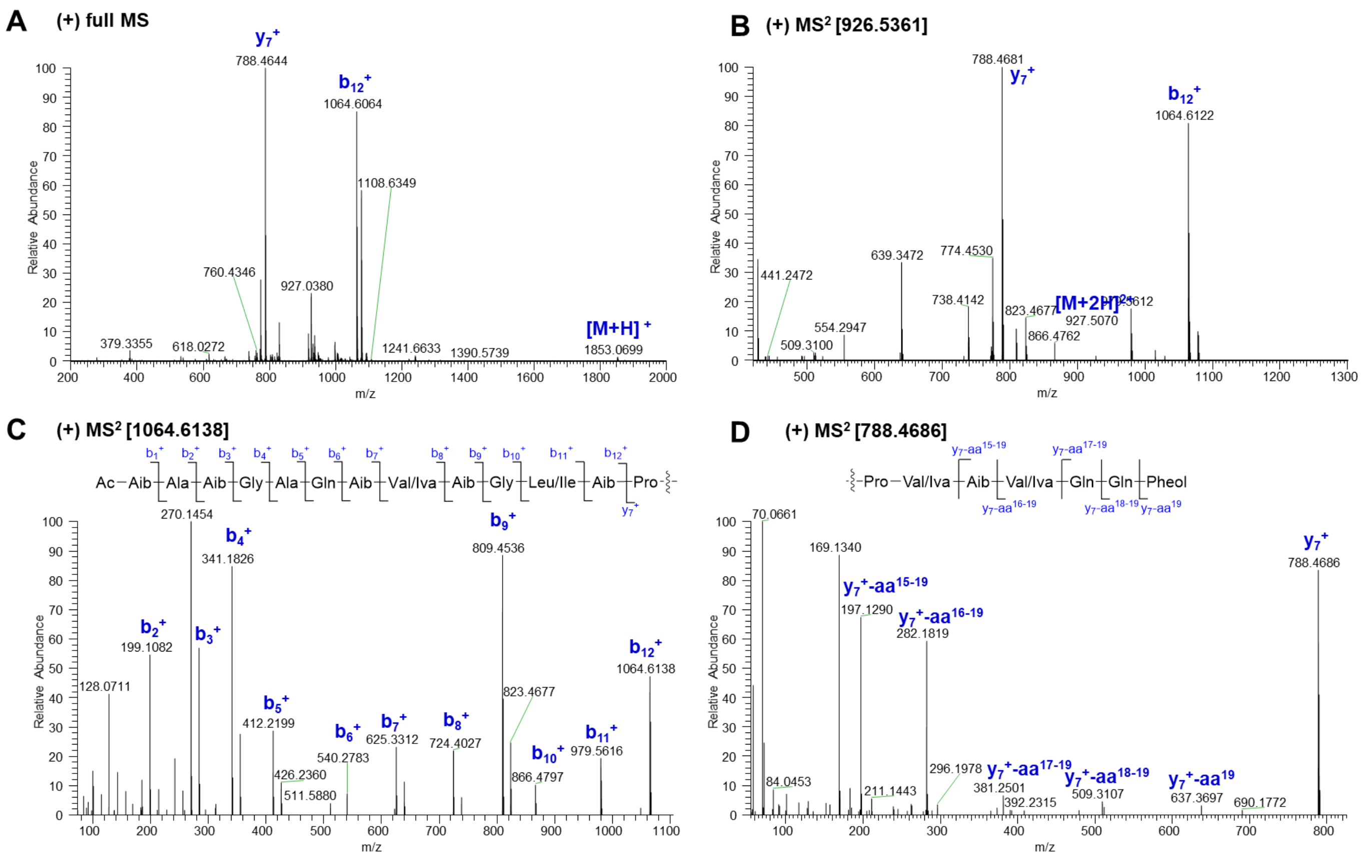

**Fig. S21** Diagnostic fragment ions [*m/z*] of **5** in subfraction 1 (SF1) by ESI-HRMS analysis

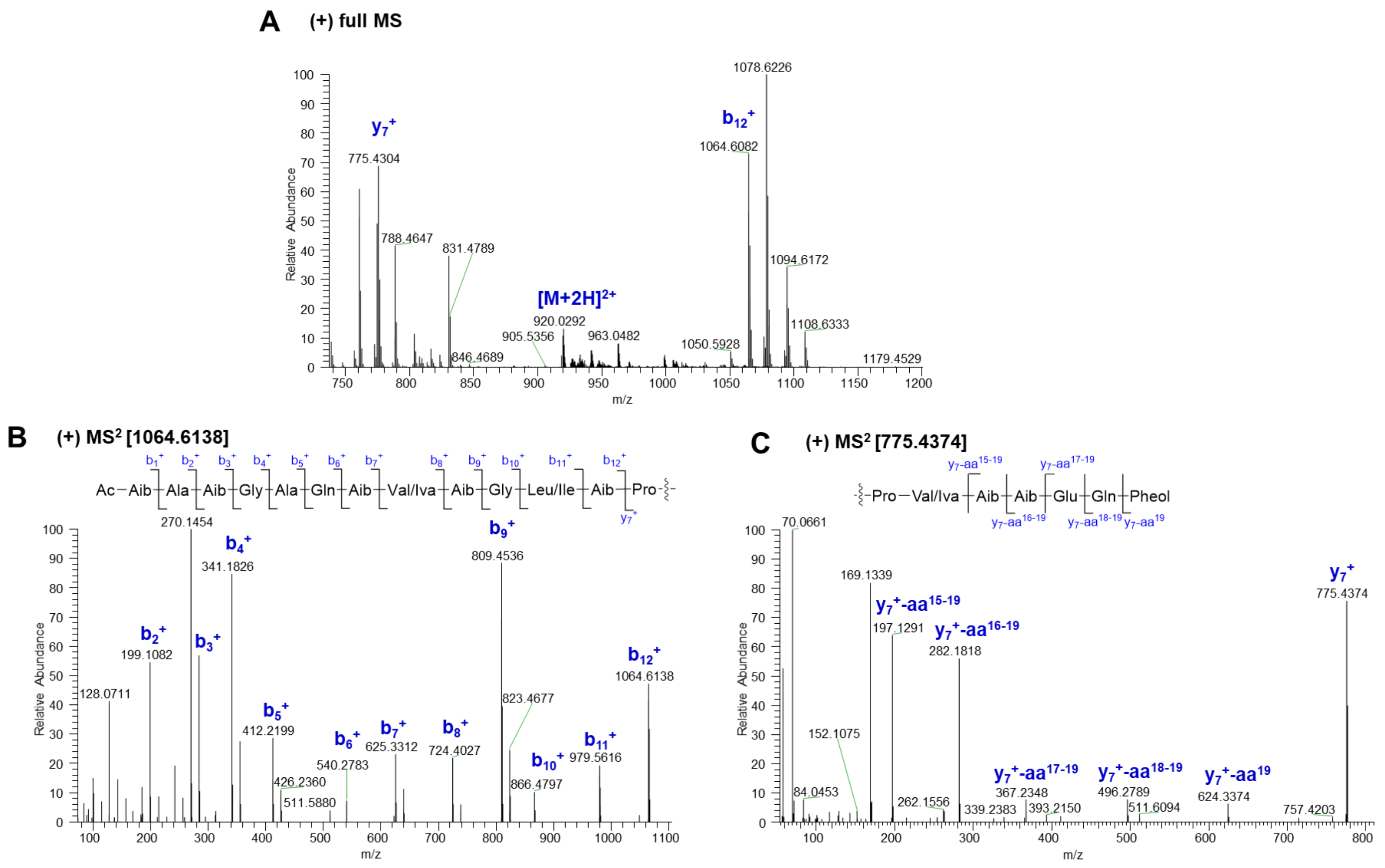

**Fig. S22** Diagnostic fragment ions [*m/z*] of **6** in subfraction 1 (SF1) by ESI-HRMS analysis

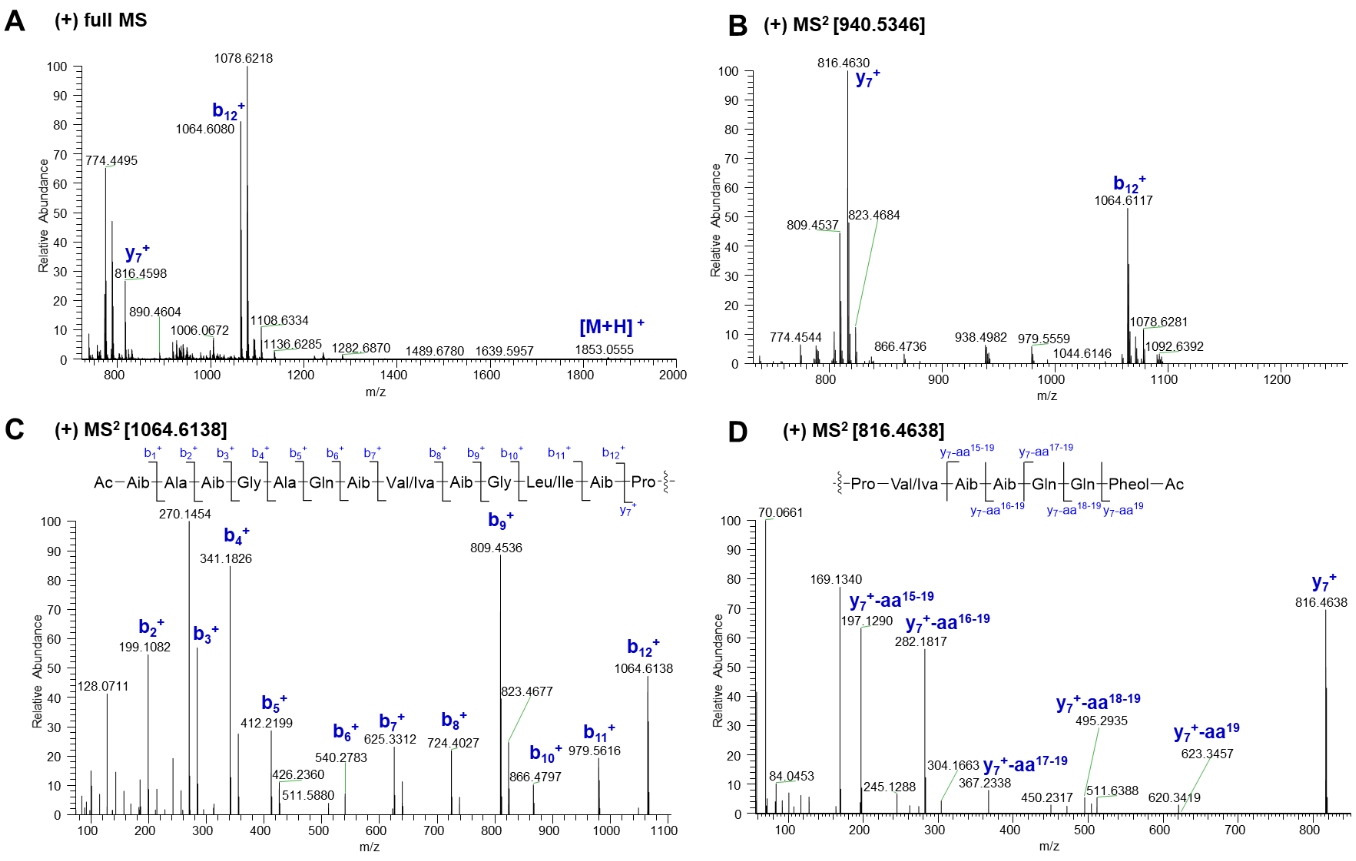

**Fig. S23** Diagnostic fragment ions [*m/z*] of **7** in subfraction 1 (SF1) by ESI-HRMS analysis

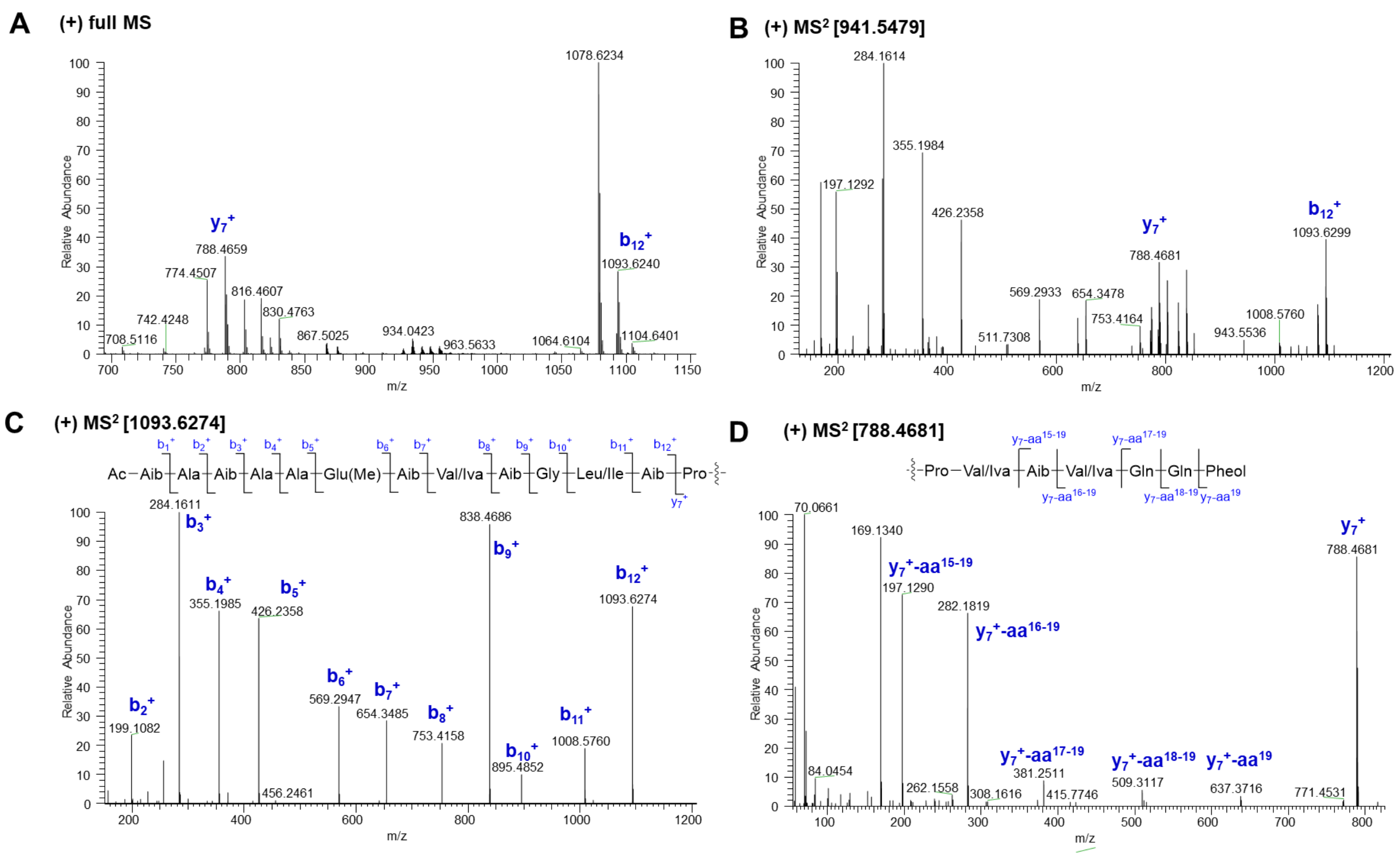

**Fig. S24** Diagnostic fragment ions [*m/z*] of **8** in subfraction 5 (SF5) by ESI-HRMS analysis

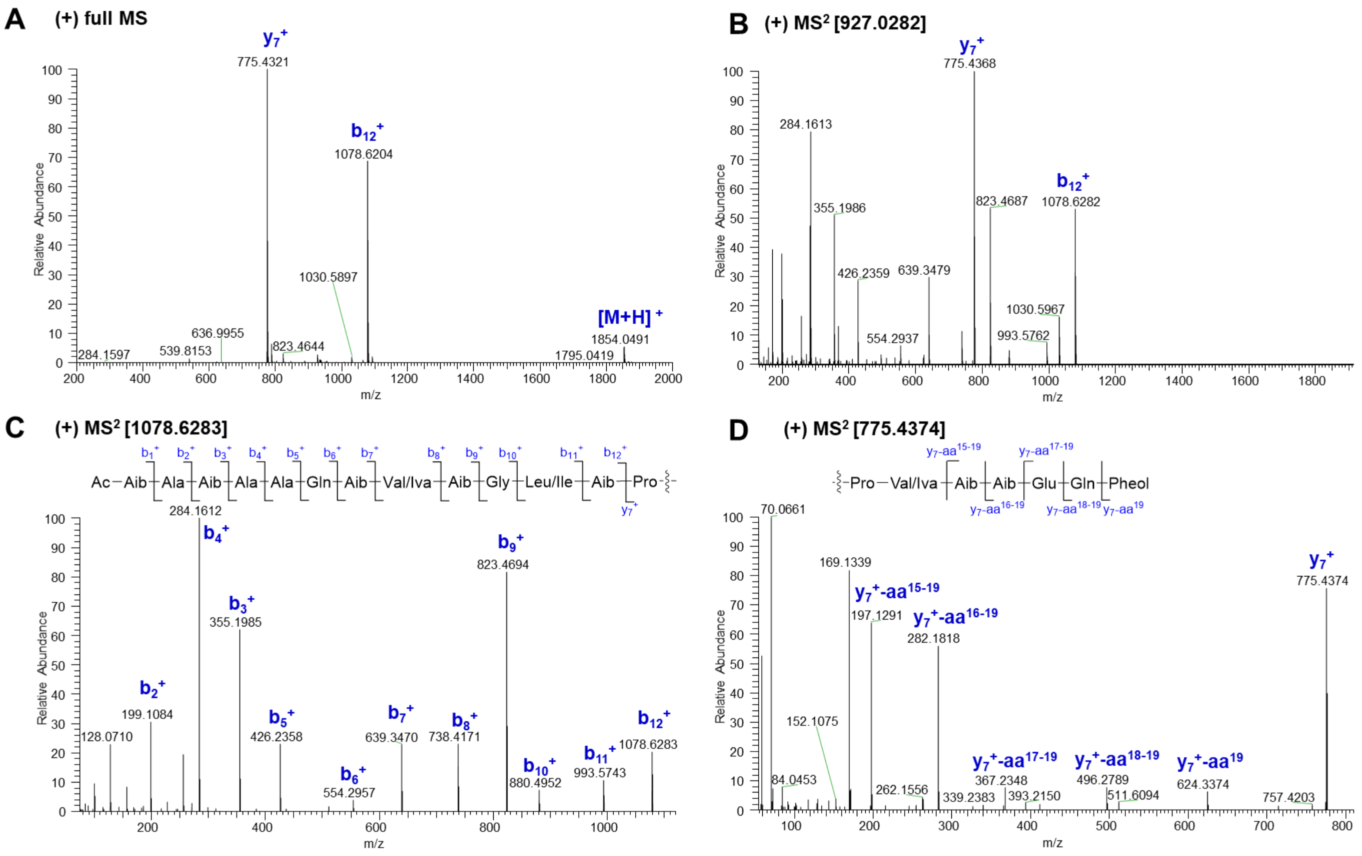

**Fig. S25** Diagnostic fragment ions [*m/z*] of **9** in subfraction 2 (SF2) by ESI-HRMS analysis

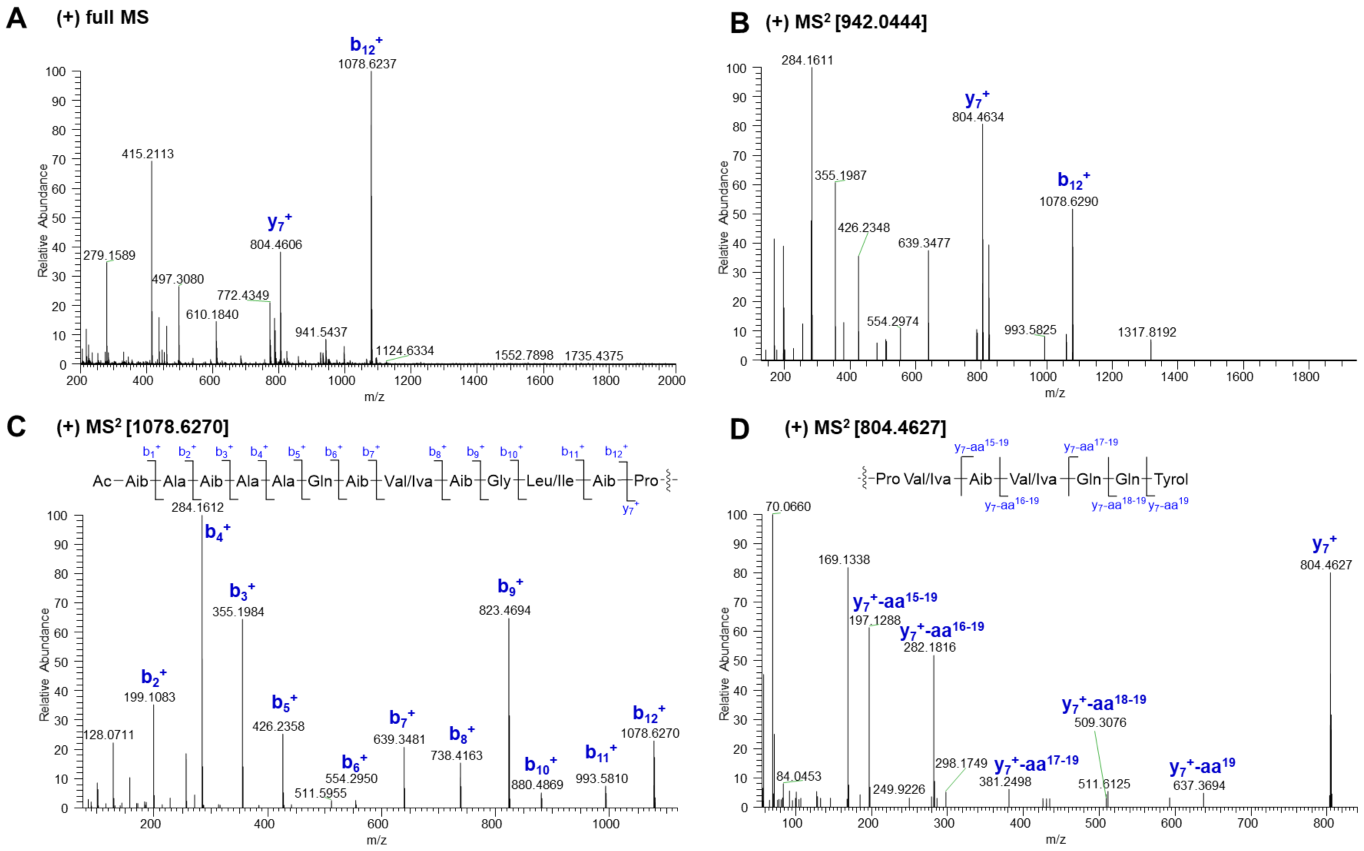

**Fig. S26** Diagnostic fragment ions [*m/z*] of **10** in subfraction 2 (SF2) by ESI-HRMS analysis

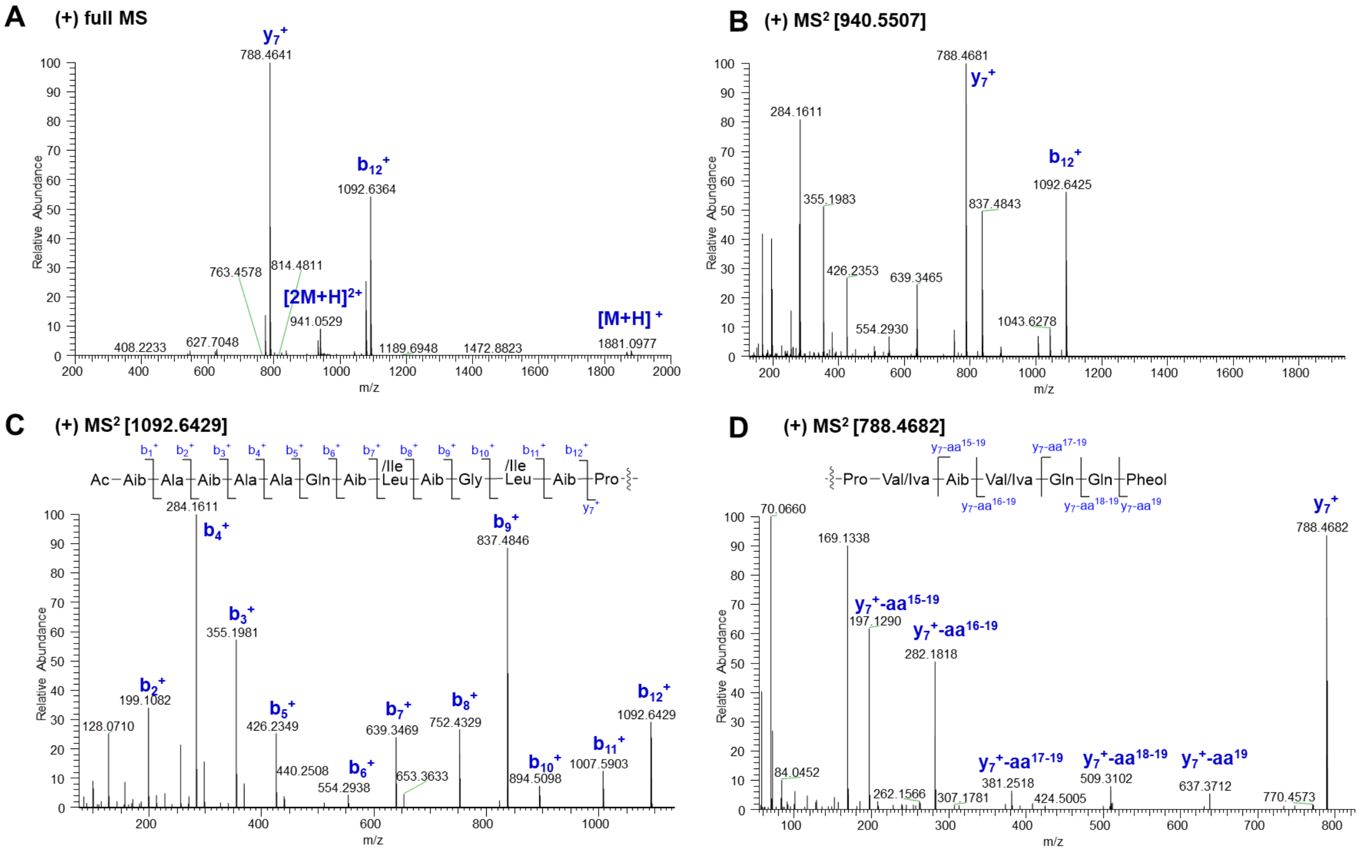

**Fig. S27** Diagnostic fragment ions [*m/z*] of **11** in subfraction 2 (SF2) by ESI-HRMS analysis

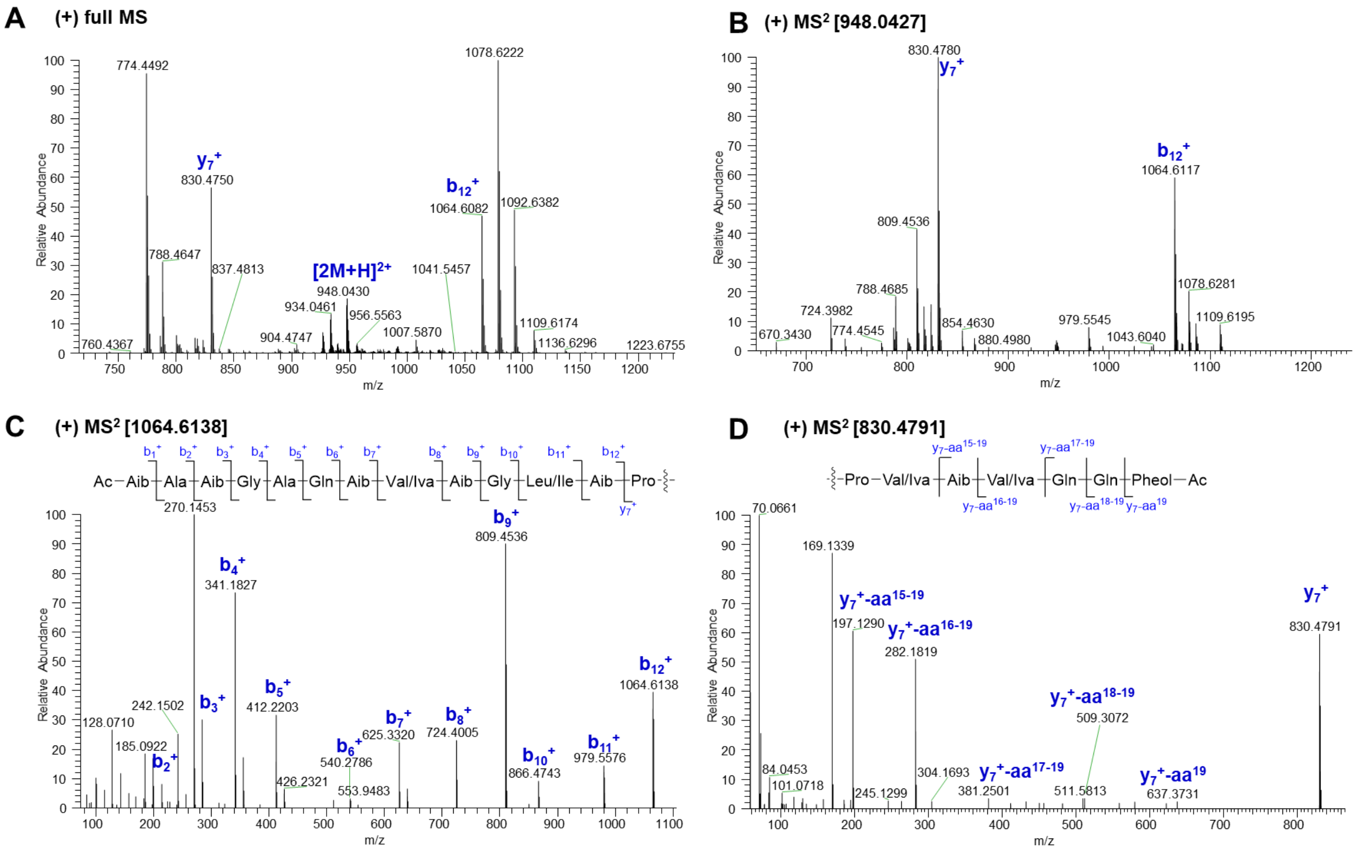

**Fig. S28** Diagnostic fragment ions [*m/z*] of **12** in subfraction 2 (SF2) by ESI-HRMS analysis

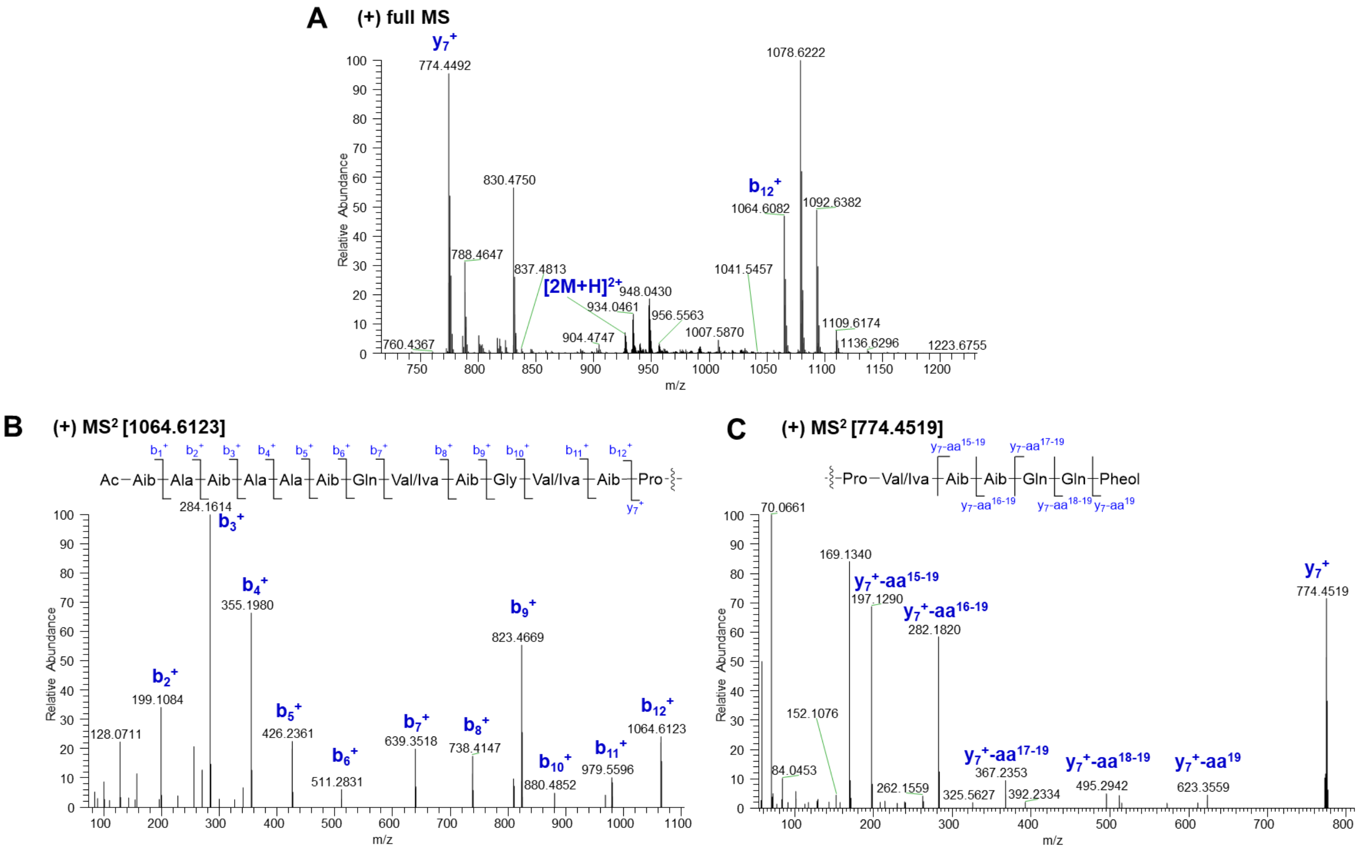

**Fig. S29** Diagnostic fragment ions [*m/z*] of **13** in subfraction 2 (SF2) by ESI-HRMS analysis

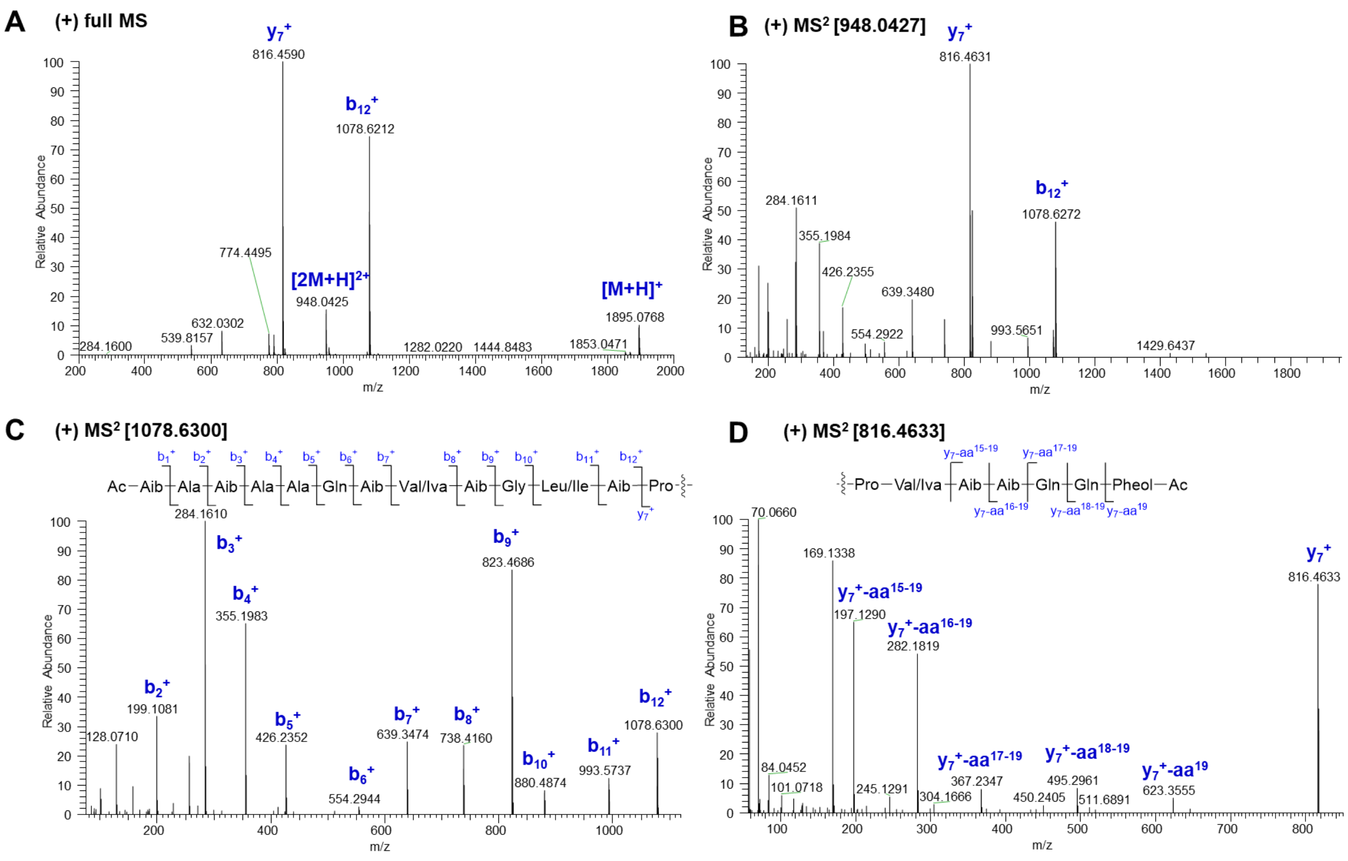

**Fig. S30** Diagnostic fragment ions [*m/z*] of **14** in subfraction 3 (SF3) by ESI-HRMS analysis

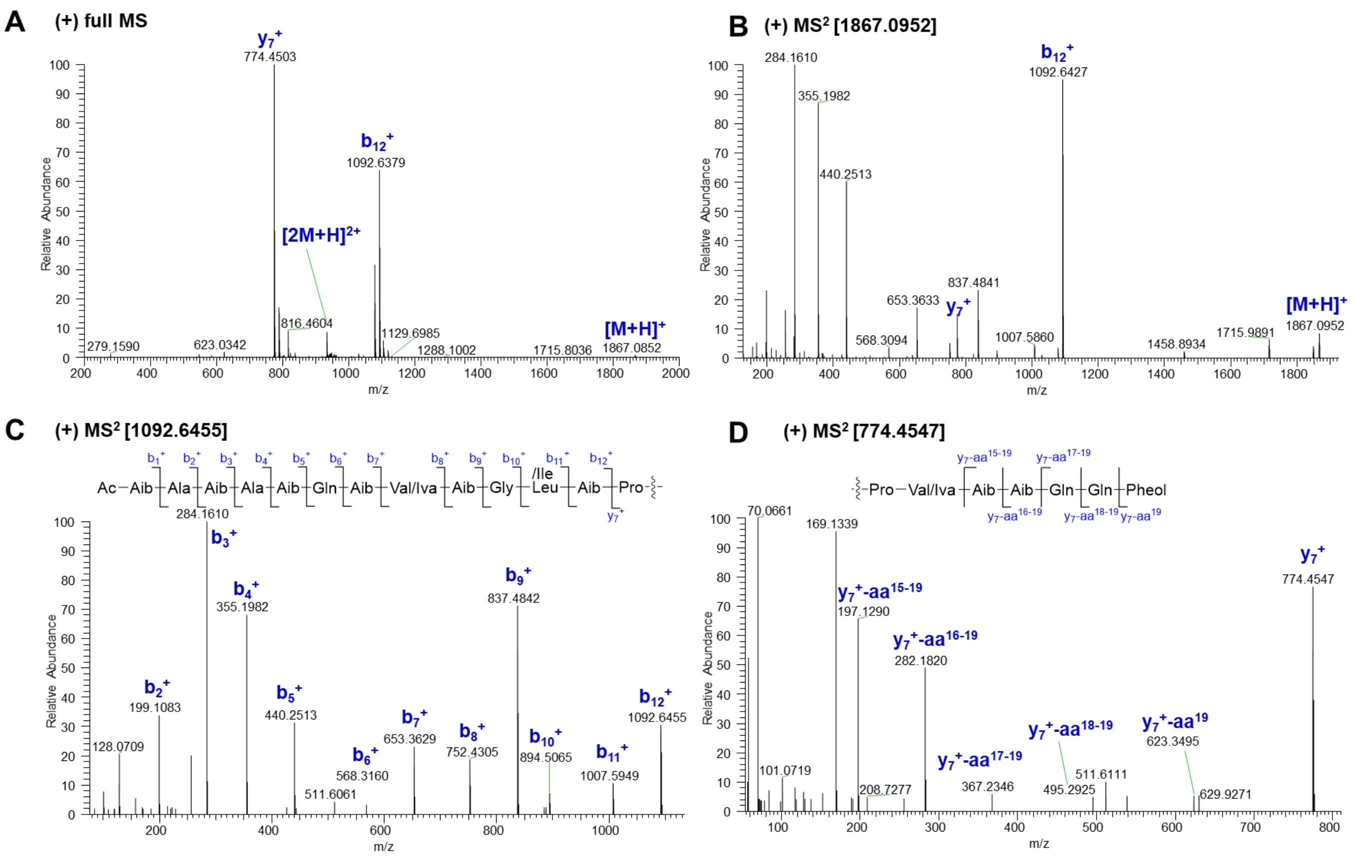

**Fig. S31** Diagnostic fragment ions [*m/z*] of **15** in subfraction 3 (SF3) by ESI-HRMS analysis

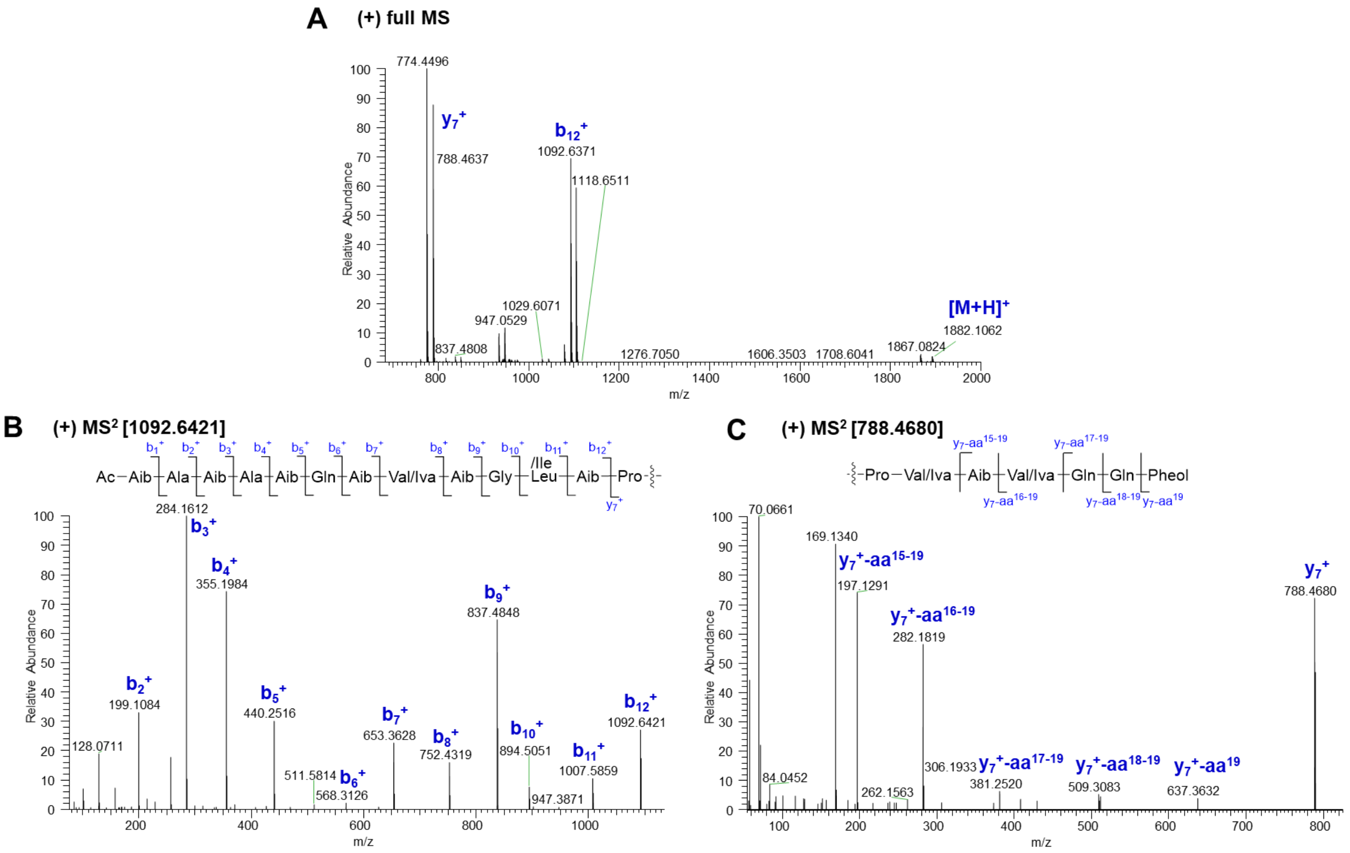

**Fig. S32** Diagnostic fragment ions [*m/z*] of **16** in subfraction 3 (SF3) by ESI-HRMS analysis

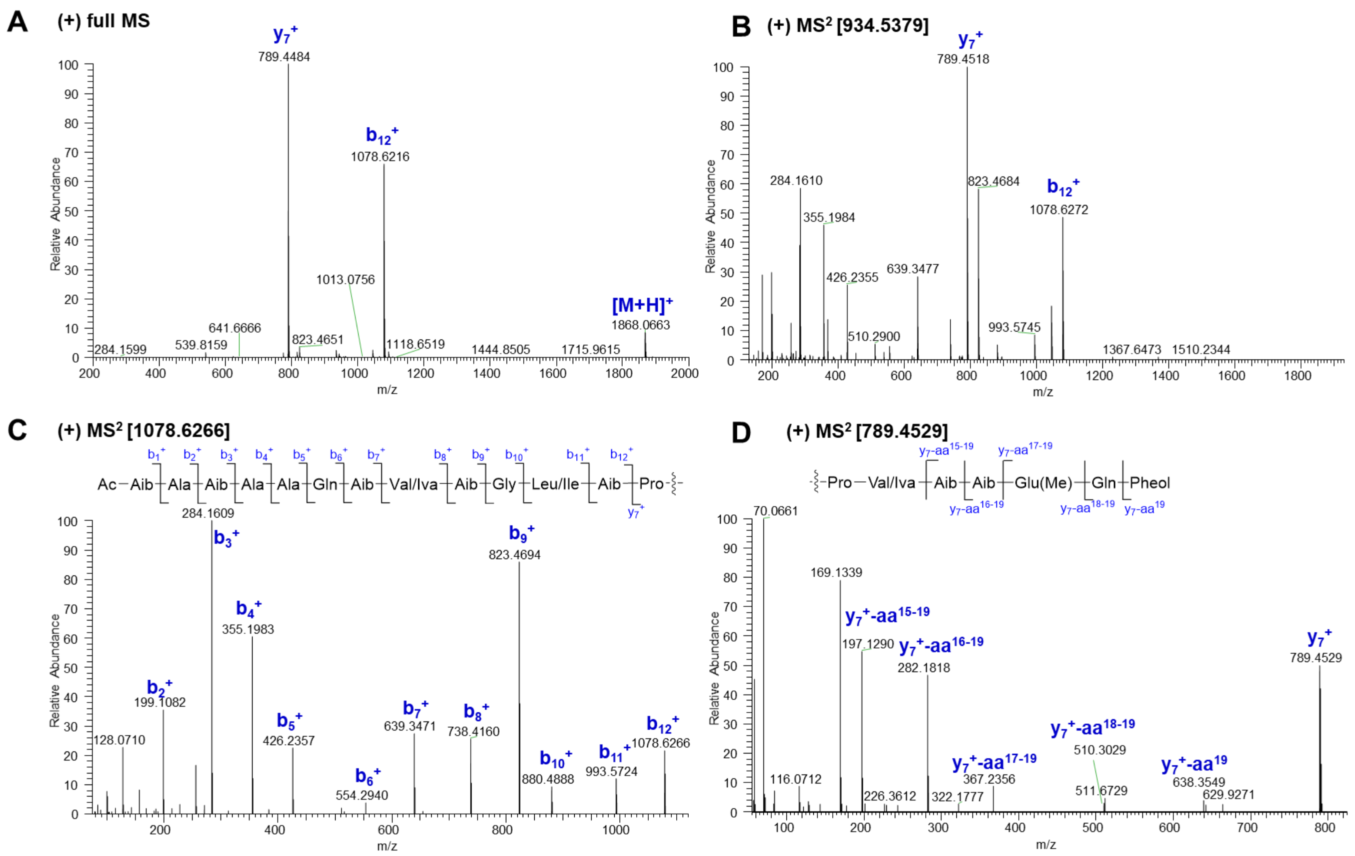

**Fig. S33** Diagnostic fragment ions [*m/z*] of **17** in subfraction 3 (SF3) by ESI-HRMS analysis

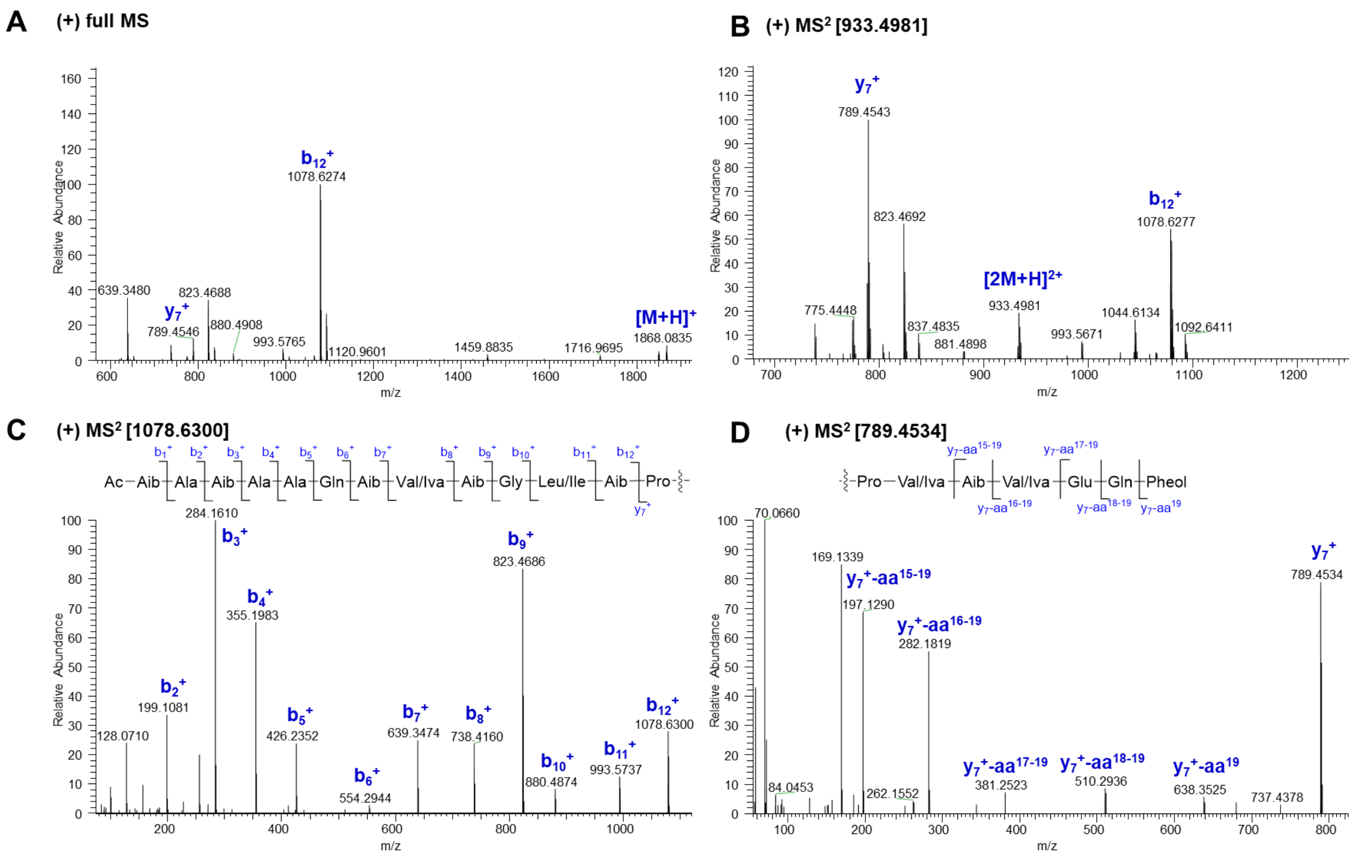

**Fig. S34** Diagnostic fragment ions [*m/z*] of **18** in subfraction 3 (SF3) by ESI-HRMS analysis

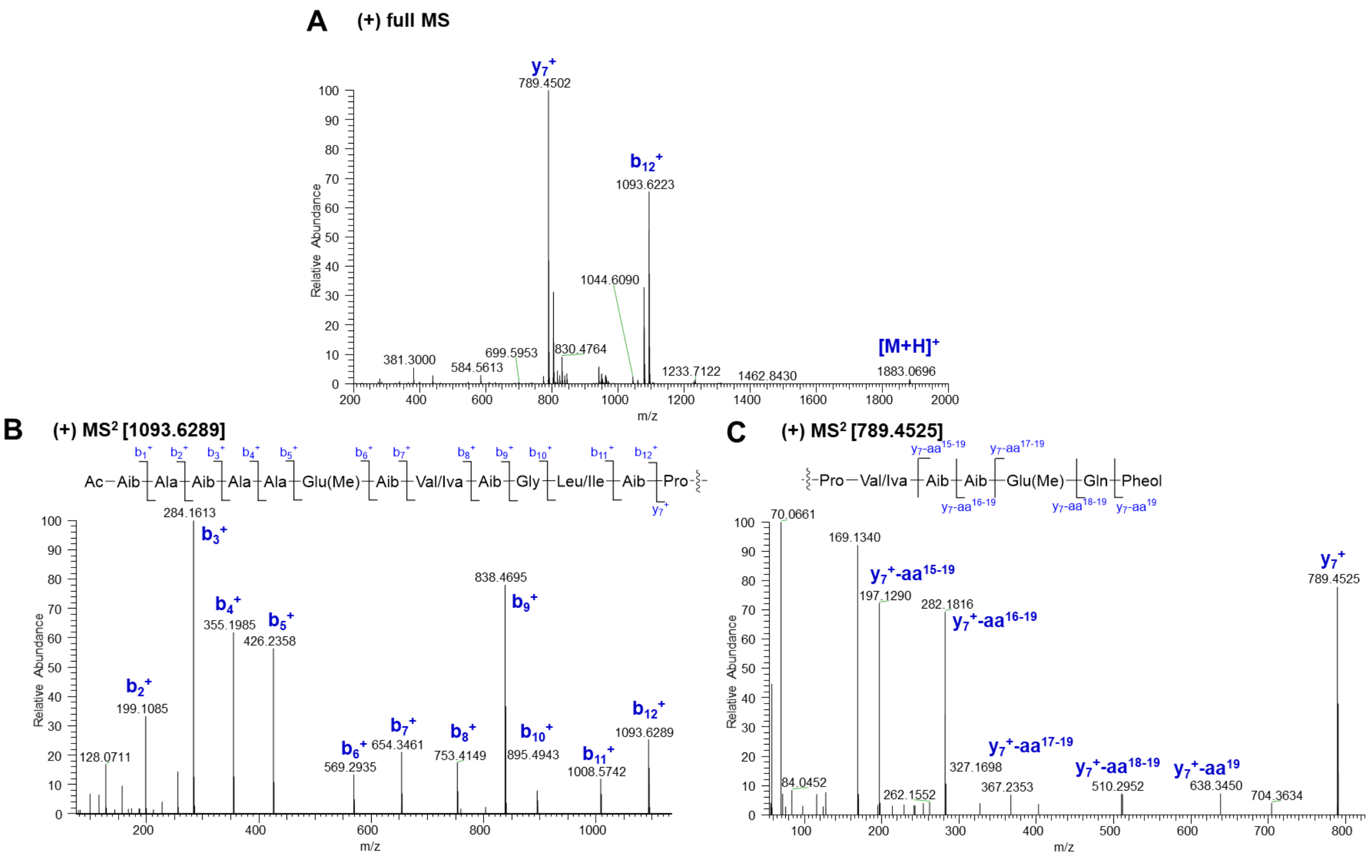

**Fig. S35** Diagnostic fragment ions [*m/z*] of **19** in subfraction 3 (SF3) by ESI-HRMS analysis

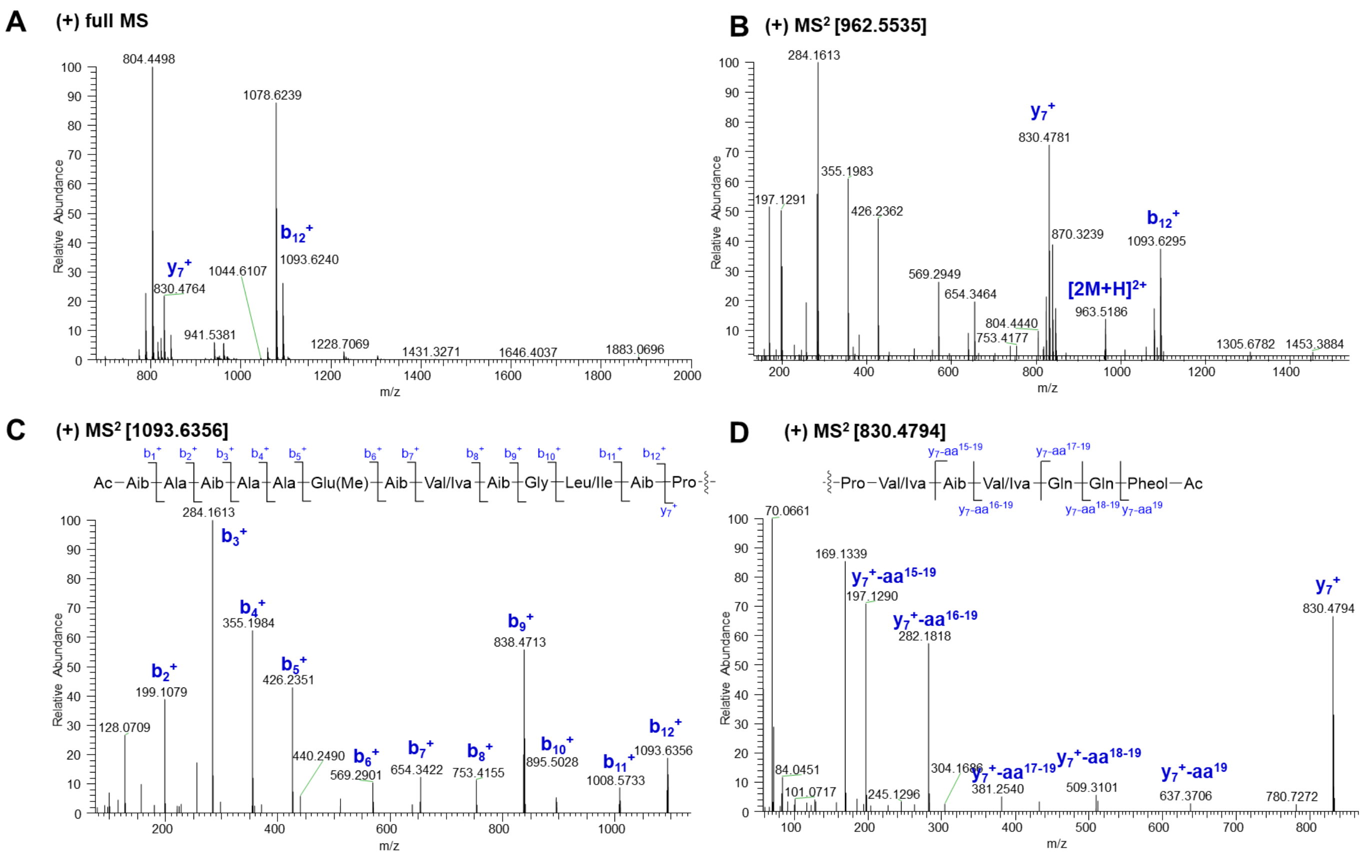

**Fig. S36** Diagnostic fragment ions [*m/z*] of **20** in subfraction 3 (SF3) by ESI-HRMS analysis

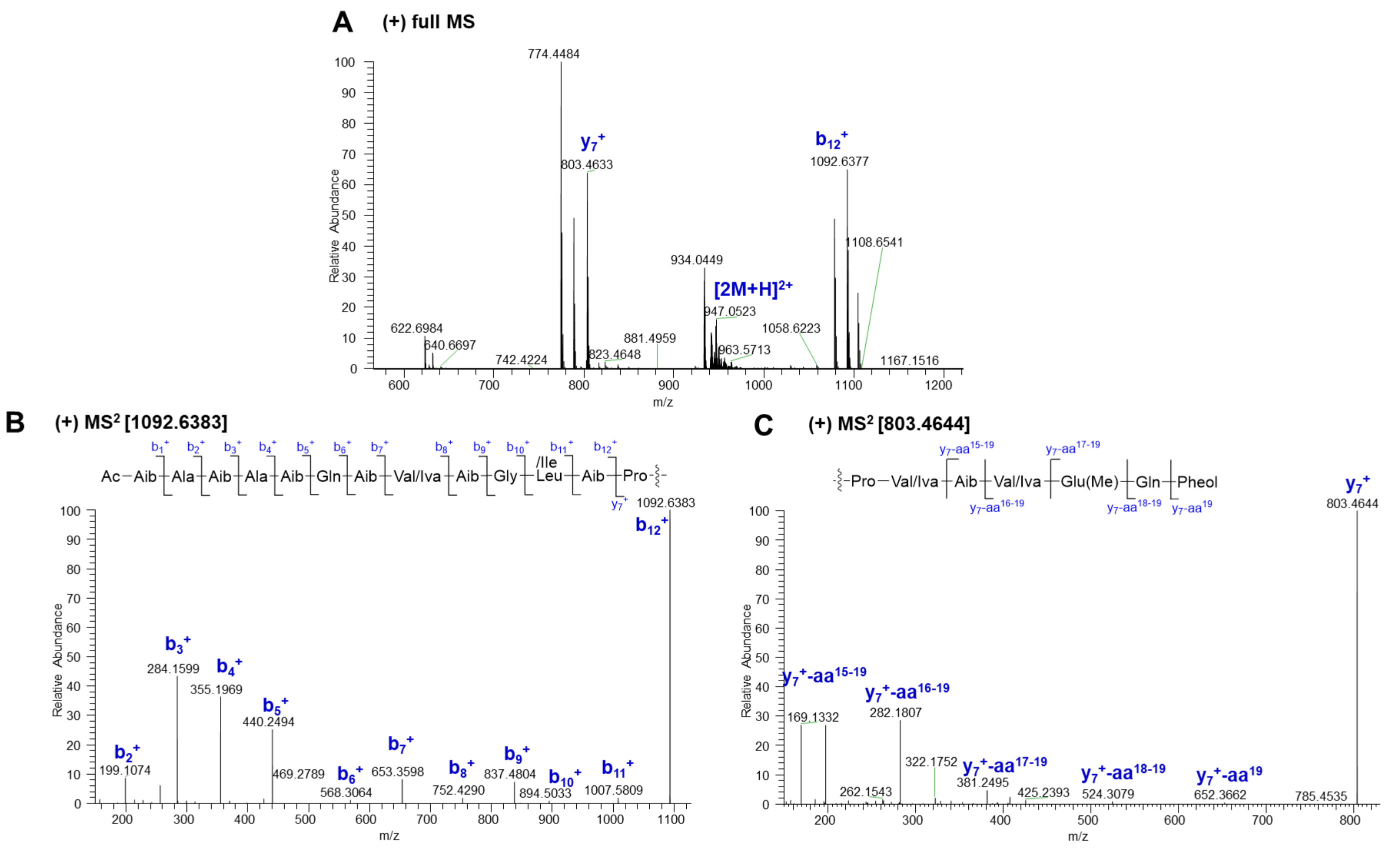

**Fig. S37** Diagnostic fragment ions [*m/z*] of **21** in subfraction 3 (SF3) by ESI-HRMS analysis

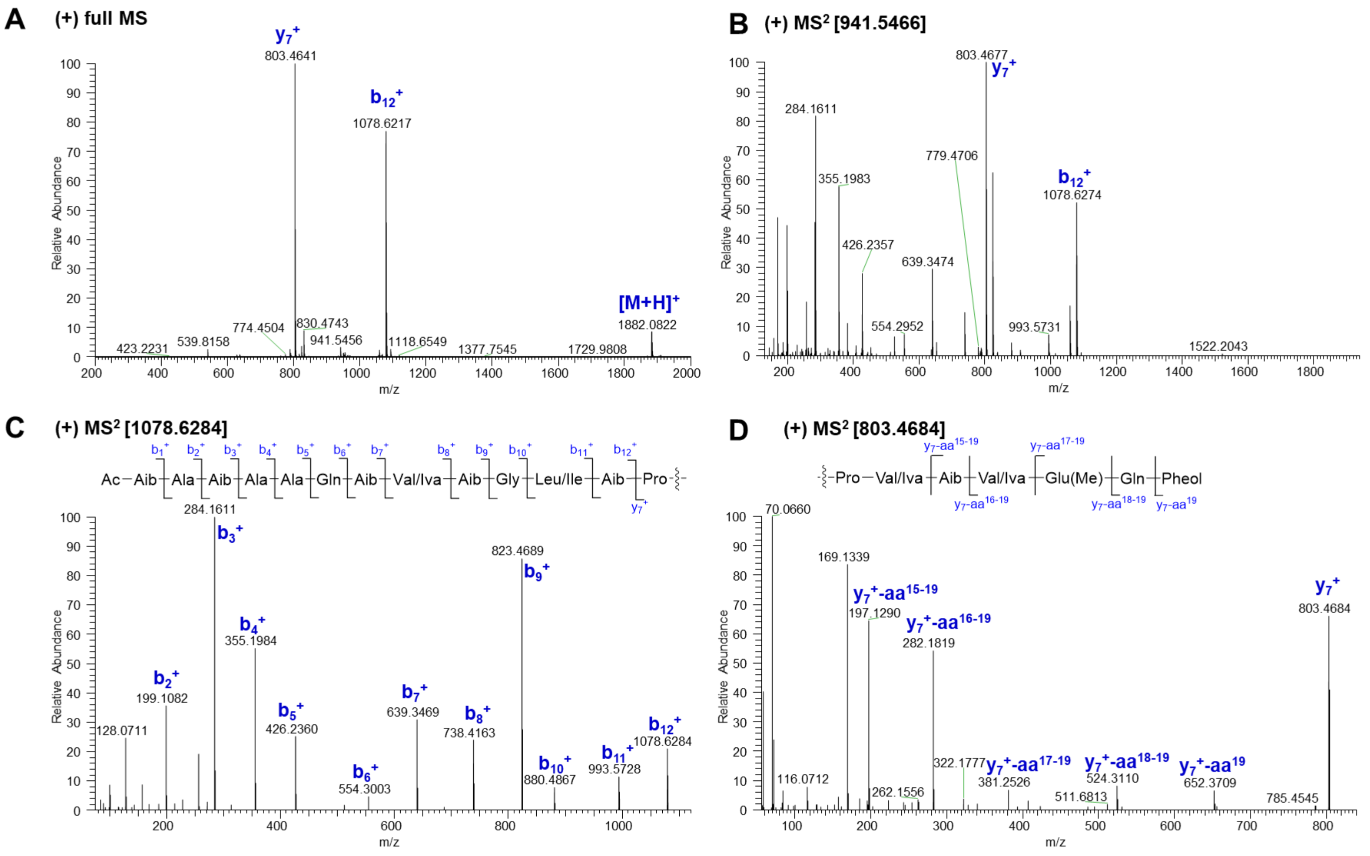

**Fig. S38** Diagnostic fragment ions [*m/z*] of **22** in subfraction 4 (SF4) by ESI-HRMS analysis

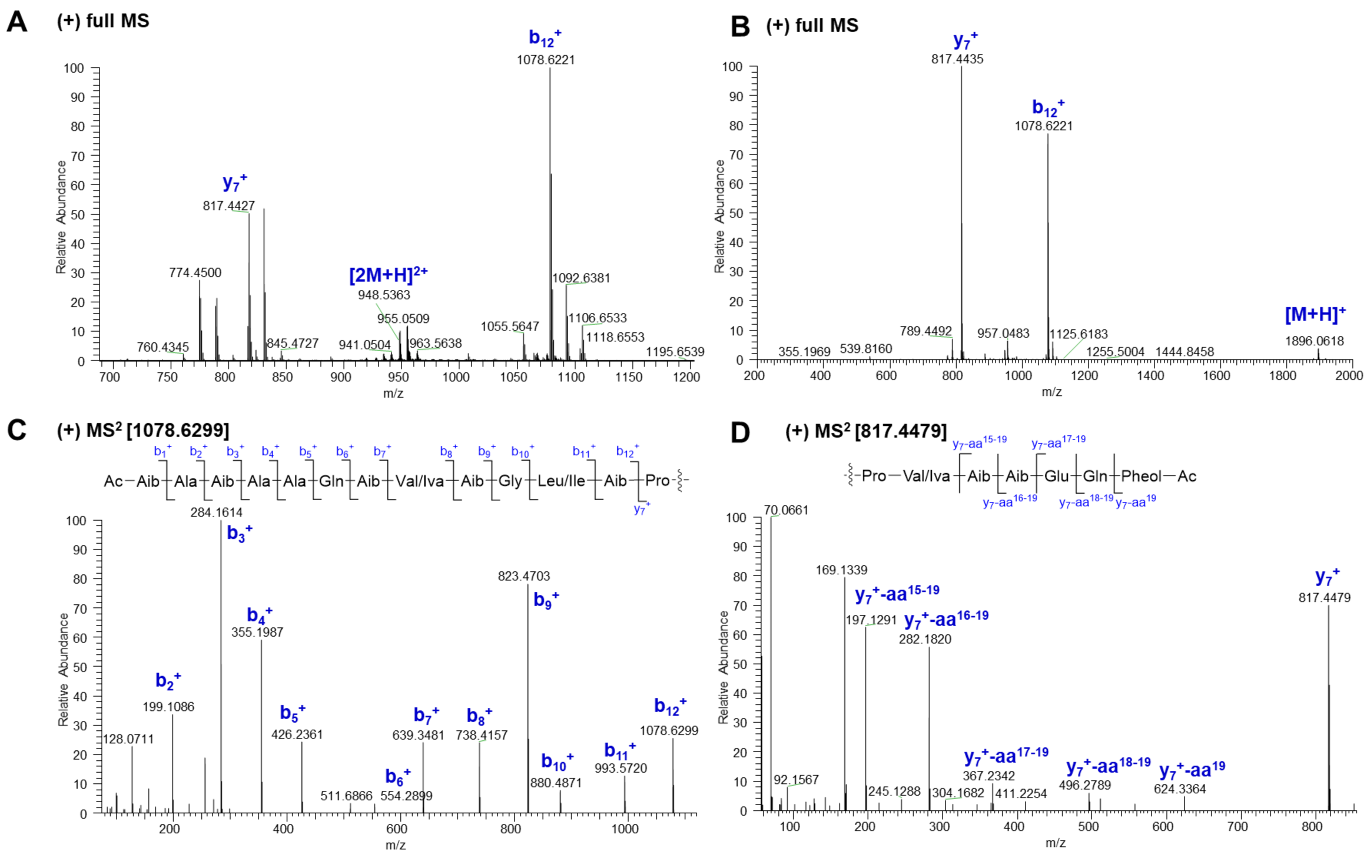

**Fig. S39** Diagnostic fragment ions [*m/z*] of **23** in subfraction 4 (SF4) by ESI-HRMS analysis

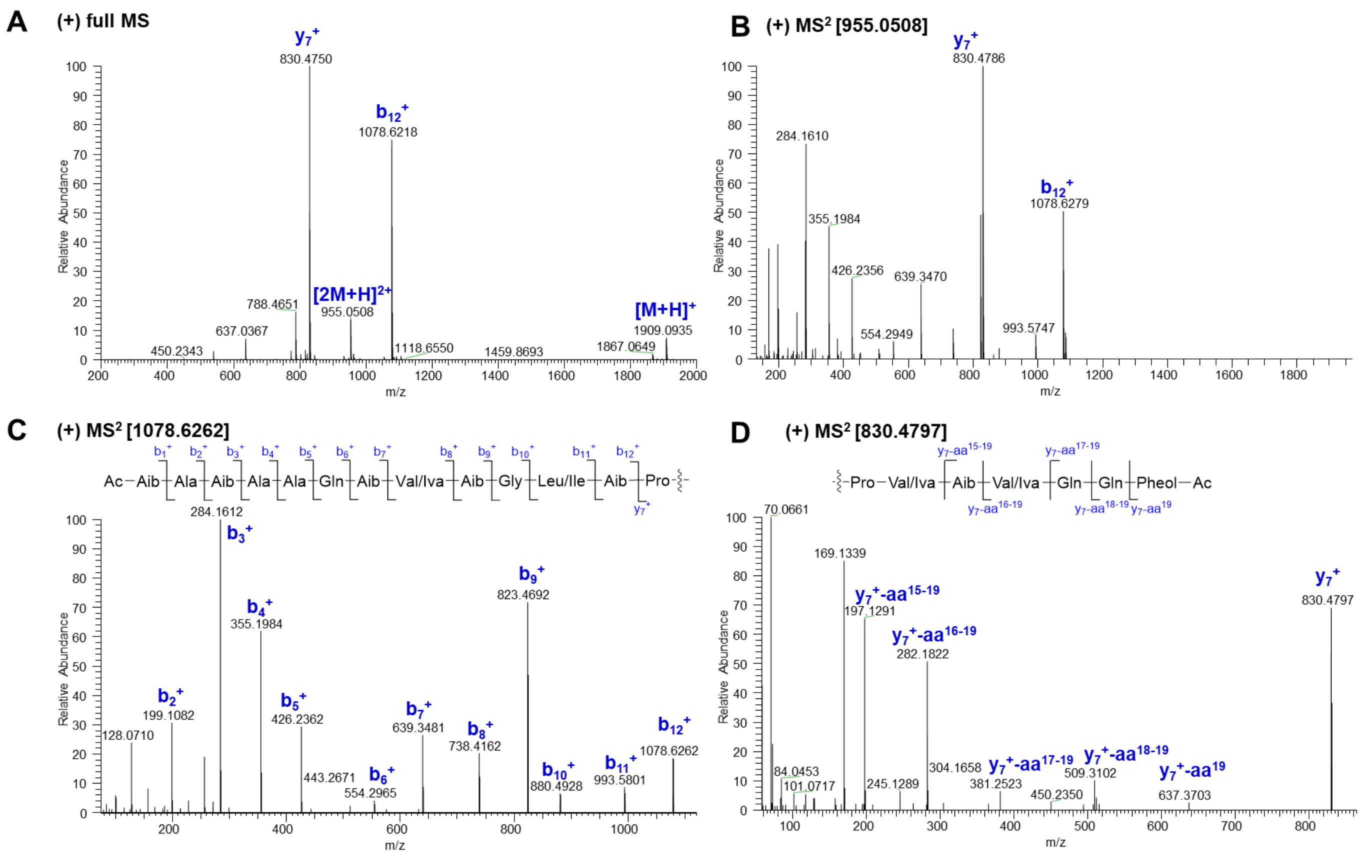

**Fig. S40** Diagnostic fragment ions [*m/z*] of **24** in subfraction 4 (SF4) by ESI-HRMS analysis

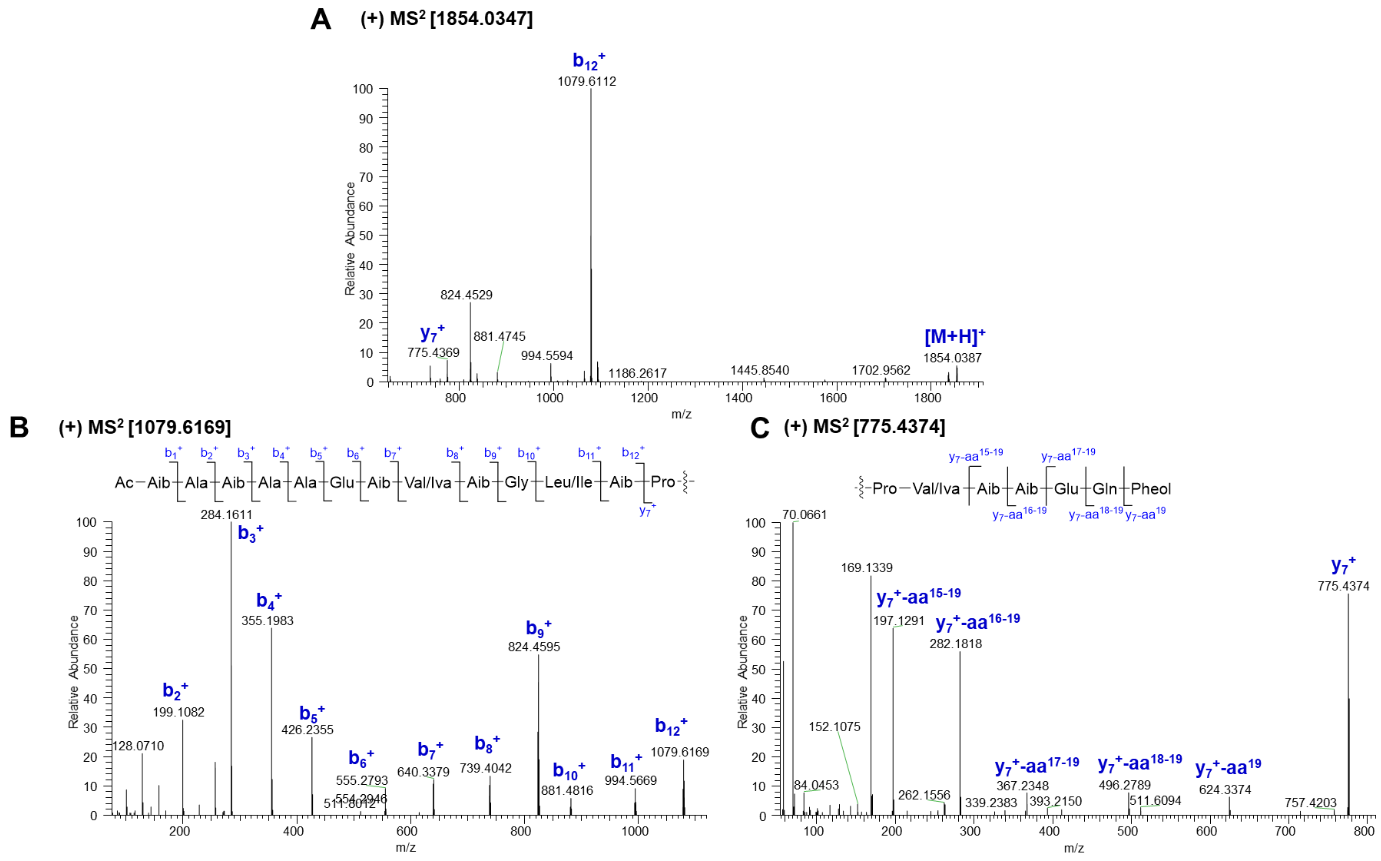

**Fig. S41** Diagnostic fragment ions [*m/z*] of **25** in subfraction 4 (SF4) by ESI-HRMS analysis

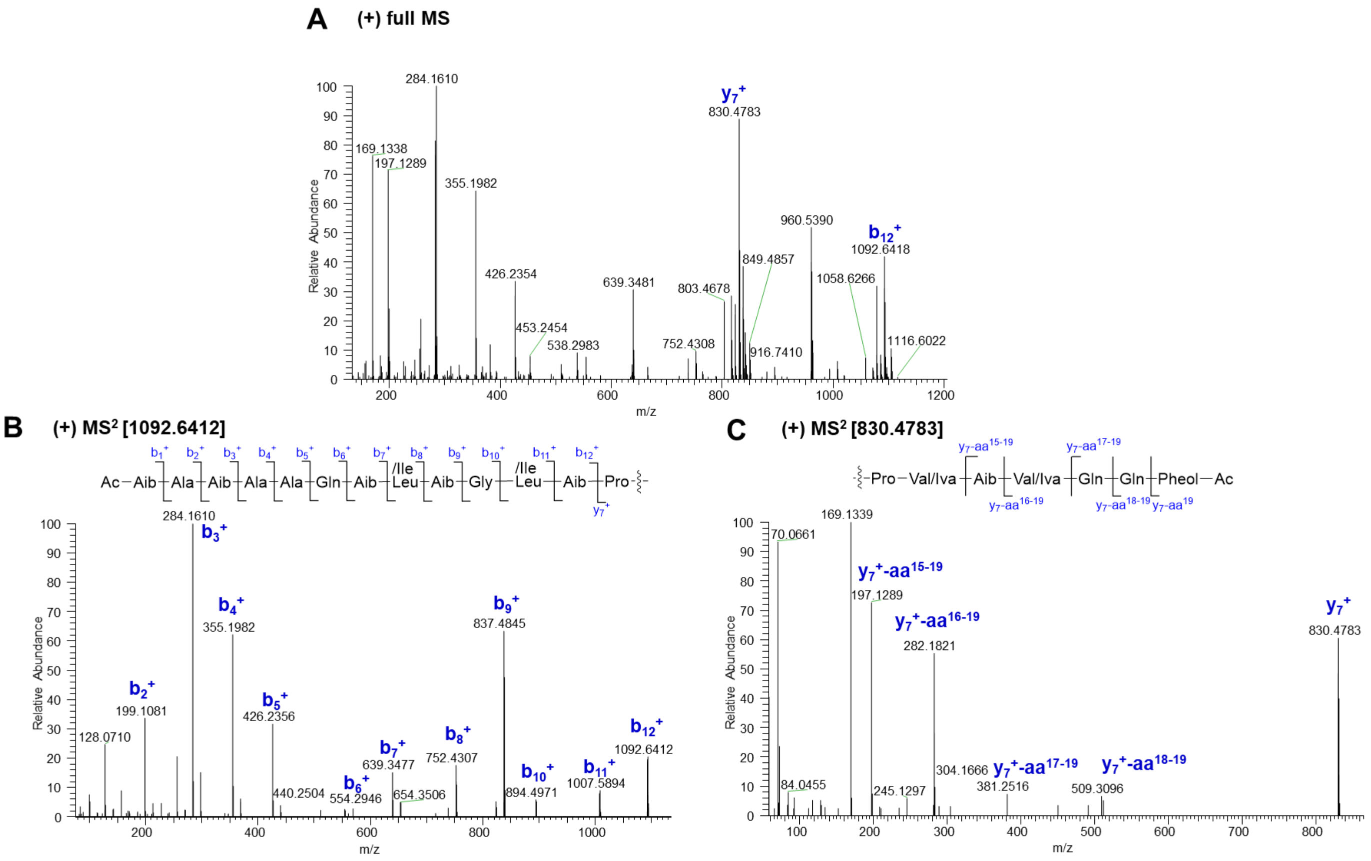

**Fig. S42** Diagnostic fragment ions [*m/z*] of **26** in subfraction 4 (SF4) by ESI-HRMS analysis

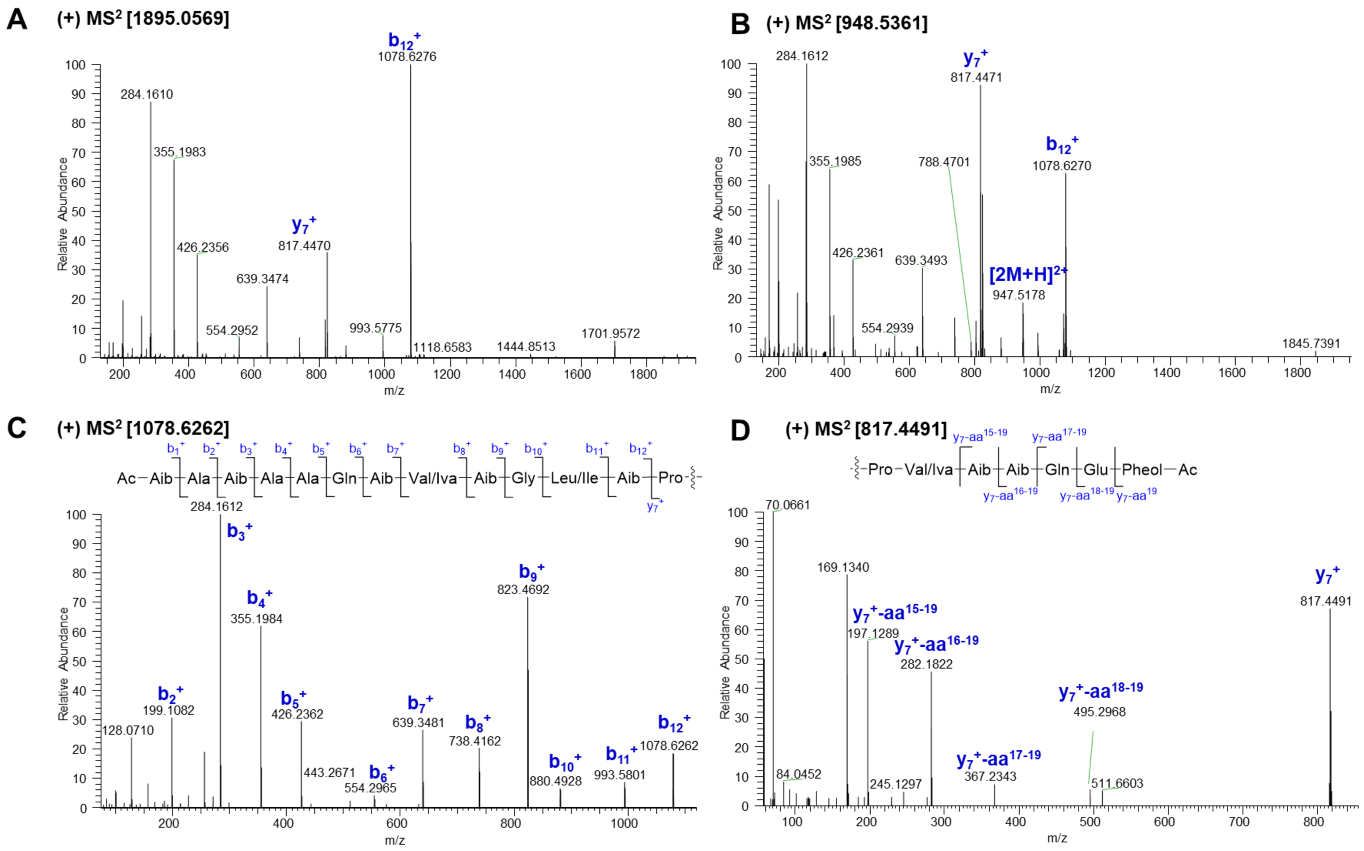

**Fig. S43** Diagnostic fragment ions [*m/z*] of **27** in subfraction 4 (SF4) by ESI-HRMS analysis

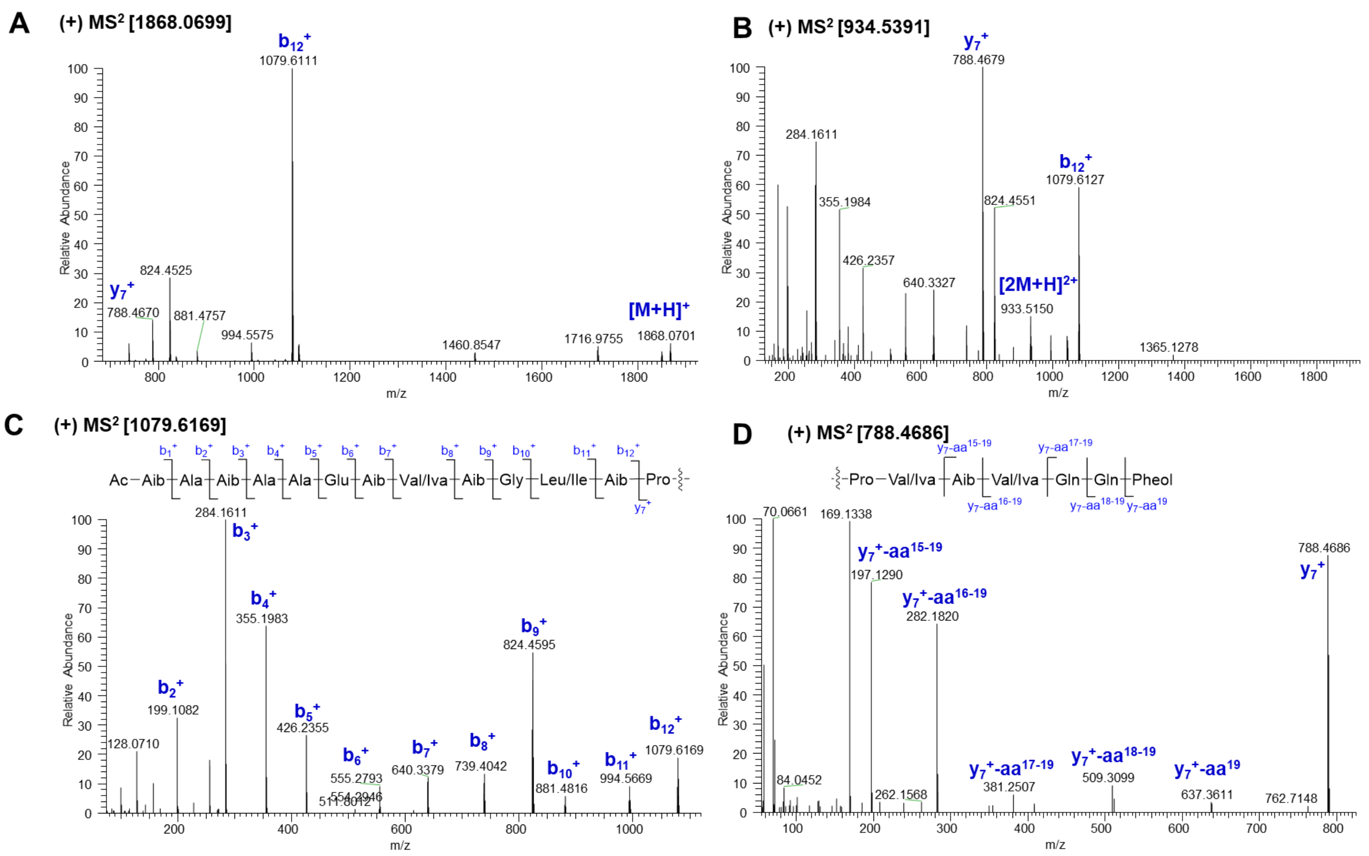

**Fig. S44** Diagnostic fragment ions [*m/z*] of **28** in subfraction 4 (SF4) by ESI-HRMS analysis

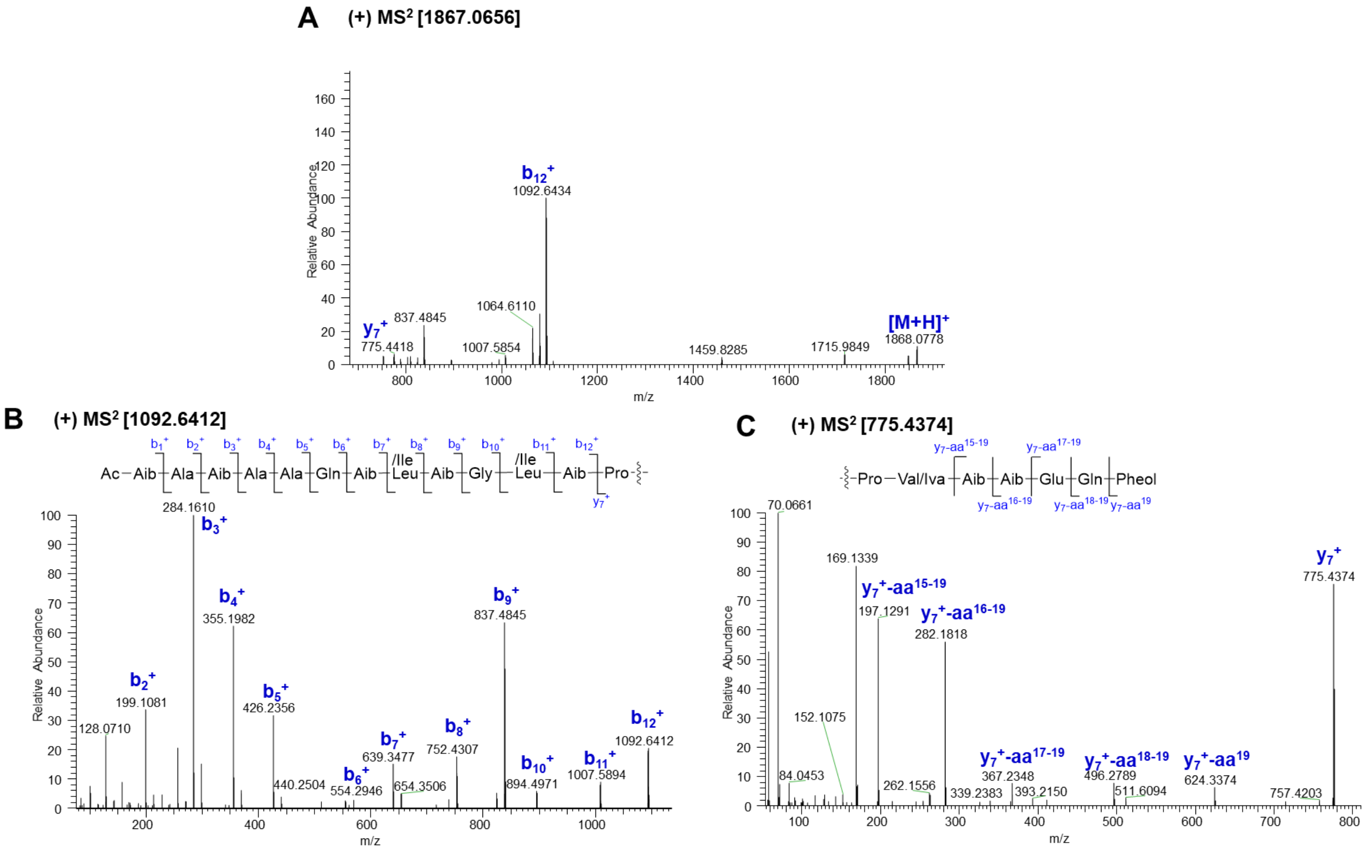

**Fig. S45** Diagnostic fragment ions [*m/z*] of **29** in subfraction 4 (SF4) by ESI-HRMS analysis

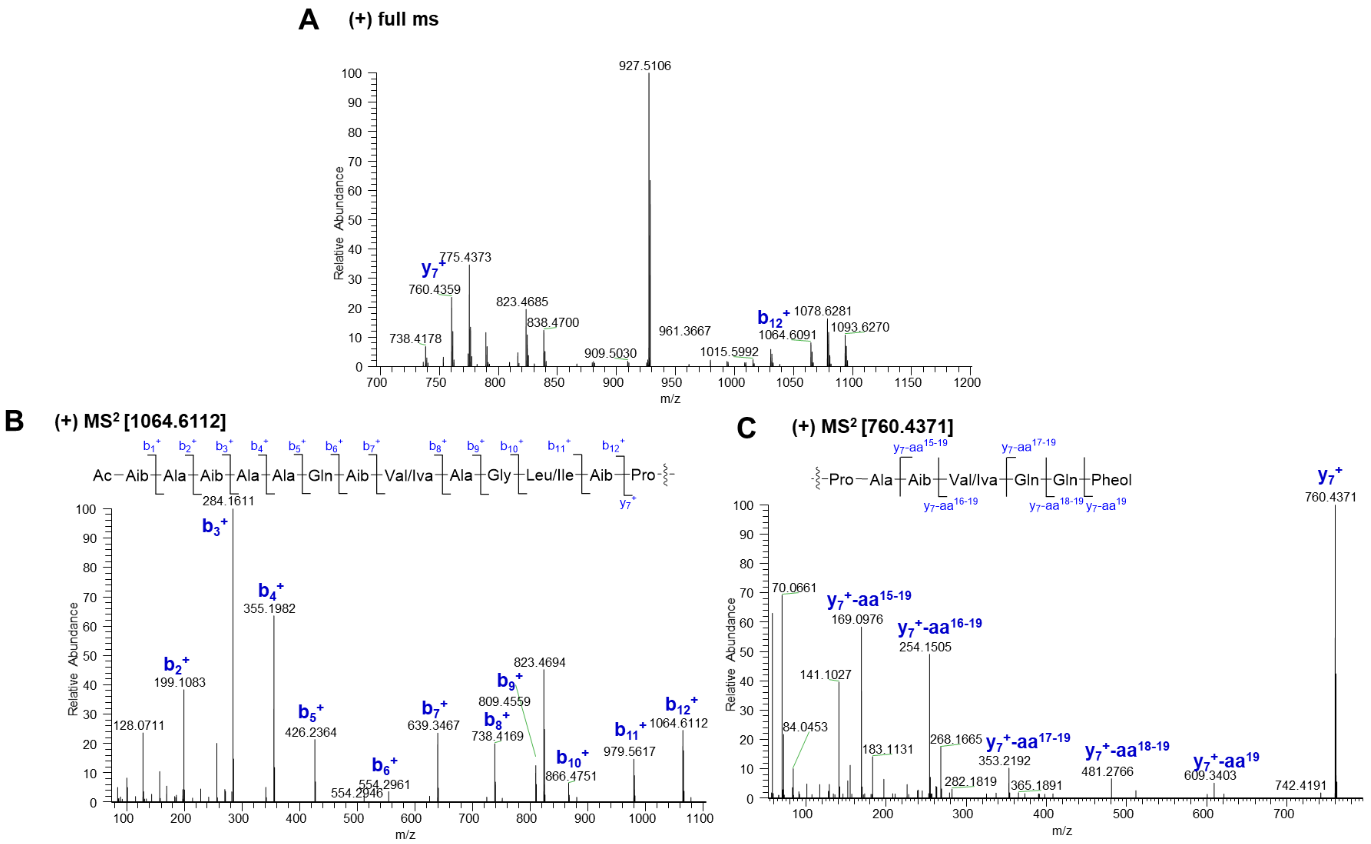

**Fig. S46** Diagnostic fragment ions [*m/z*] of **30** in subfraction 4 (SF4) by ESI-HRMS analysis

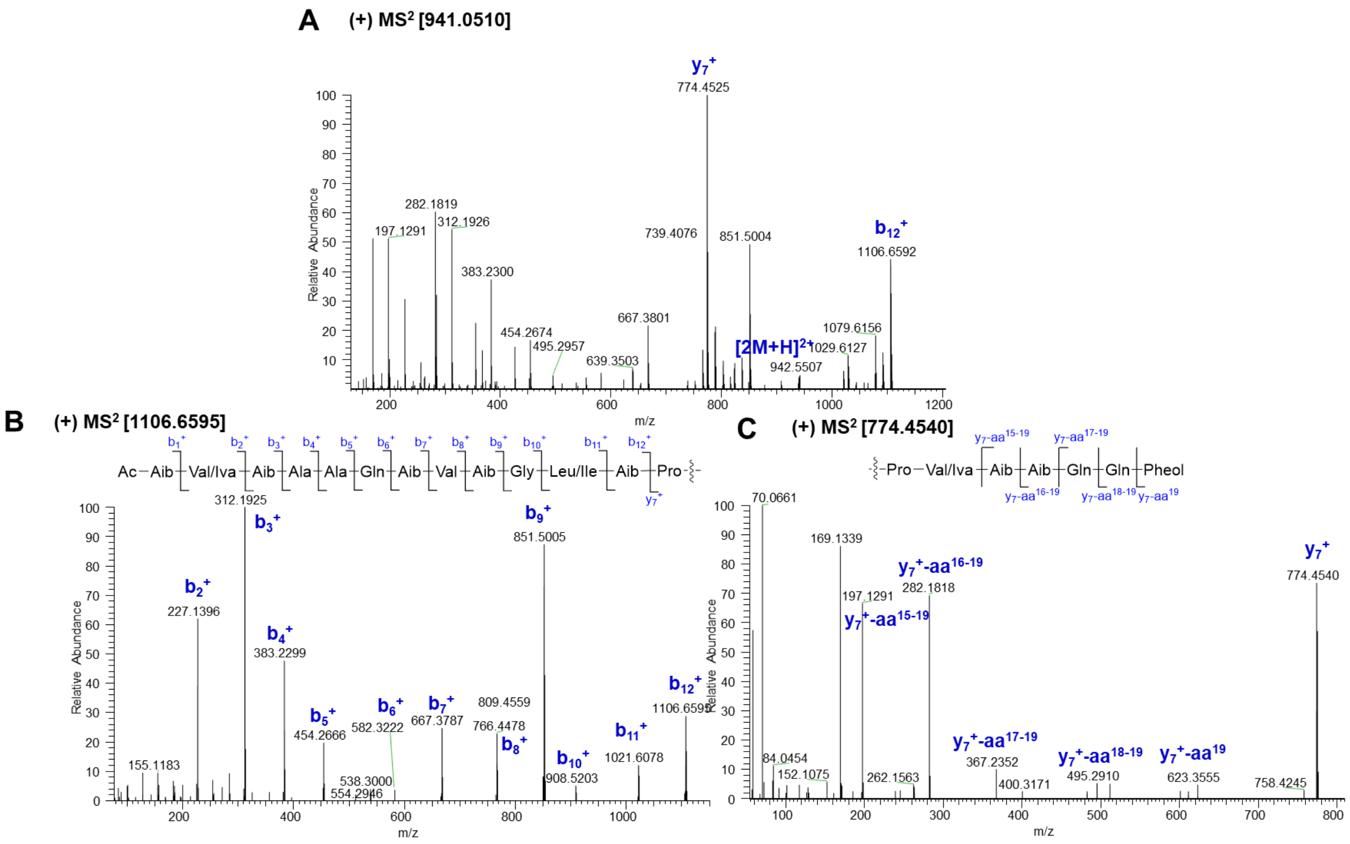

**Fig. S47** Diagnostic fragment ions [*m/z*] of **31** in subfraction 4 (SF4) by ESI-HRMS analysis

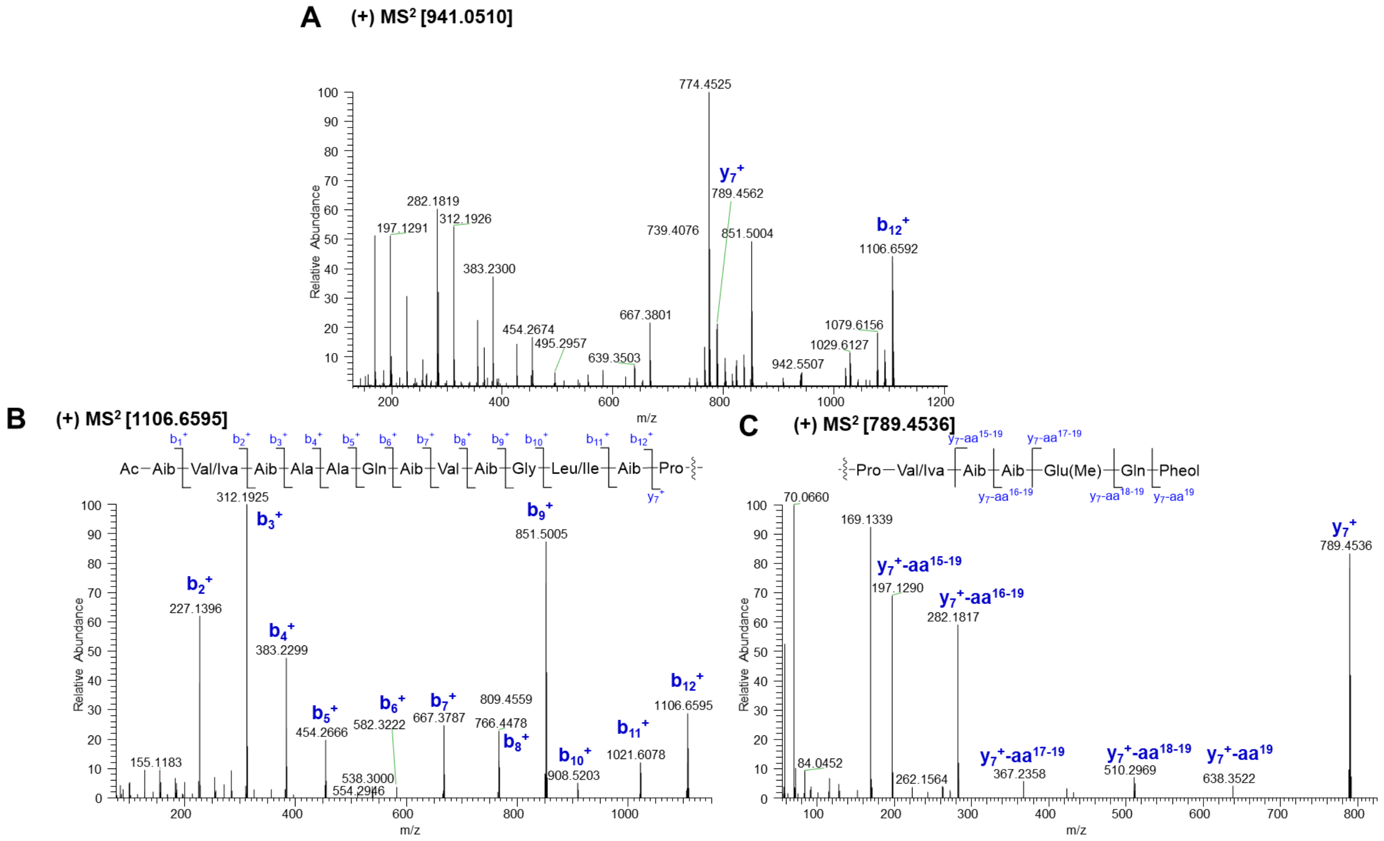

**Fig. S48** Diagnostic fragment ions [*m/z*] of **32** in subfraction 4 (SF4) by ESI-HRMS analysis

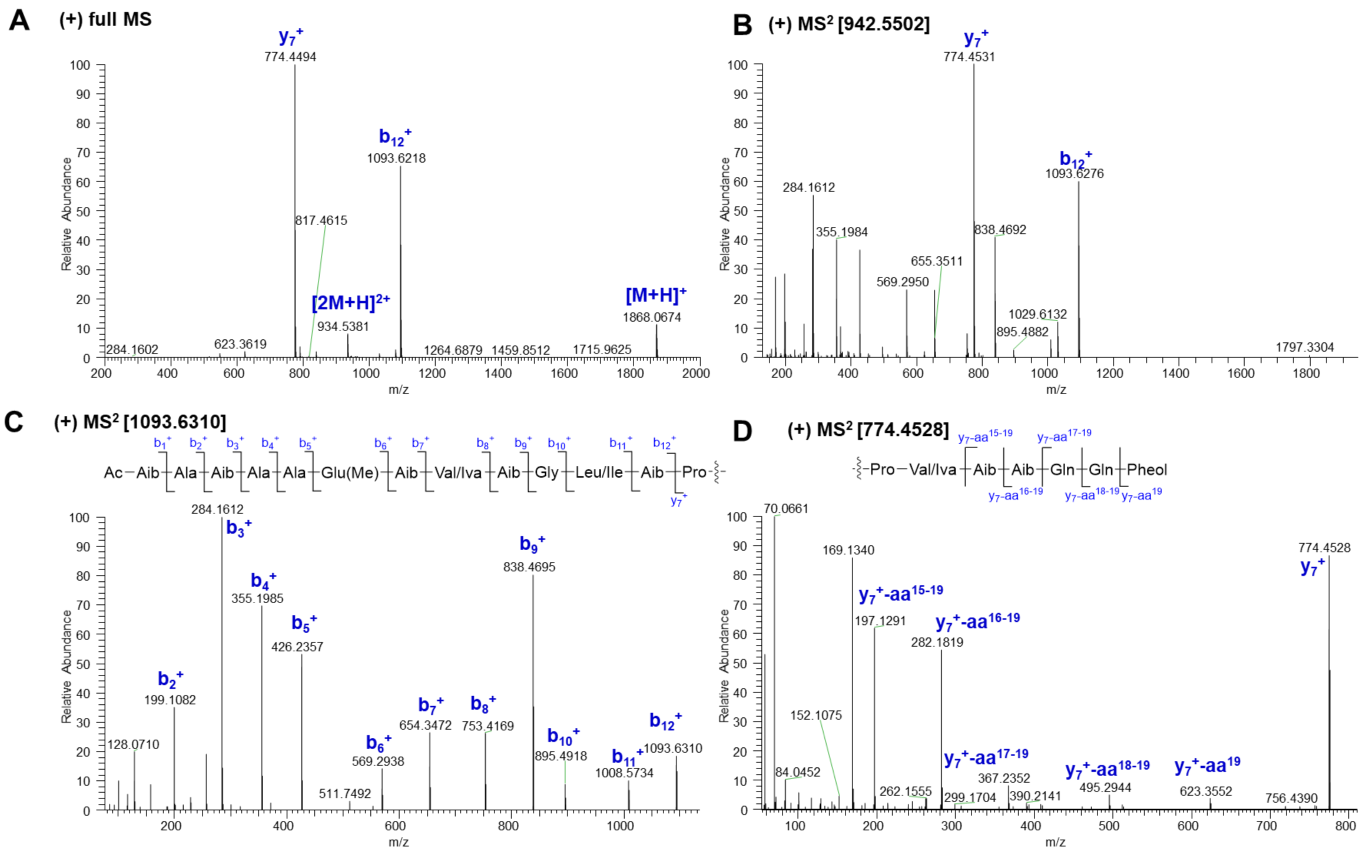

**Fig. S49** Diagnostic fragment ions [*m/z*] of **33** in subfraction 5 (SF5) by ESI-HRMS analysis

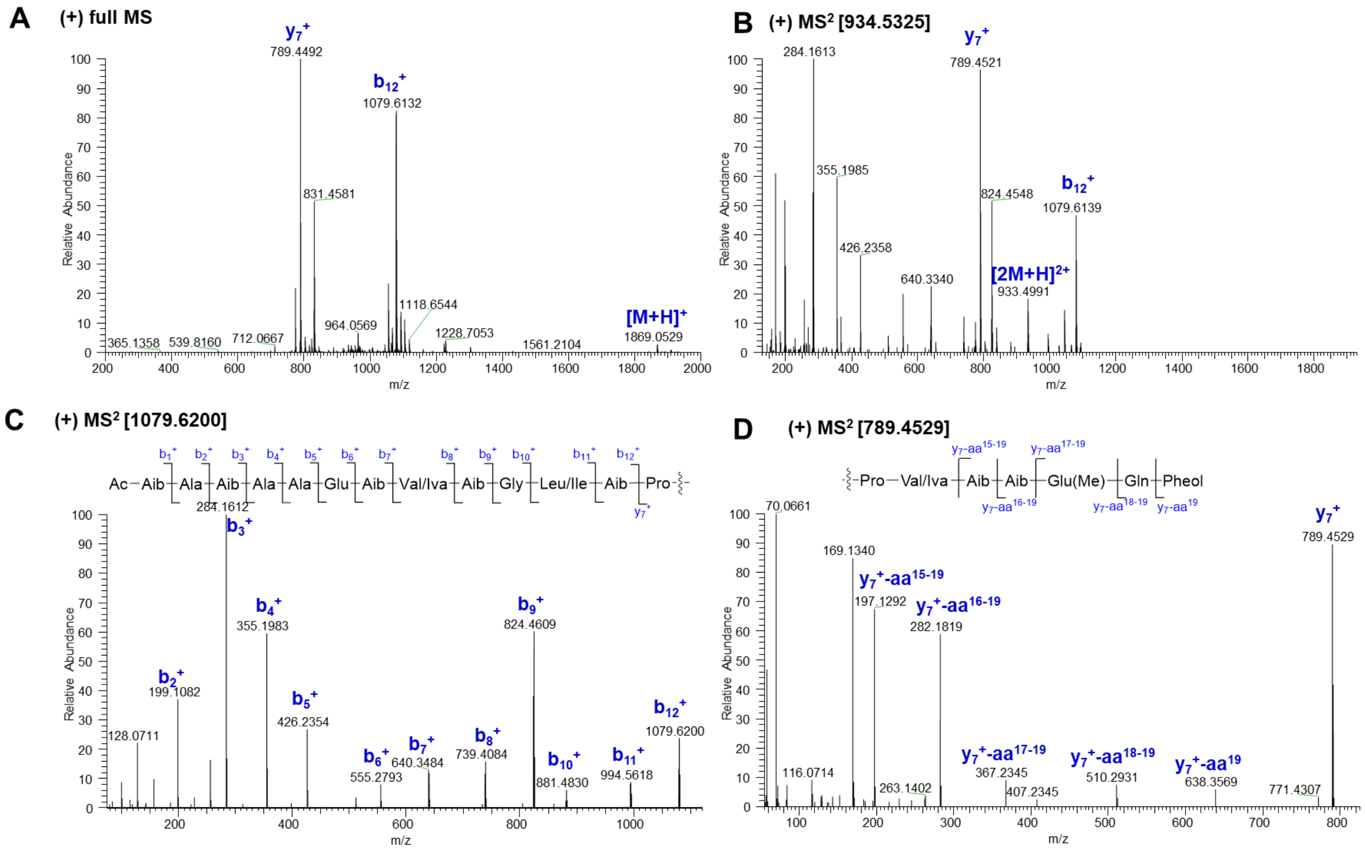

**Fig. S50** Diagnostic fragment ions [*m/z*] of **34** in subfraction 5 (SF5) by ESI-HRMS analysis

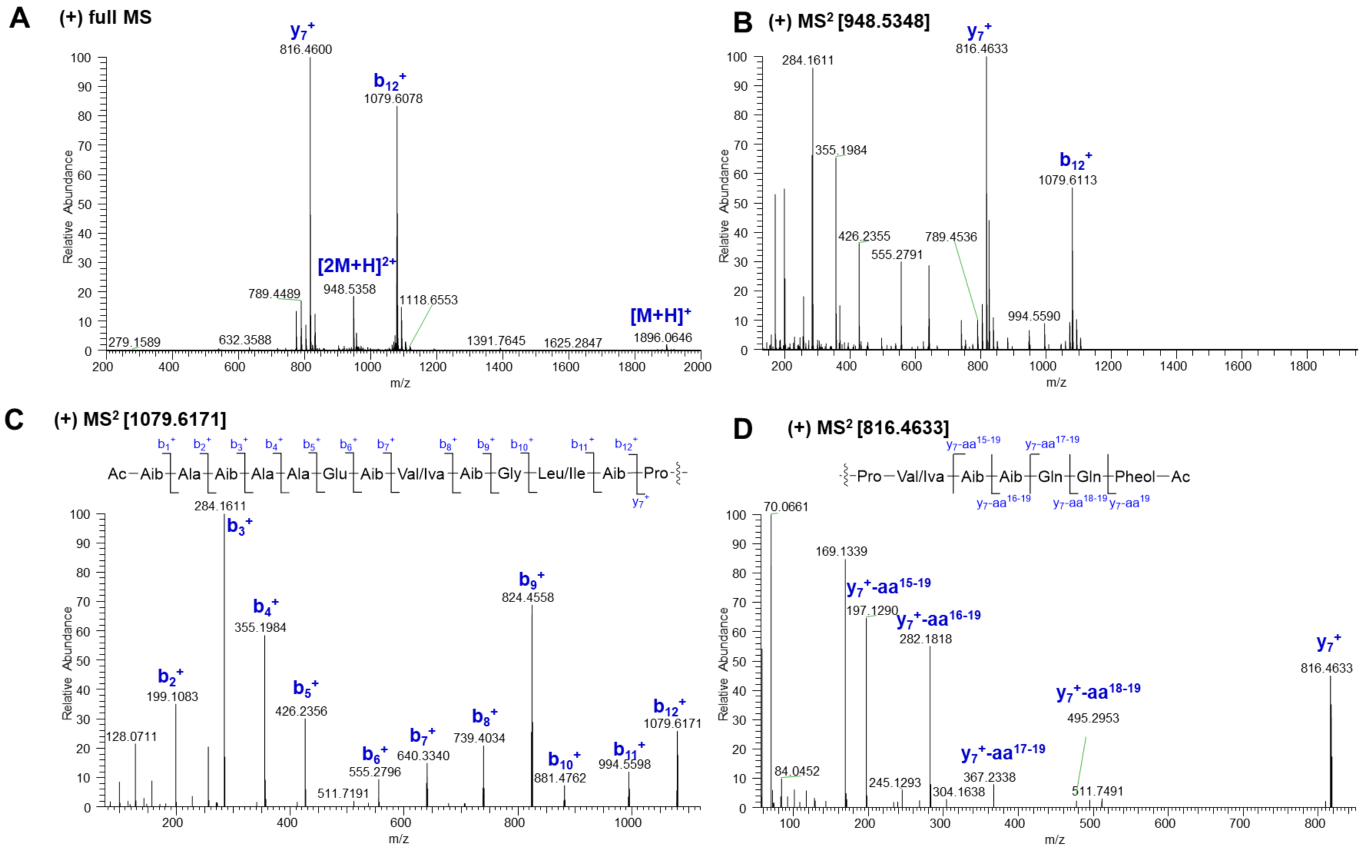

**Fig. S51** Diagnostic fragment ions [*m/z*] of **35** in subfraction 5 (SF5) by ESI-HRMS analysis

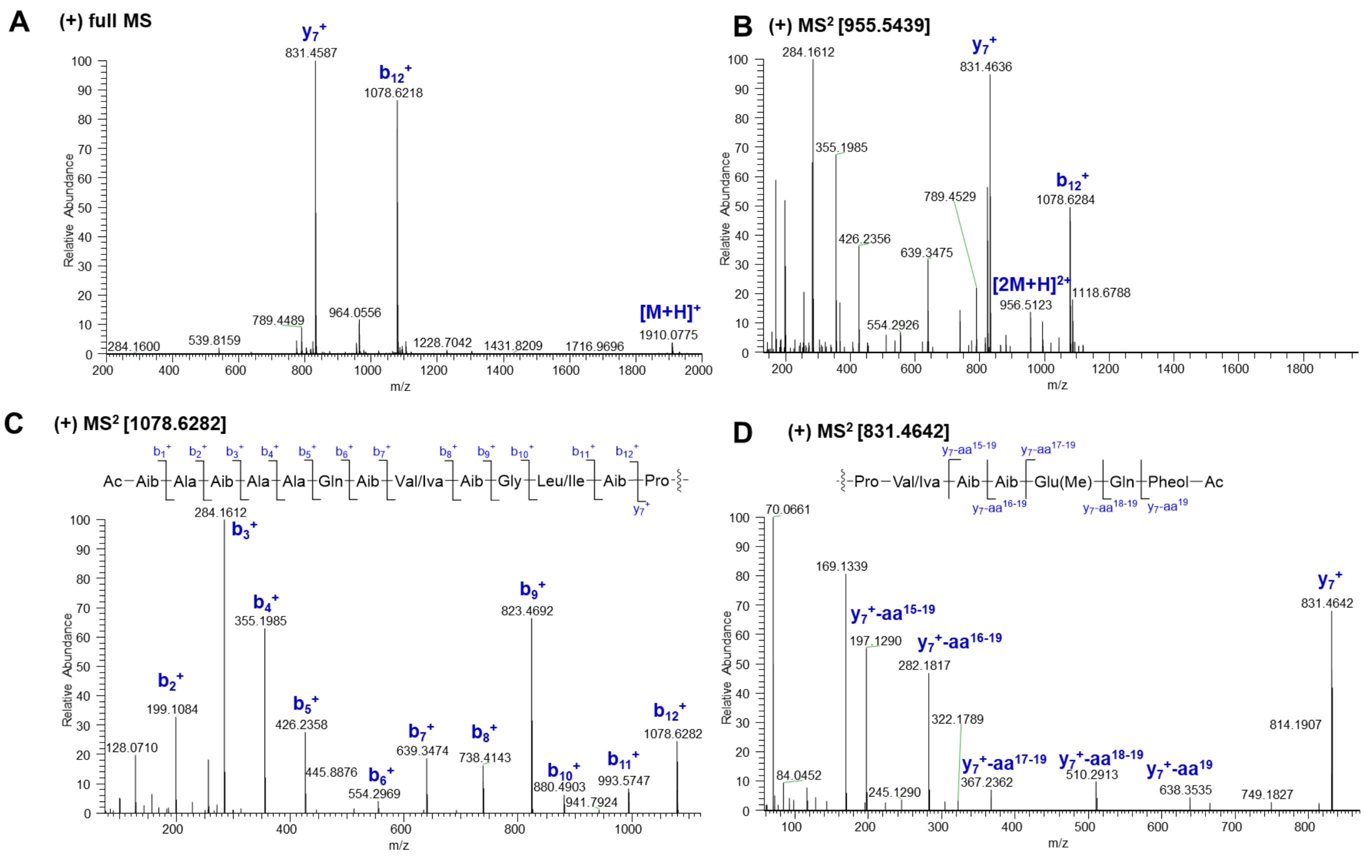

**Fig. S52** Diagnostic fragment ions [*m/z*] of **36** in subfraction 5 (SF5) by ESI-HRMS analysis

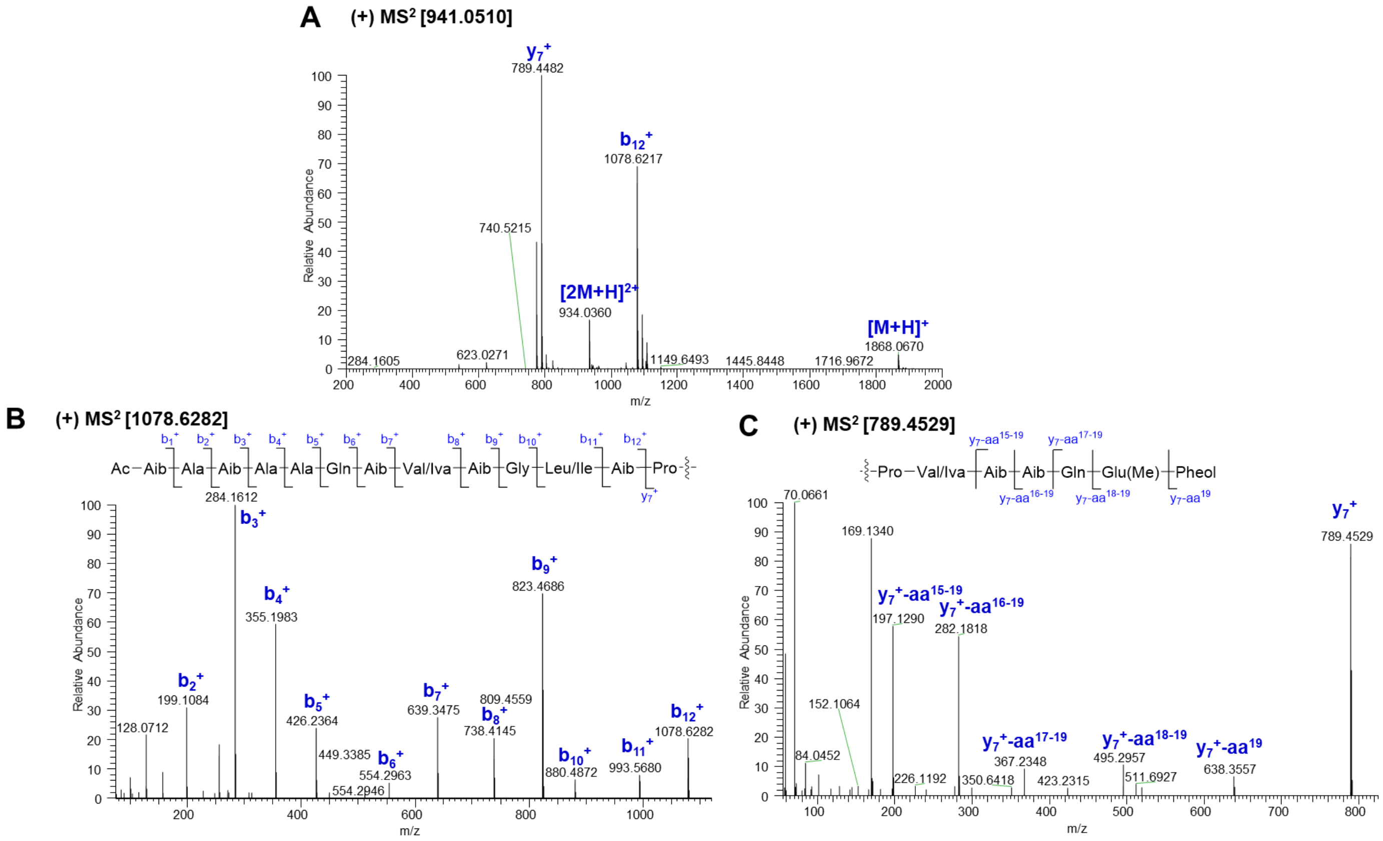

**Fig. S53** Diagnostic fragment ions [*m/z*] of **37** in subfraction 5 (SF5) by ESI-HRMS analysis

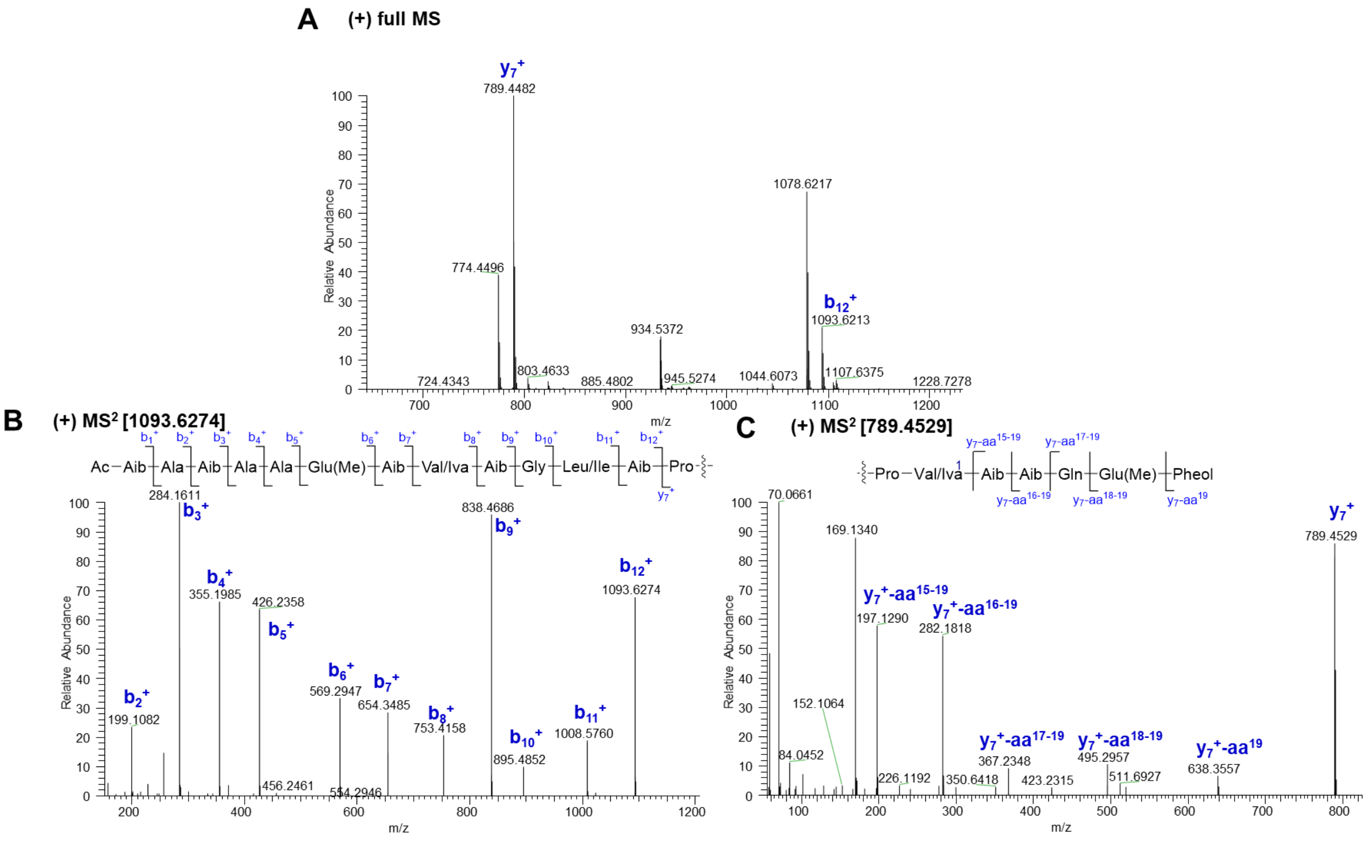

**Fig. S54** Diagnostic fragment ions [*m/z*] of **38** in subfraction 5 (SF5) by ESI-HRMS analysis

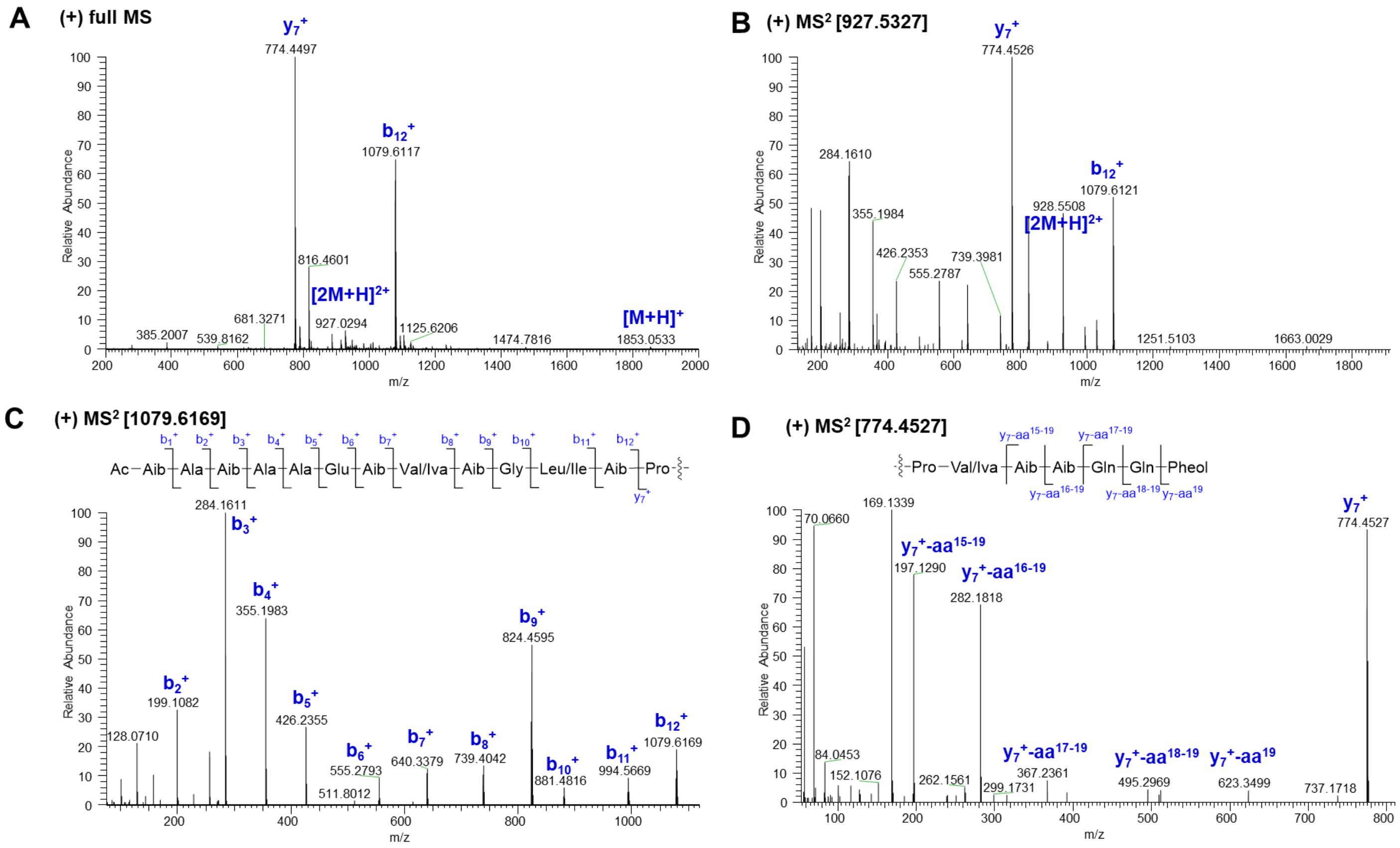

**Fig. S55** Diagnostic fragment ions [*m/z*] of **39** in subfraction 5 (SF5) by ESI-HRMS analysis

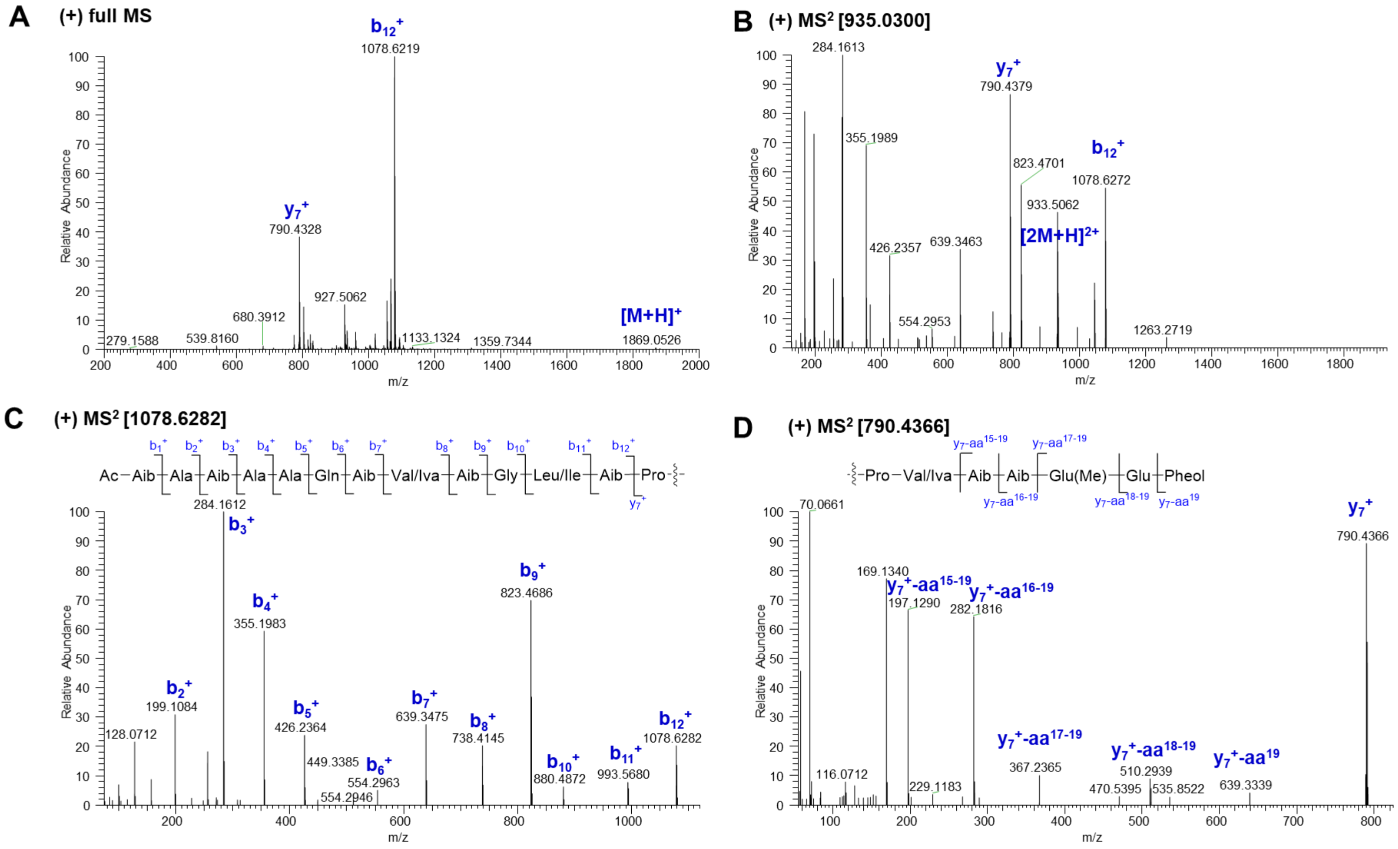

**Fig. S56** Diagnostic fragment ions [*m/z*] of **40** in subfraction 5 (SF5) by ESI-HRMS analysis

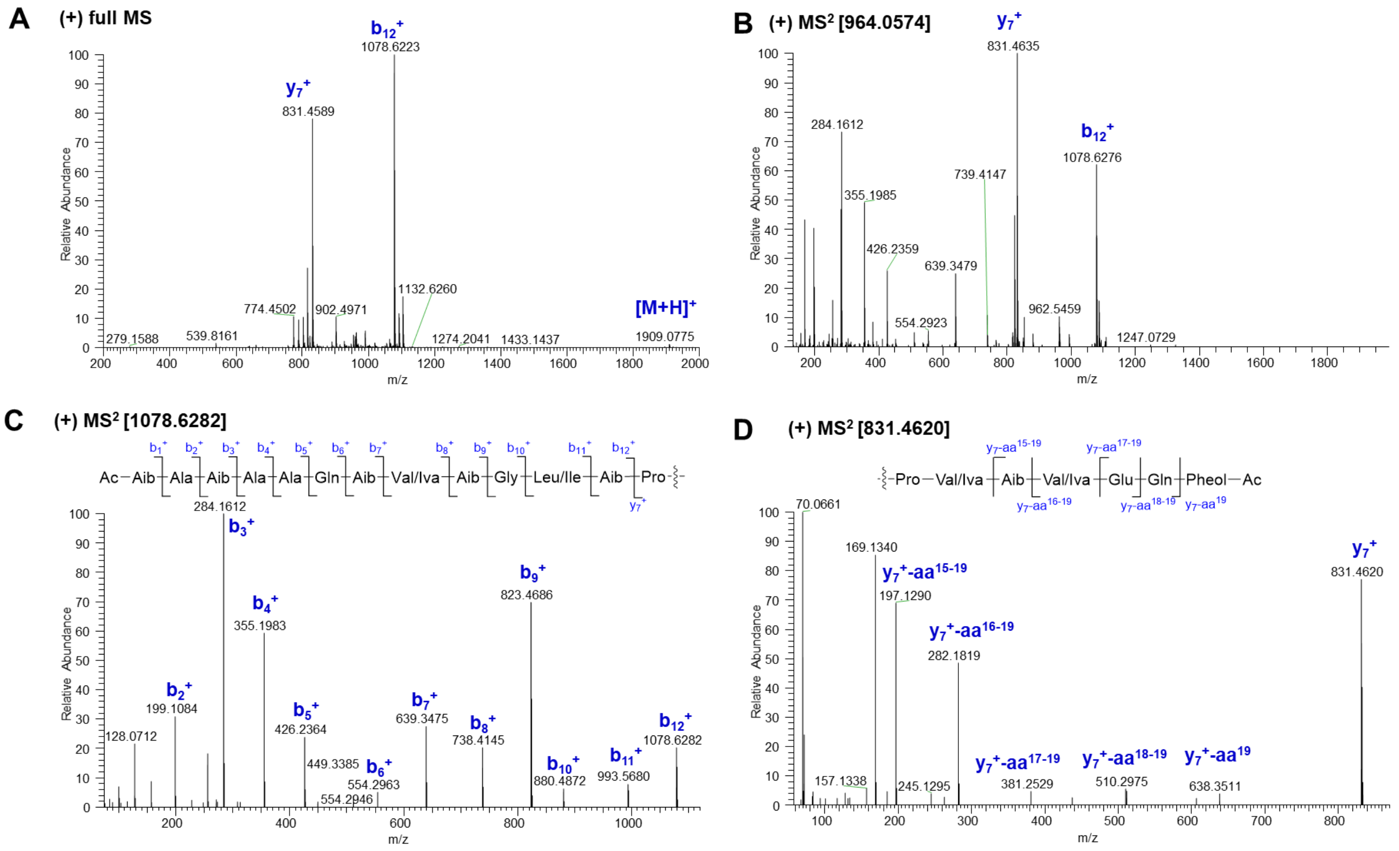

**Fig. S57** Diagnostic fragment ions [*m/z*] of **41** in subfraction 5 (SF5) by ESI-HRMS analysis

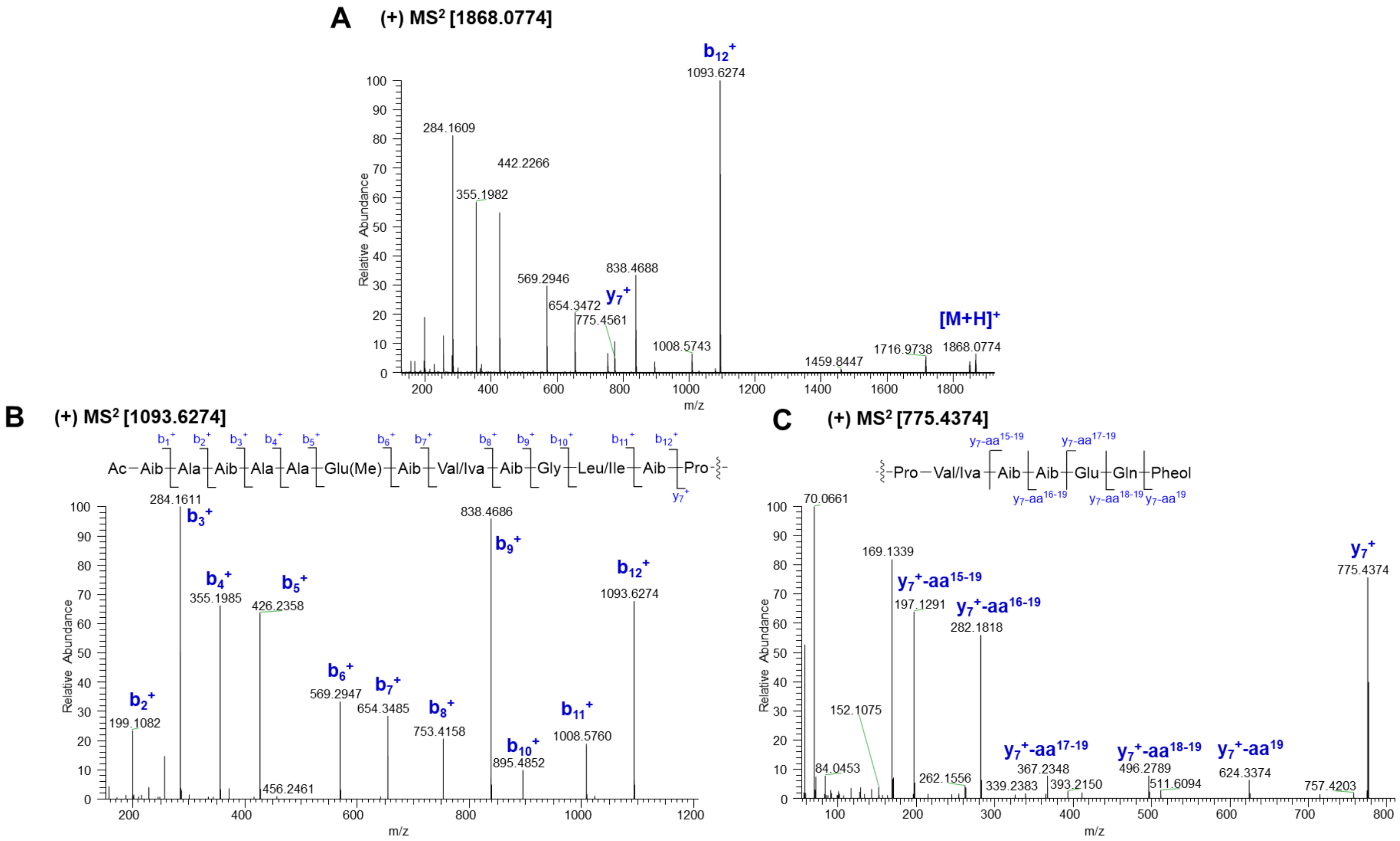

**Fig. S58** Diagnostic fragment ions [*m/z*] of **42** in subfraction 5 (SF5) by ESI-HRMS analysis
